# On the hybrid origin of the C_2_ *Salsola divaricata* agg. (Amaranthaceae) from C_3_ and C_4_ parental lineages

**DOI:** 10.1101/2021.09.23.461503

**Authors:** Delphine T. Tefarikis, Diego F. Morales-Briones, Ya Yang, Gerald Edwards, Gudrun Kadereit

## Abstract

- C_2_ photosynthesis is characterized by recapturing photorespiratory CO_2_ by RuBisCO in Kranz-like cells and is therefore physiologically intermediate between C_3_ and C_4_ photosynthesis. C_2_ can be interpreted as an evolutionary precursor of C_4_ and/or as the result of hybridization between a C_3_ and C_4_ lineage.
- We compared the expression of photosynthetic traits among populations of the *Salsola divaricata* agg. (C_2_) from humid subtropical to arid habitats on the coasts of the Canary Islands and Morocco and subjected them to salt and drought treatments. We screened for enhanced C_4_-like expression of traits related to habitat or treatment. We estimated species trees with a transcriptome dataset of Salsoleae and explored patterns of gene tree discordance. With phylogenetic networks and hybridization analyses we tested for hybrid origin of the *Salsola divaricata* agg.
- We observed distinct independent variation of photosynthetic traits within and among populations and no clear evidence for selection towards C_4_-like trait expression in more stressful habitats or treatments. We found reticulation and gene tree incongruence in Salsoleae supporting a putative hybrid origin of the *Salsola divaricata* agg.
- C_2_ photosynthesis in the *Salsola divaricata* agg. combines traits inherited from its C_3_ and C_4_ parental lineages and seems evolutionarily stable, possibly well adapted to a wide climatic amplitude.

## Introduction

In current models of C_4_ evolution, the C_3_-C_4_ intermediate phenotypes (including C_2_ plants) are interpreted as a transitional evolutionary link between the ancestral C_3_ photosynthesis pathway and the derived C_4_ pathway and showcase the complexity of the numerous structural, genetic, and functional changes necessary to establish a functioning C_4_ pathway (Bräutigam & Gowik, 2016; Schlüter & Weber, 2016). There are about 50 known species with intermediate C_3_-C_4_ traits, some of which do not share a common ancestor with one of the more than 60 independent C_4_ lineages, and some of which represent lineages up to 20 or even 30 million years in their crown age (Sage *et al*., 2018). The fact that we observe these intermediate phenotypes in nature repeatedly implies that they can be evolutionary stable (Lundgren, 2020). Their phenotypic diversity has been classified into four photosynthetic categories of C_3_-C_4_ intermediacy: proto-Kranz, C_2_ type I, C_2_ type II and C_4_-like (Sage *et al*., 2014). C_2_ photosynthesis has been repeatedly interpreted as a crucial stepping stone towards C_4_ photosynthesis (Edwards, 2019). In C_2_ species the photorespiratory enzyme glycine decarboxylase (GDC) is restricted to the bundle sheath or Kranz-like cells and mitochondria are absent or distinctly reduced in the mesophyll cells. This induces a photorespiratory glycine shuttle to the bundle sheath cells where the glycine is then processed by GDC and the photorespiratory CO_2_ is recaptured by RuBisCO (Schulze *et al*., 2016). Bräutigam & Gowik (2016) proposed that once the photorespiratory CO_2_ pump is active, establishing the C_4_ cycle is inevitable. According to this model, a strong selective pressure towards an increase of phosphoenolpyruvate carboxylase (PEPC) activity in C_2_ species should be expected under carbon deficient conditions (Bräutigam & Gowik, 2016).

A possible alternative, but not mutually exclusive hypothesis, is the origin of C_3_-C_4_ intermediates through hybridization (Monson *et al*., 1984). Experimental hybrids of C_3_ and C_4_ species reveal similar phenotypes as naturally occurring C_3_-C_4_ intermediate species and a segregating of photosynthetic traits in F2+ generations (Björkman *et al*., 1969; Holaday *et al*., 1985; Cameron *et al*., 1989; Brown *et al*., 1993; Brown & Bouton, 1993; Oakley *et al*., 2014). Furthermore, conflicting topologies in molecular phylogenetic studies, that includes C_3_-C_4_ intermediate species, indicate possible hybridization events in the evolutionary history of these lineages (reviewed in Kadereit *et al*., 2017). Hybridization at some point in the history of a C_4_ lineage might have perturbed the sequence of events depending on the photosynthetic phenotypes of the parental lineages. To investigate the possibility of a hybrid origin of a C_3_-C_4_ intermediate lineage, detailed phylogenomic studies are needed. For example, evidence of recent and ancient events of reticulate evolution in a phylogenomic study of *Flaveria* revealed a potential major role of hybridization in the evolution of C_4_ photosynthesis in that genus (Morales-Briones and Kadereit, 2022).

Here we conducted a study of a C_2_ species aggregate at the population level and looked at C_4_-adaptive traits to search for empirical evidence for the expected evolutionary shifts towards more C_4_-like traits under stressful conditions according to the model proposed by Bräutigam & Gowik (2016). At the same time, we investigated the origin of this aggregate using a phylogenomic approach. The C_2_ species we chose for our study are *Salsola divaricata* Moq. (Amaranthaceae, subfamily Salicornioideae, tribe Salsoleae; Schüssler *et al*., 2017; Morales-Briones *et al*., 2021) and its close relatives *S. verticillata* Schousb., *S. gymnomaschala* Maire and *S. deschaseauxiana* Litard. & Maire (hereafter called the *S. divaricata* agg.). The distribution area of the *Salsola divaricata* agg. on the Canary Islands and the coasts of Western Morocco, spans a considerable climatic gradient with an arid climate in the east and a mesic mediterranean climate in the west (García-Herrera *et al*., 2001). All four species of the *Salsola divaricata* agg. are salt tolerant, perennial shrubs with terete succulent leaves which are morphologically difficult to distinguish (Padrón Mederos, 2012) and seem mainly geographically defined. They all have a Kranz-like salsoloid leaf anatomy (Voznesenskaya *et al*., 2013; Schüssler *et al*., 2017). *Salsola divaricata* was categorized as a C_2_ type I which is characterized by the photorespiratory CO_2_ pump through GDC being confined to the bundle sheath cells while having C_3_-like activity of C_4_ enzymes (e.g., PEPC and NADP-ME). Their CO_2_ compensation points around 30 µmol mol^-1^ are intermediate to C_3_ and C_4_ species (Schüssler et al., 2017). The *Salsola divaricata* agg. belongs to Salsoleae, a tribe rich in C_4_ species, but also with several C_3_ species and several C_3_-C_4_ intermediate species. The *Salsola divaricata* agg. forms a monophyletic, well-supported subclade nested in a clade of C_3_-C_4_ intermediate and C_3_ species with uncertain placement (Schüssler *et al*., 2017).

The two aims of this study were, to test whether the lineage is stable in its C_2_ state or is shifting towards stronger C_4_ physiology under more stressful climatic conditions on the eastern Canary Island and Morocco, and to test for a hybrid origin of the *Salsola divaricata* agg. To achieve these aims we studied C_4_ adaptive traits at the population level in the 26 populations of the *S. divaricata* agg. distributed from La Gomera in the west to Morocco in the east and searched for more C_4_-like phenotypes in the more arid eastern parts of the distribution area. We also analyzed 991 loci from a transcriptome data set comprising C_3_, C_4_ and C_2_ species of the Salsoleae and explored incongruences to determine if a hybridization event is plausible in the lineage of the *S. divaricata* agg.

## Materials and Methods

### Plant Material

Seeds and vouchers of 26 populations of the *S. divaricata* agg. were collected in 2013 and 2014 in the Canary Islands and Morocco (**Supporting Information Table S1**; **Fig. 1**) and grown in a greenhouse at the Botanical Garden Mainz with multiple individuals per population. Most individuals of a population share a mother plant, when this is not the case different mother plants are indicated with roman numbers in the individual names (LS No.). Plants were grown in custom mixed soil in standardized clay pots. Day/night cycles of 14h/10h were artificially maintained with natural light and supplementary light providing a light intensity of 200–400 µmol m^−2^ s^−1^. The minimum temperature at night was 18 °C and daytime temperatures ranged between 25 to 35 °C in summer and 20 to 25 °C in winter. All plants were watered once a week in the winter and twice a week in the summer. The individuals included in the salt treatment were watered once a week and every two weeks with 2% saltwater (20 g NaCl per liter; 34.3 mM) from June 2015 to April 2021. Individuals in the dry treatment were only watered every two weeks from June 2015 to March 2017. The analyses of the photosynthesis traits began in 2016. **Supporting Information Table S1** lists the populations included in this study as well as the collection and voucher information.

**Fig. 1:**
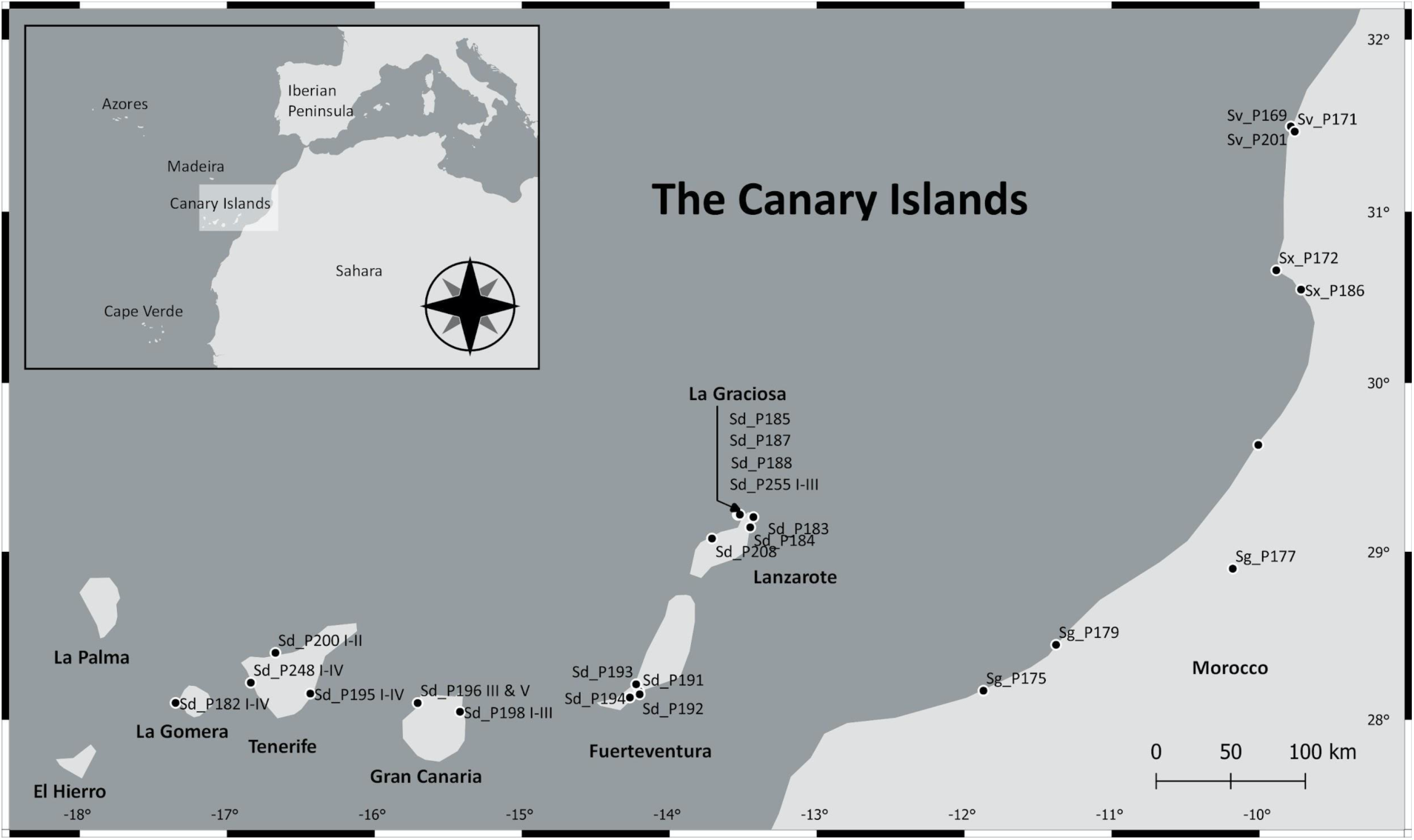
Map of the collection locations of seeds and vouchers for the *S. divaricata* agg. (See also **Supporting Information Table S1** for voucher information). In total we analyzed 26 populations of the *S. divaricata* agg. on the Canary Islands and Morocco. Species name abbreviation and population number: Sd = *Salsola divaricata*, Sg = *S. gymnomaschala*, Sv = *S. verticillata*, Sx = *S. deschaseauxiana;* P*xxx* = population number. Note: Morphological studies on the species complex (Brullo, 1982; Fennane and Ibn Tattou, 1998) consider *S. verticillata* and *S. deschaseauxiana* as conspecific, while *S. gymnomaschala* seems more like *S. divaricata* s.s.

### Carbon isotope measurements

In C_3_ plants Rubisco discriminates against assimilating atmospheric ^13^CO_2_. C_4_ plants exhibit much lower discrimination against ^13^CO_2_ through its conversion to bicarbonate and assimilation by PEPC. Thus, calculation of δ^13^C values can be used to detect C_4_ cycle activity where more negative values indicate higher discrimination against fixing ^13^CO_2_ (values for C_3_ plants are -22 to -32 °% and C_4_ plants -9 to -16 ^**o**^%, Sage 2016). We measured samples of dried leaves collected in the field from the same mother plants from which seeds were collected. We compared their δ^13^C values with samples taken from the greenhouse cultivated offspring. We also determined and compared δ^13^C values between and within greenhouse cultivated populations of *S. divaricata*. To determine carbon isotope composition, a standard procedure with PDB (Pee Dee Belemnite) limestone as the carbon isotope standard was used (Bender *et al*., 1973). Leaf samples were dried in silica gel, and then 1–2 mg was placed in a tin capsule and combusted in an EuroVector elemental analyser (EuroVector, Italy). After separating the resulting N_2_ and CO_2_ gasses by gas chromatography, they were fed into the IsoPrime™ isotope ratio mass spectrometer (IRMS; GV Instruments Ltd. (Micromass Ltd., United Kingdom) for determination of ^13^C/^12^C ratios (R). δ^13^C values were determined according to δ^13^C(‰)=1000(R_sample_/R_standard_)^-1^, where R is ^13^C/^12^C. Measurements were taken at Washington State University and at the Geology Department of the University of Mainz.

### CO_2_ compensation point measurements

The CO_2_ compensation point (Γ), the concentration where there is no net uptake of CO_2_ in the light, is a trait that is often used to identify C_4_ species due to their relatively low values (usually less than 4 µmol CO_2_ mol^-1^) compared to C_3_ species. We measured individuals of several populations per island of all species in the *S. divaricata* agg. (n = 53; 1 to 3 technical replicates for each individual). The C_3_ *S. webbii* and C_4_ *S. oppositifolia* were measured for comparison. CO_2_ compensation points were determined using the portable gas exchange measurement system GFS-3000 (Heinz Walz GmbH, Germany) with the standard measuring head 3010-S equipped with a standard leaf area cuvette and a LED-Array (for details of the cuvette settings see **Supporting Information Material & Methods S1**). As a general calculation parameter in GFS-Win, we used leaf area which was measured using Easy Leaf Area v1.02 or Image J 1.49d (Wayne Rasband, National Institutes of Health, USA) followed by a calculation according to the approximately cylindrical leaf shape. To determine the CO_2_ compensation point, net rates of CO_2_ assimilation were plotted against intercellular CO_2_ concentration.

### PEPC activity measurements

Phosphoenolpyruvate carboxylase (PEPC) is a key enzyme in the C_4_ pathway. It fixes atmospheric CO_2_ after its conversion to HCO_3_^-^ by carbonic anhydrase into a four-carbon compound which gave this photosynthesis pathway its name (Reyna-Llorens & Hibberd, 2017). C_4_ plants show high levels of PEPC activity compared to C_3_ plants and an increased activity is seen as an important stage in evolving from an intermediate to a full C_4_ photosynthesis pathway (Sage *et al*., 2012). PEPC activity measurements were conducted using the Tecan infinite M1000 (Tecan Trading AG, Männedorf, Switzerland) and a Greiner UV-Star 96-well plate (Sigma-Aldrich Chemie GmbH, Munich, Germany). For details of sample preparation see **Supporting Information Material & Methods S1**.

The assay of PEPC was initiated by adding leaf extract and 3.9 mM phosphoenolpyruvate (PEP). The chlorophyll content of the leaf extract was measured according to Wintermans and De Mots (1965) in 96% ethanol. PEPC activity (*a*) was calculated based on the linear part of the progress curve where *a*= *SU*[µmol ml^-1^ min^-1^]*V*_assay_[ml]/(1000*V*_e*nzym*e_[ml]*c*_*Chla*+*b*_[mg ml^-1^]).

SU is the substrate turnover rate, *V*_assay_ is the total assay volume, *V*_e*nzym*e_ is the volume of the leaf extract and *c*_*Chla*+*b*_ is the concentration of chlorophyll in the leaf extract. Statistical analyses were conducted in R v3.1.2 (R Core Team, 2014).

### Transcriptome processing and nuclear phylogenetic analyses

To evaluate the potential hybrid origin of the *S. divaricata* agg., we assembled a dataset comprising 13 transcriptomes representing 12 C_3_, C_4_ and C_2_ species of Salsoleae. The data set included ten publicly available transcriptomes and three newly sequenced species on an Illumina HiSeq 2500 platform at the University of Minnesota Genomics Center (paired-end 125 bp). For RNA extraction protocol and library preparation refer to Morales-Briones *et al*. (2021). Additionally, we sampled three closely related Amaranthaceae species as outgroups. **Table 1** lists all samples with their photosynthesis type, ploidy if known and SRA accession number.

**Table 1:**
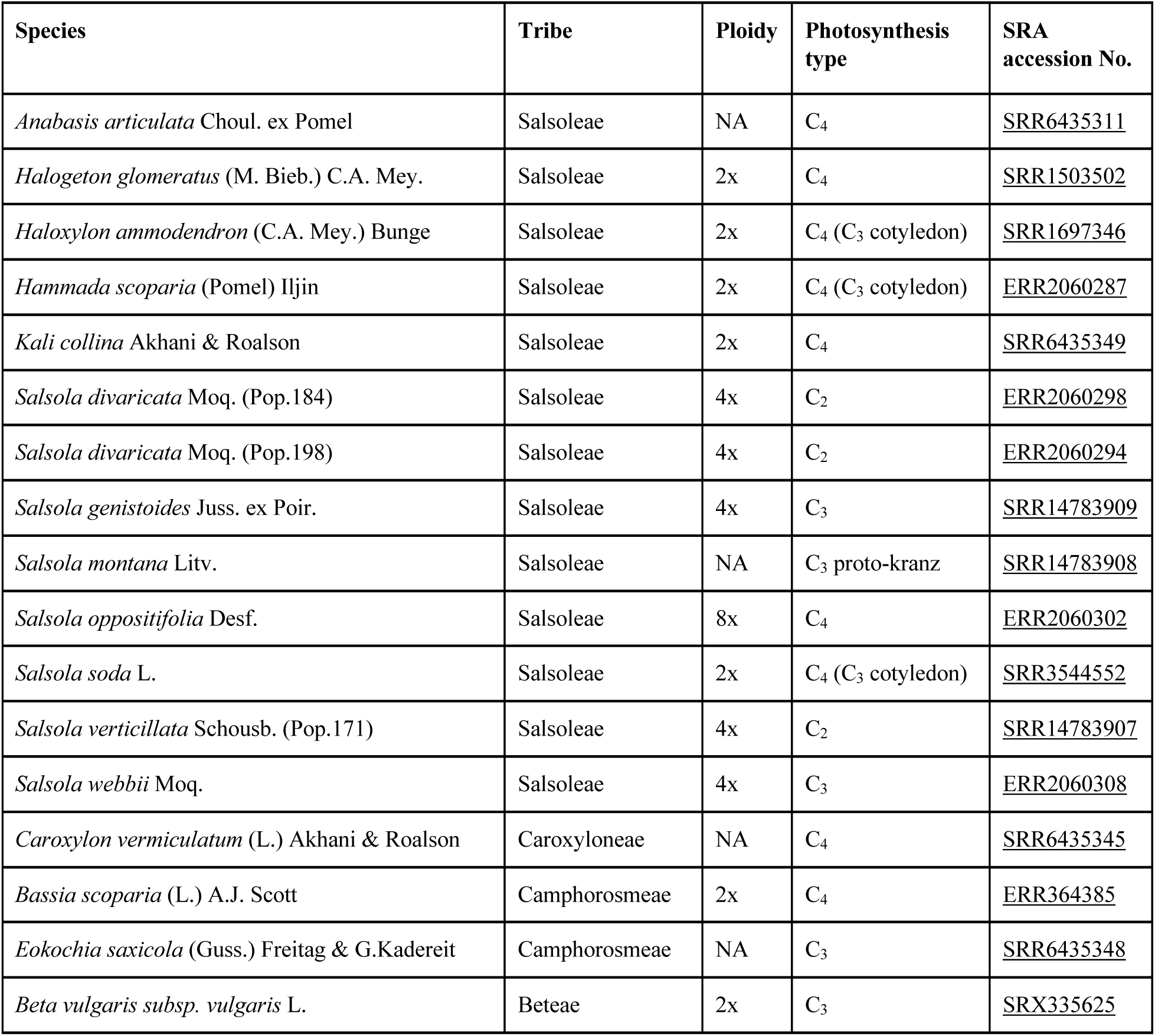
List of the 17 samples in the transcriptome phylogeny; with photosynthesis type, ploidy (Kew C value database 2018) and NCBI Sequence Read Archive (SRA) accession number.

Raw read processing, transcriptome assembly, low-quality and chimeric transcript removal, transcript clustering into putative genes, translation, final coding sequences (CDS) redundancy assessment, and homology and orthology inference were carried out following Morales-Briones et al. (2021; for details see **Supporting Information Material & Methods S1**). Briefly, an all-by-all BLASTN search was performed, and putative homologs groups were clustered using MCL (van Dongen 2000). Homolog trees were built using RAxML (Stamatakis 2014) and spurious tips were removed with TreeShrink (Mai and Mirarab 2018). Monophyletic and paraphyletic tips that belonged to the same taxon were removed, leaving one tip with the most aligned characters to obtain final homologs. Orthology inference was carried out following the ‘monophyletic outgroup’ approach from Yang and Smith (2014), keeping only ortholog groups with all 17 taxa present.

We applied concatenation and coalescent-based methods for phylogenetic reconstruction. For the concatenation approach, we prepared a supermatrix by keeping only ortholog alignments with at least 300 bp. We estimated a maximum likelihood (ML) tree with RAxML v8.2.11 (Stamatakis, 2014) using a partition by gene scheme. Clade support was assessed with 100 rapid bootstrap (BS) replicates. To estimate a species tree that is statistically consistent with the multi-species coalescent (MSC), individual gene trees were used to infer a species tree using ASTRAL-III v5.6.3 (Zhang *et al*., 2018) using local posterior probabilities (LPP; Sayyari & Mirarab, 2016) to assess clade support. To examine nuclear gene tree discordance, we first calculated the Internode Certainty All score (ICA; Salichos *et al*., 2014). Also, we calculated the number of concordant and conflicting bipartitions on each node of the species trees. We calculated both the ICA scores and conflict analyses with Phyparts (Smith *et al*., 2015). Additionally, to distinguish strong conflict from weakly supported branches, we evaluated tree conflict and branch support with Quartet Sampling (QS; Pease *et al*., 2018) using 1,000 replicates (for details see **Supporting Information Material & Methods S1**).

### Assessment of hybridization

To detect possible hybridization, first we inferred species networks under a maximum pseudo-likelihood (Yu & Nakhleh, 2015) approach using PhyloNet v3.6.9. (Than *et al*., 2008). To estimate the best number of hybridizations and test whether the species network fits our gene trees better than a bifurcating tree, we performed model selection using the bias-corrected Akaike information criterion (Sugiura, 1978) and the Bayesian information criterion (Schwarz, 1978). We also used HyDe (Blischak *et al*., 2018) to estimate the amount of admixture (γ) in putative hybrid lineages. We tested all triples combinations and significance was assessed with a Bonferroni correction (for details see **Supporting Information Material & Methods S1**).

### Plastome assembly and phylogenetic analysis

To investigate phylogenetic signals from plastid sequences, *de novo* assemblies were carried out with the Fast-Plast v1.2.6 pipeline (https://github.com/mrmckain/Fast-Plast) using the filtered organelle reads obtained from the transcriptome raw read processing. A ML tree was inferred with IQ-TREE v1.6.1 (Nguyen *et al*., 2015) using the automated model selection (Kalyaanamoorthy *et al*., 2017) and 200 standard non-parametric BS replicates for branch support (for details see **Supporting Information Material & Methods S1**). Additionally, we used QS with 1,000 replicates, to detect potential plastome conflict in the backbone as seen in other groups of Amaranthaceae s.l. (Morales-Briones *et al*., 2021).

### Analysis of photosynthetic gene trees

We inferred trees for 50 genes that are important in photosynthesis to determine if the genes in *Salsola divaricata* agg. show a tendency to group with either a C_3_ or a C_4_ species. We downloaded coding and protein sequences from *Arabidopsis thaliana* as baits from the TAIR website (https://www.arabidopsis.org/tools/bulk/sequences/index.jsp). These loci were determined based on Lauterbach *et al*. (2017a, b) and included two RuBisCO subunit genes, four genes coding for proteins of the glyoxylate cycle, 17 genes encoding photorespiratory proteins, and 25 genes coding for C_4_-associated proteins (Lauterbach *et al*., 2017b) as well as two C_4_-associated transcription factors (Lauterbach *et al*., 2017a). Gene annotation and pathway assignment can be found in **Supporting Information Table S3**. We examined whether the *S. divaricata* agg. was sister to the C_3_ species *S. montana* or the C_4_ clade. When there were several gene copies or weak bootstrap support, we additionally noted whether species of the *S. divaricata* agg. were in a clade with C_3_ or C_4_ species (for details see **Supporting Information Material & Methods S1**).

## Results

### Carbon isotope measurements

*Salsola divaricata* plants growing in the wild on the Canary Islands generally had higher carbon isotope (δ^13^C) values (−21 to -28; mean -24.44, n = 30) than those grown from seeds under greenhouse conditions (−22 to -34; mean -27.77, n = 165). Overall, the δ^13^C values of the *S. divaricata* agg. showed a distinct range but not a single individual (stressed or not stressed) came close to a C_4_-like δ^13^C value of > -16 (**Fig. 2**). The control and drought treatment showed no significant difference of δ^13^C values among populations (Kruskal-Wallis chi-squared = 5.1081 and 8.61, df = 5, n = 165 and 36, p-value = 0.40 and 0.13, respectively). In the salt treatment there was a significant difference of δ^13^C values among populations (Kruskal-Wallis chi-squared = 15.47, df = 5, n = 36, p-value = 0.009), which was due to significantly higher δ^13^C values in salt treated individuals of populations 191 compared to 182 and 194. We tested for differences of carbon isotope values between treatments within populations of the *Salsola divaricata* agg. and found a significant difference for populations 182, 191 and 198 (**Fig. 3**). However, there was no significant difference among the three treatments for populations 184, 188 and 194 (**Fig. 3**). After grouping δ^13^C values according to island, significant differences were found among islands (ANOVA: df = 6, F value = 4.08, p-value = 0.001) due to differences between La Gomera and the mainland (SW Morocco), as well as between Lanzarote, Tenerife, and the mainland, respectively. All other differences were not statistically significant. We found that the populations of *S. verticillata, S. deschaseauxiana* and *S. gymnomaschala* on the Moroccan mainland often had higher δ^13^C values than the populations of *S. divaricata* on the Canary Islands (the non-stressed control groups that are regularly watered, **Supporting Information Fig. S1**). However, due to very different sample sizes further statistical tests including all populations were inconclusive.

**Fig. 2:**
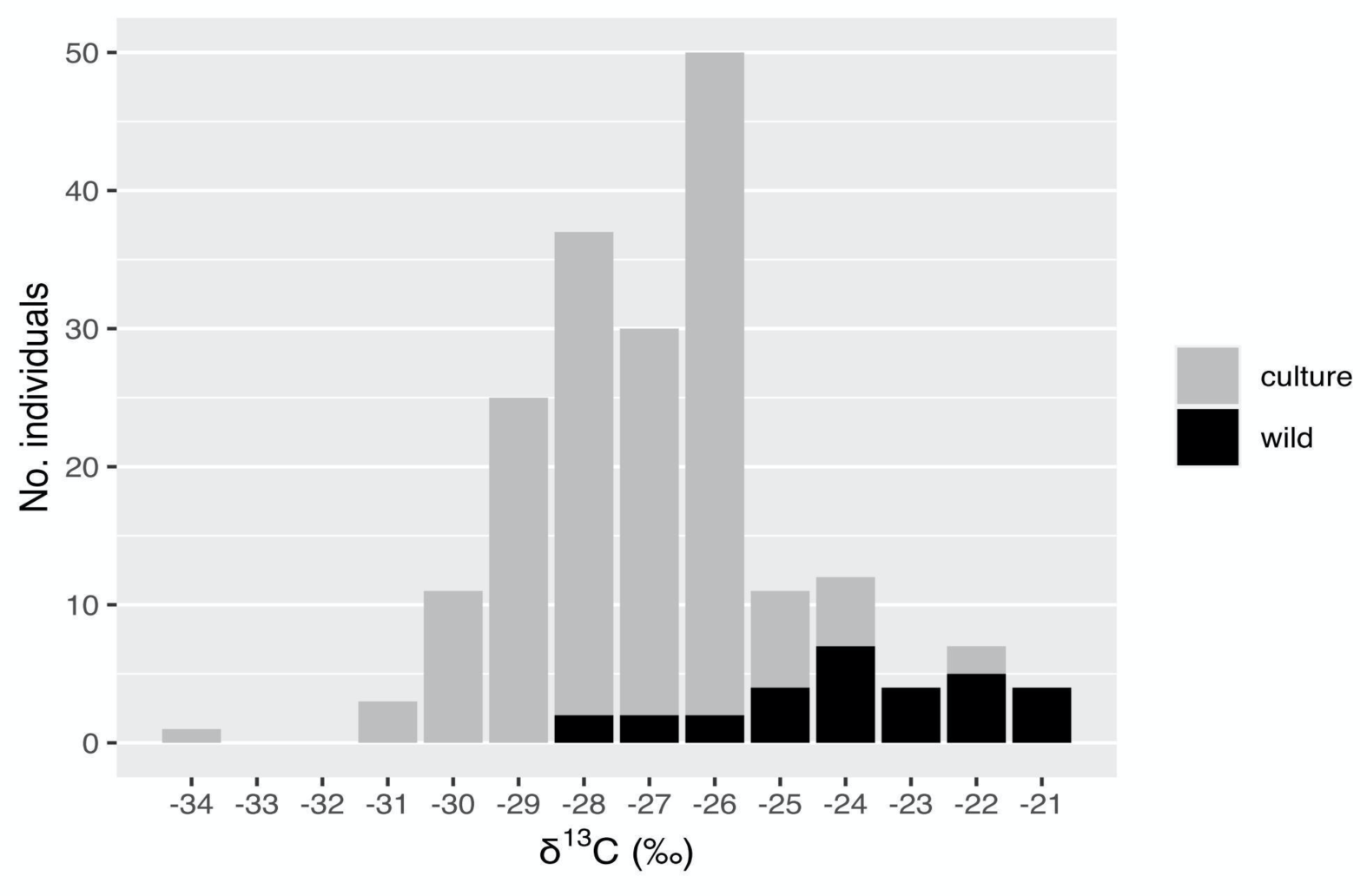
Distribution of δ^13^C (‰) of 195 samples of the *Salsola divaricata* agg., of these 30 were collected from plants growing in the wild and 165 from plants in cultivation at the Botanical Garden Mainz. This includes 36 individuals in dry treatment and 36 in salt treatment, all others are in the control group. See **Table S1** for information on populations included.

**Fig. 3:**
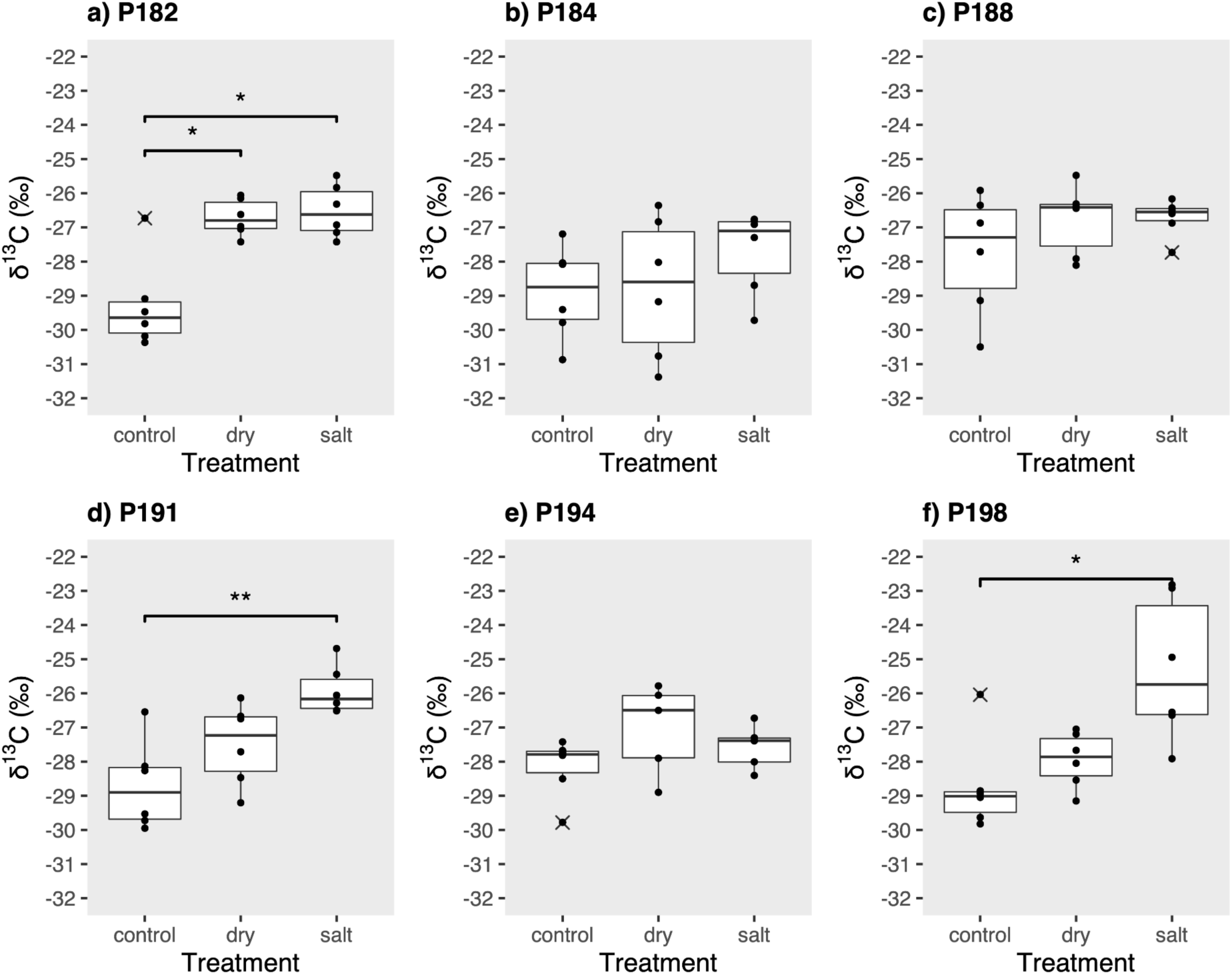
Comparison of carbon isotope values (δ^13^C [‰]) between treatments within populations (P) of the *Salsola divaricata* agg. Significant differences were found for populations 182 from La Gomera (a; Kruskal-Wallis chi-squared = 7.94, df = 2, n = 18, p-value = 0.02), 191 from Fuerteventura (d; Kruskal-Wallis chi-squared = 10.84, df = 2, n = 18, p-value = 0.004), and 198 from Gran Canaria (f; Kruskal-Wallis chi-squared = 8.84, df = 2, n = 18, p-value = 0.01). No significant differences were found between the three treatments for 184 from Lanzarote (b; Kruskal-Wallis chi-squared = 2.33, df = 2, n = 18, p-value = 0.31), 188 from La Graciosa (c; Kruskal-Wallis chi-squared = 0.78, df = 2, n = 18, p-value = 0.68), and 194 from Fuerteventura (e; Kruskal-Wallis chi-squared = 2.27, df = 2, n = 18, p-value = 0.32).

Overall, these results showed variation in δ^13^C values among populations associated with source population and growth conditions; but all were well within the range typical of C2 and C_3_ species. The significantly higher δ^13^C values of salt treated individuals indicated that salinity rather than drought affects the δ^13^C in *Salsola divaricata* agg. and could be responsible for the occasional observation of relatively high δ^13^C values of our samples collected in the wild close to the shoreline.

### CO_2_ compensation point measurements

The CO_2_ compensation points (Γ) of the four species of the *Salsola divaricata* agg. lained between those of the C_3_ (*Salsola webbii*) and C_4_ species (*Salsola oppositifolia*) as expected (between 20 and 30 µmol CO_2_ mol^-1^, **Fig. 4a**). The mean values of *S. deschaseauxiana* and *S. verticillata* were slightly lower than those of *S. divaricata* and *S. gymnomaschala*. When looking at individual measurements we again observed a broad range from 17 to 40 µmol CO_2_ mol^-1^ with one outlier at 9 µmol CO_2_ mol^-1^ (**Supporting Information Fig. S2**). Populations growing on the more arid Islands such as Lanzarote and Fuerteventura or in Morocco did not show lower CO_2_ compensation points. Due to varying values within each population (**Supporting Information Fig. S2**) and the relatively small sample sizes (one to two individuals per population) we did not conduct further statistical analysis.

**Fig. 4:**
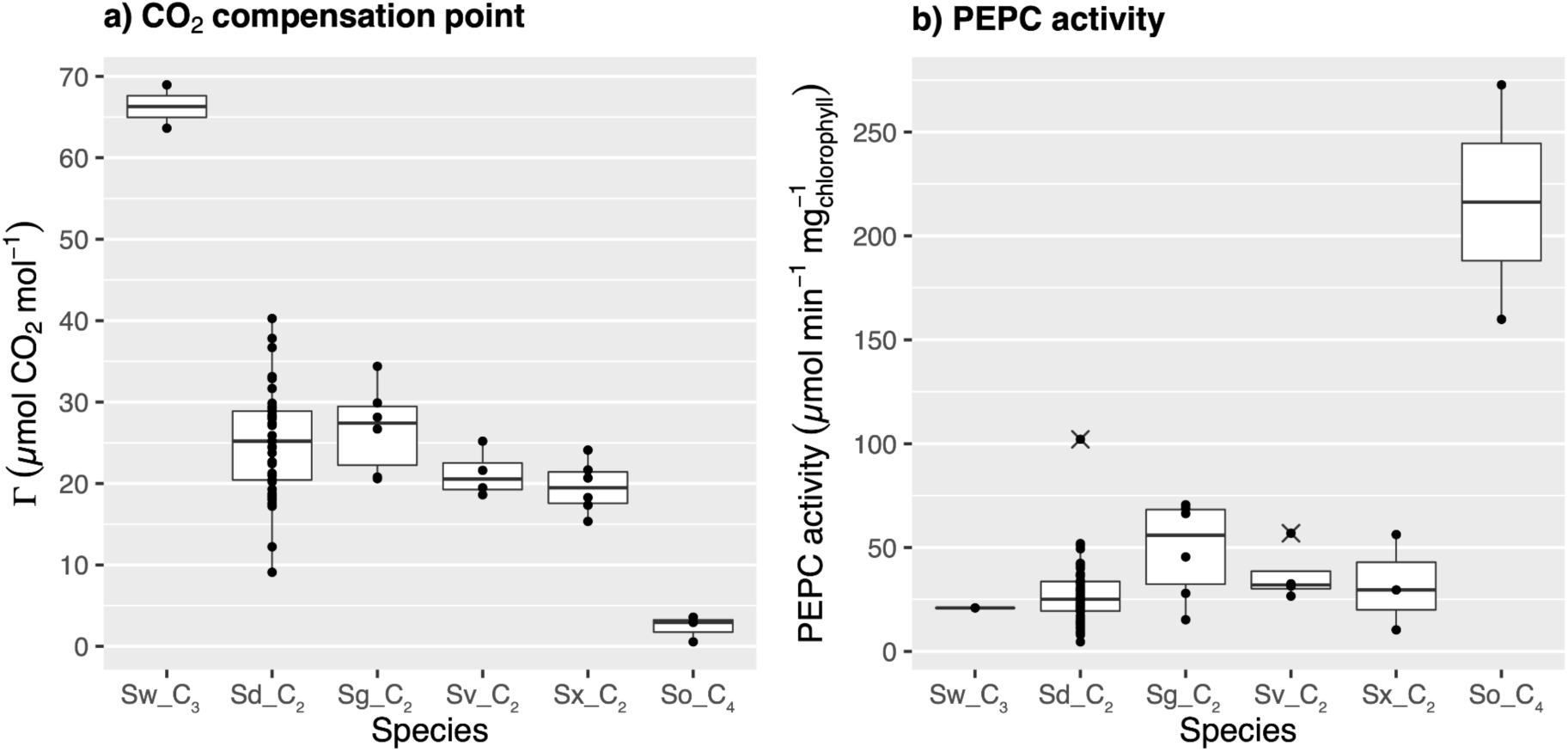
**a)** CO_2_ compensation points (Γ in µmol CO_2_ mol^-1^) of the separate species of the C_2_ *Salsola divaricata* agg. (Sd_C_2_ = *Salsola divaricata*, mean 25.14 µmol CO_2_ mol^-1^, n = 21; Sg_C_2_ = *S. gymnomaschala*, mean 26.75 µmol CO_2_ mol^-1^, n = 4; Sv_C_2_ = *S. verticillata*, mean 22.21 µmol CO_2_ mol^-1^, n = 4; Sx_C_2_ = *S. deschaseauxiana*, mean 19.56 µmol CO_2_ mol^-1^, n = 4) compared to the values of C_3_ species *Salsola webbii* (Sw_C_3_, 66.3 µmol CO_2_ mol^-1^, n = 2) and C_4_ species *Salsola oppositifolia* (So_C_4_, 2.35 µmol CO_2_ mol^-1^, n = 3). We measured 1–3 technical replicates per individual. **b)** PEPC activity [µmol min^-1^ mg_chlorophyll_^-1^] of the separate species of the C_2_ *Salsola divaricata* agg. (*Salsola divaricata*, mean activity 33 µmol min^-1^ mg_Chlorophyll_^-1^, n = 82; *S. gymnomaschala*, mean 46.8 µmol min^-1^ mg_Chlorophyll_^-1^, n = 10; *S. verticillata*, mean 37.4 µmol min^-1^ mg_Chlorophyll_^-1^, n = 7; *S. deschaseauxiana*, mean 27.6 µmol min^-1^ mg_Chlorophyll_^-1^, n = 6) compared to C_3_ *S. webbii* (mean 30 µmol min^-1^ mg_Chlorophyll_^-1^, n = 1) and C_4_ *S. oppositifolia* (mean 216 µmol min^-1^ mg_Chlorophyll_^-1^, n = 2).

### PEPC activity measurements

First, we compared the PEPC activity of the separate species of the *Salsola divaricata* agg. to the values of the closely related C_3_ species *Salsola webbii* and C_4_ species *Salsola oppositifolia*. C_3_ and C_4_ values were as expected clearly distinguishable (**Fig. 4b**). The values of the C_2_ species were closer to those of the C_3_ species and much lower than the C_4_ species (**Fig. 4b**). We used a Kruskal-Wallis test to determine if statistically significant differences exist between C_2_ species (Kruskal-Wallis chi-squared = 9.94, df = 3, p-value = 0.019). This significant difference was observed due to the values of *S. divaricata* on the Canary Islands having a median of 30.05 µmol min^-1^ mg_Chlorophyll_^-1^ and *S. gymnomaschala* in Morocco having a median of 48.32 µmol min^-1^ mg_Chlorophyll_^-1^. Due to the high level of variation within species we took a closer look at 14 populations of *S. divaricata* (n = 5–7 per population) and observed a significant difference between them (Kruskal-Wallis chi-squared = 34.64, df = 13, p-value = 0.001, see also **Fig. 5a**). When grouped according to island, we found a significant difference (Kruskal-Wallis chi-squared = 22.199, df = 5, p-value = 0.0005) which is due to the difference in values between populations on Gran Canaria and Lanzarote, as well as populations on La Graciosa and Fuerteventura and on La Graciosa and Gran Canaria (**Fig. 5b**). We did not find a significant difference between treatments in the greenhouse in *S. divaricata* (Kruskal-Wallis chi-squared = 1.21, df = 2, p-value = 0.55). Again, we observe a high variation of PEPC activity values within populations and islands.

**Fig. 5:**
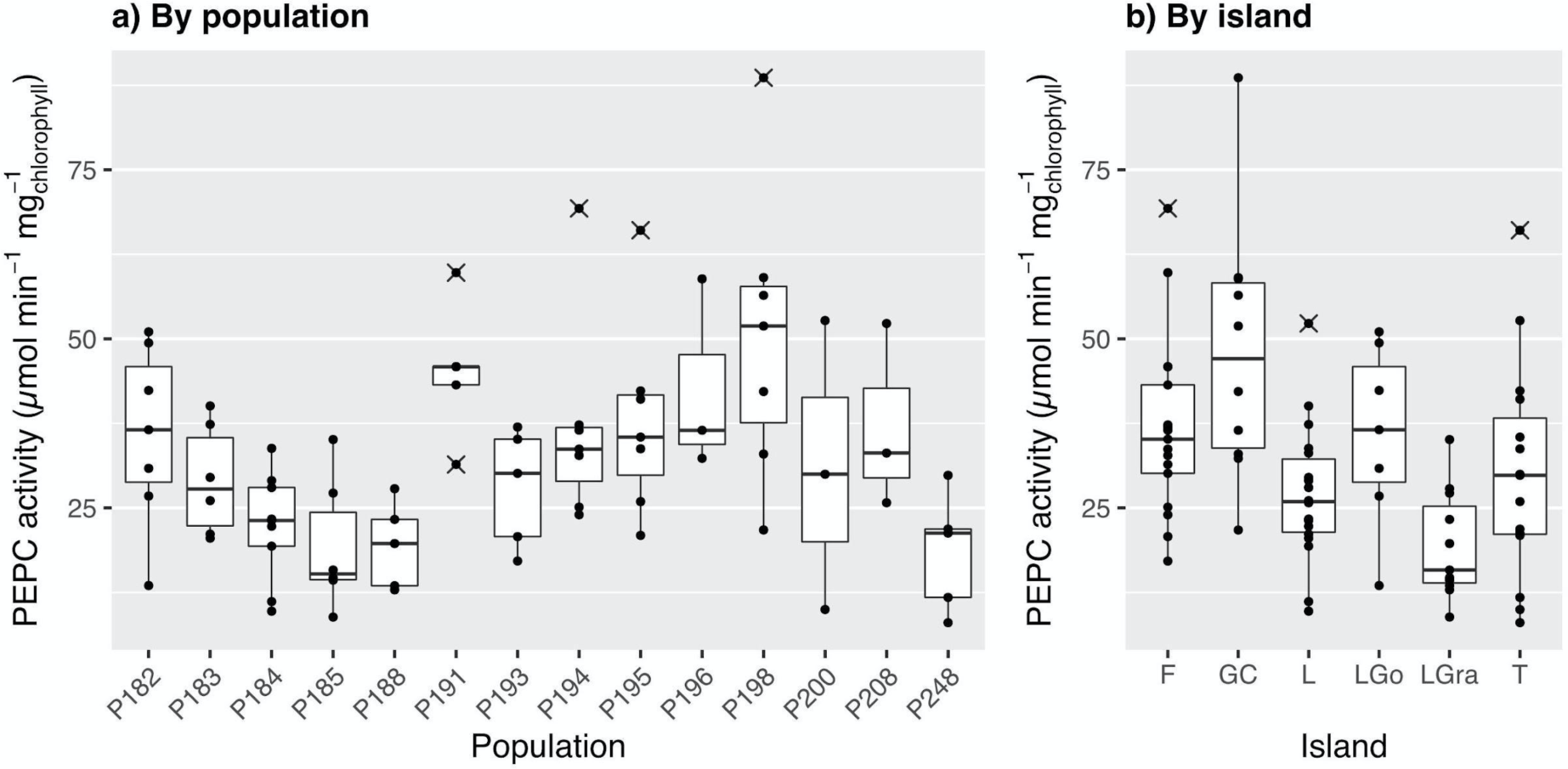
Boxplots of PEPC activity [µmol min^-1^ mg_chlorophyll_^-1^] of *Salsola divaricata* Moq. **a)** Values grouped by population (P). **b)** Values grouped by island; F = Fuerteventura, GC = Gran Canaria, L = Lanzarote, LGo = La Gomera, LGra = La Graciosa, T = Tenerife.

Linear regression to test for correlation between PEPC activity and carbon isotope (δ^13^C) values (Multiple R^2^ = 0.008, F-statistic = 0.35, df = 44, p-value = 0.56), between PEPC activity and CO_2_ compensation point (Γ) (Multiple R^2^ = 0.04, F-statistic = 1.19, df = 29, p-value = 0.28), and between δ^13^C values and Γ (Multiple R^2^ = 0.017, F-statistic = 0.46, df = 27, p-value = 0.5) recovered no significant correlation among the three.

### Phylogenomic analyses

The raw reads for the newly generated transcriptomes are available from the NCBI Sequence Read Archive (**Table 1**). The 991 final orthologs had alignments of lengths between 315 and 6621 bp. The concatenated matrix comprised 1,427,449 aligned columns with overall matrix occupancy of 86% (see **Supporting Information Table S2** per details).

The ASTRAL species tree and the concatenated RAxML tree (**Supporting Information Fig. S3**) had similar topologies with most nodes receiving the maximum support (BS = 100%; LPP = 1). The only difference between the two topologies is that *Anabasis articulata* and *Salsola soda* were sisters with maximum support in the RAxML tree but formed a grade with lower support (LPP = 0.74) in the ASTRAL tree. Concordance analyses (**Supporting Information Fig. S3** and **S4**) showed high support for the monophyly of Salsoleae and the *S. divaricata* agg., while signaling conflict for the remaining taxa. The placement of *S. genistoides* had low support with only 261 (of 794) concordant gene trees (ICA = -0.45) and a QS score (0.22/0/0.98) suggesting an alternative topology. Monophyly of the *S. divaricata* agg. clade composed of *S. divaricata* + *S. verticillata* had strong support with 808 (of 913) concordant gene trees (ICA = 0.71) and maximum QS support (1/–/1; i.e., all sampled quartets supported that branch). However, the two samples of *S. divaricata* showed signals of alternative topologies with only 332 (of 900) gene trees supporting their sister relationship (ICA = 0.56) and QS counter support (−0.25/0/1). The sister relationship of *S. divaricata* agg. and the C4 remaining Salsoleae had low support with only 178 (of 705) concordant gene trees (ICA = 0.29) and low QS support (0.085/0/1 [ASTRAL]; (0.11/0/1) [RAxML]) suggesting an alternative topology. The placement of the C4 *Anabasis articulata* and *S. soda*, which varied between RAxML and ASTRAL trees, also had low gene tree and QS support with signals of alternative topologies. The clade composed of *S. oppositifolia, Halogeton, Hammada*, and *Holoxylon* was supported only by 280 (of 692l ICA = 0.24), but had maximum QS support, while within the clade all relationships had low gene tree and QS support signaling alternative topologies.

The final plastid alignment had 80,903 characters with a matrix occupancy of 70%. The plastid tree largely differed from the nuclear topologies and had most nodes with maximum support (BS = 100; **Fig. 6** and **S5**). *Caroxylon vermiculatum, Kali collina*, and *S. genistoides* form a well-supported clade (BS = 100; QS = 0.56/0/1) that is sister to all Salsoleae. *S. webbii* and *Anabasis articulata* form a grade with strong support (BS =100; QS = 0.6/0/0.99 [*S. webii*]; QS = 1/–/1 [*A. articulata*]). The remaining species form an unsupported clade (BS = 0.47; QS = -0.0096/0.88/0.68) composed of two subclades. The first subclade (BS = 93; QS = 0.19/0.91/0.78) is composed of *S. verticillata* + *S. montana* (BS = 100; QS = 0.61/0/1) and the two samples of *S. divaricata* (BS = 100; QS = 1/–/0.77). The second subclade (BS = 100; QS = 0.59/0/0.95) has *Salsola soda* as the sister the clade (BS =100; QS = 1/–/1) composed of *S. oppositifolia, Halogeton, Hammada*, and *Haloxylon*.

**Fig. 6:**
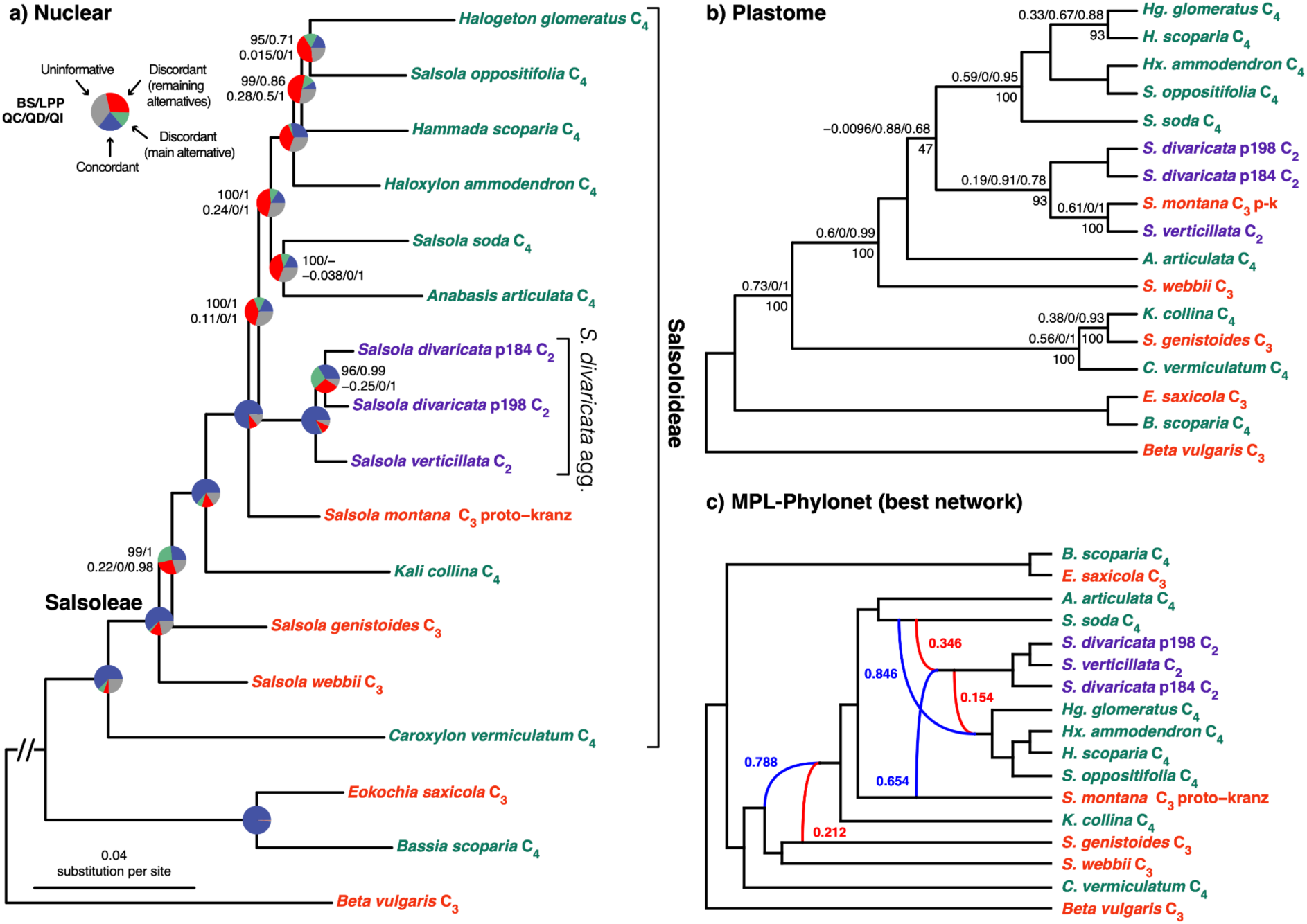
a) Maximum likelihood phylogeny of Salsoloideae inferred from the RAxML analysis of the concatenated 991-nuclear gene supermatrix from the ‘monophyletic outgroup’ (MO) orthologs. Bootstrap support and local posterior probabilities (BS/LPP) are shown above branches. Nodes with full support (BS = 100/LPP = 1) are not shown. Em dashes (—) denote alternative topology compared to the ASTRAL tree (**Supporting Information Fig. S3**). Quartet Sampling (QS) scores are shown below branches. QS score: Quartet concordance (QC)/Quartet differential (QC)/Quartet informativeness QI). Full QS support (1/–/1) not shown. Pie charts represent the proportion of gene trees that support that clade (blue), the proportion that support the main alternative bifurcation (green), the proportion that support the remaining alternatives (red), and the proportion (conflict or support) that have < 50% bootstrap support (gray). Branch lengths are proportional to the number of substitutions per site (scale bar on the bottom). b) Cladogram of Salsoloideae inferred from IQ-TREE analysis of concatenated complete and partial plastomes. QS scores and BS support are shown above and below branches, respectively. See Supporting Information Fig. S5 for phylogram. c) Best maximum pseudo-likelihood species network inferred with PhyloNet. Red and blue curved branches indicate the minor and major edges, respectively of hybrid nodes. Numbers next to curved branches indicate inheritance probabilities for each hybrid node.

### Assessment of hybridization

Model selection preferred a network as a better model than any bifurcating tree (**Table 2**). A network with three reticulation events was the best model overall (PhyloNet-4-hybridization search; AICc = 37875.93; BIC = 38054.302; **Fig. 6c**). The next model was a significantly worse model (also a network with three reticulation events [PhyloNet-3-hybridization search] with ΔAICc and ΔBIC of 753.208. The *S. divaricata* agg. clade was shown as the product of a hybridization event between *S. soda* (C_4_) (inheritance probability p_i_ = 0.346) and *S. montana* (C_3_) (p_i_ = 0.654). Then the C_4_ clade *S. oppositifolia* + *Halogeton* + *Hammada* + *Haloxylon* was the product of reticulation between *S. soda* (C_4_) (p_i_ = 0.846) and the stem lineage of the C2 *S. divaricata* agg. (p_i_ = 0.154). These two clades were also recovered as hybrids in the networks with one, two and three reticulation events with similar parental lineages and inheritance probabilities (**Supporting Information Fig S6**). The third event was a deeper reticulation within Salsoleae which showed that most species of the group (excluding *Caroxylon*) were a product of an ancient hybridization event between *S. genistoides* (p_i_ = 0.212) and the sister lineage of *S. genistoides + S. webbii* (p_i_ = 0.788). The HyDe analysis resulted in 1,680 hybridization tests, of which 298 triples were significant (**Supporting Information Table S3, Fig. S7**). The significant triples showed that 12 out of 13 individuals of Salsoleae are involved in multiple hybridization events, which is mostly in agreement with the nested reticulation events detected with Phylonet. All sampled accessions of the *S. divaricata* aggr. showed similar hybridization patterns (**Fig. S7;** i.e. same potential parental lineages and admixture parameter [γ]) consistent with a single ancient hybrid origin of *S. divaricata* agg. as shown in the Phylonet analyses. Overall, the admixture parameter (γ) ranged from 0.013 to 0.986 (average 0.458). These results indicate multiple, complex hybridizations within Salsoleae. Additionally, Hyde also identified *Eokochia saxicola* as a hybrid (**Supporting Information Table S3, Fig. S7**).

**Table 2:** Model selection between bifurcating trees and networks. The best model (lowest score) is the network with three hybridization events from PhyloNet (best MPL run allowing four hybridization events). k = No. branch lengths + hybridization probabilities [(2n-3)+(2h)]; h = number of hybridization events. All trees and networks had 17 taxa (n). Networks and bifurcating trees log likelihood values were inferred with 991 loci.

| tree/network | ln L (log likelihood) | k | $h^1$ | AICc | $\Delta AICc$ | BIC | $\Delta BIC$ |
| --- | --- | --- | --- | --- | --- | --- | --- |
| ASTRAL-nuclear | -19754.54543 | 31 | N/A | 39573.130 | 1697.190 | 39722.951 | 1668.649 |
| RAxML-nuclear | -19741.89558 | 31 | N/A | 39547.830 | 1671.891 | 39697.651 | 1643.349 |
| IQTREE-platome | -22836.08408 | 31 | N/A | 45736.207 | 7860.268 | 45886.028 | 7831.726 |
| PhyloNet. 1-hybridization search (network 1) | -19451.79848 | 31 | 1 | 38967.636 | 1091.696 | 39117.457 | 1063.155 |
| PhyloNet. 1-hybridization search (network 2) | -19605.00833 | 33 | 1 | 39278.323 | 1402.383 | 39437.674 | 1383.372 |
| PhyloNet. 1-hybridization search (network 3) | -19624.56237 | 33 | 1 | 39317.431 | 1441.491 | 39476.782 | 1422.480 |
| PhyloNet. 2-hybridization search | -19435.04197 | 35 | 2 | 38942.674 | 1066.734 | 39111.539 | 1057.237 |
| PhyloNet. 3-hybridization search | -19276.12872 | 37 | 3 | 38629.147 | 753.208 | 38807.510 | 753.208 |
| PhyloNet. 4-hybridization search | -18899.52481 | 37 | 3 | 37875.940 | 0.000 | 38054.302 | 0.000 |
| PhyloNet. 5-hybridization search | -19283.48635 | 41 | 5 | 38652.512 | 38652.512 | 38849.820 | 38849.820 |

<sup>1</sup>The number of reticulation events in the best networks of a search is not necessarily the same number of reticulation events allowed in the search.

### Analysis of individual genes involved in photosynthesis or photorespiration

The 50 genes ranged from 7 to 65 sequences due to variation in the number of accessions and gene copies (**Supporting Information Table S4**). Summarizing all genes, *Salsola divaricata* agg. appeared 22 times as sister to C_3_ *S. montana* and 26 times as sister to C_4_ species (**Fig 7**; **Supporting Information Table S4**). *Salsola divaricata* agg. was not detected in three genes (Isocitrate lyase, RuBisCO subunits). When multiple copies per gene were recovered for *S. divaricata* agg, they were often non-monophyletic, but instead separated in clades with copies from the C_3_ and/or C_4_ species. In these 23 gene trees, sequences of the *S. divaricata* agg. appeared 21 times with C_3_ and 26 times with C_4_ species. Summarizing the results for genes typical for C_4_ pathways, the *S. divaricata* agg. was 27 times sister to and/or nested in a C_3_ clade and 32 times sister to and/or nested in a C_4_ clade). Taking into consideration bootstrap support (BS ≥ 70), the numbers reduced to 18 times sister to and/or nested in a C_3_ clade and 23 times sister to and/or nested in a C_4_ clade for C_4_ genes. For the two C_4_-associated transcription factors, *S. divaricata* agg. was sister to the C_4_ clade for SHORTROOT with strong support (BS = 86). The sister relationship to the C_4_ clade was moderately supported for BEL1-like homeodomain 7 (BS = 57 in clade with C_4_). For the genes assigned to the Glyoxylate cycle *S. divaricata* agg. had multiple copies that were grouped with either C_3_ or C_4_ with high support (BS > 80). The gene for Isocitrate lyase was missing in the *S. divaricata* agg. and *S. montana. Salsola divaricata* agg. is also missing in the alignments for the RuBisCO subunits. For photorespiration-associated genes many species, including those of *S. divaricata* agg. had multiple copies, even though during the transcriptome processing putative genes were inferred to remove isoforms and assembly artifacts. Copies of *S. divaricata* agg. were nested within C_3_ clades with strong support (9 times, BS > 70). Affinity to the C_4_ clade is well supported in 13 of 19 cases, in which they appear as sister or nested within the clade. Just like the *S. divaricata* agg., the C_4_ clade is also not monophyletic in most cases. Clade assignment for all 50 genes is listed in **Supporting Information Table S4** and the individual trees are shown in the **Supporting Information Fig S8**.

**Fig. 7.**
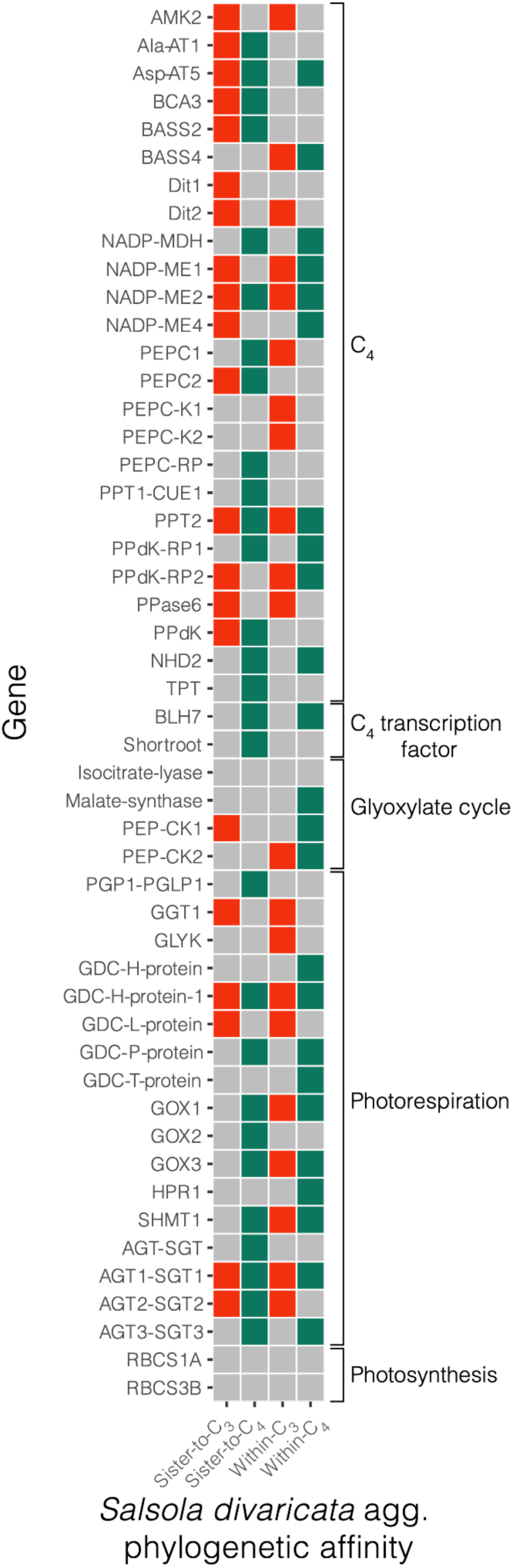
Summary of the phylogenetic affinity of the *Salsola divaricata* agg. in analyzed photosynthetic gene trees. Red and green colored boxes represent phylogenetic affinity with C_3_ or C_4_ species in the specified gene, respectively. Gray boxes denote no affinity. Gene with only grey boxes denote that the gene was not detected in *Salsola divaricata* agg. transcriptomes we sequenced. For detailed phylogenetic affinity see **Supporting Information Table S4 and Fig S8**.

## Discussion

Hybridization experiments with parental lineages exhibiting different photosynthetic types have shown the transferability of C_4_ traits (Apel et al., 1988). For a few examples, it has been suggested that this might also occur in nature (Dunning et al., 2017) which could mean an immediate adaptive advantage of the hybrid lineage in more stressful environments. A phylogenomic study of *Flaveria* revealed a major role of hybridization and transgressive segregation in C_4_ evolution (Morales-Briones and Kadereit, 2022). In this study we aimed to infer the phylogenetic origin of C_3_-C_4_ intermediacy of the *Salsola divaricata* agg. and possible phenotypic variation across a steep climate gradient. *Salsola divaricata* is currently classified as a C_2_ Type I according to Sage *et al*. (2014). According to the model of C_4_ evolution an increase of PEPC activity should be expected under more stressful (more carbon deficient) conditions in the more arid areas (Lanzarote, Fuerteventura, Morocco) with a transition to the C_2_ Type II or even C_4_-like intermediate type, resulting in a reduction of photorespiration (Sage *et al*., 2014; Bräuigan & Gowik, 2026). Alternatively, C_2_ photosynthesis in *Salsola divaricata* agg. may represent a stable condition across mesic and arid growing conditions as presently observed in this study.

### Carbon isotope measurements depict a stress reaction in the *S. divaricata* agg.; but without occurrence of a more C_4_-like photosynthesis in populations growing in drier environments

Although samples collected in the wild had slightly higher δ^13^C values than those sampled in our greenhouse (stressed or control), there is still a clear difference to typical values of C_4_ species. None of the samples showed a δ^13^C value typical for a C_4_-like or a C_4_ plant, which would be around -9 to -16 ‰ (Sage, 2016); instead, all samples had values falling within the range typical for C_3_ species, which would be around -22 to -32 ‰ (Sage, 2016). In C_3_ plants the δ^13^C values of leaves depend on several factors including stomatal conductance and capacity to capture CO_2_ in photosynthesis, which is dependent on light intensity, growth temperature, photochemistry, and capacity for carbon assimilation. The plants growing in the greenhouse are under less stress for potential water loss than field conditions due to lower light intensity, and temperature being more moderate than some field conditions. The results suggest a greater stomatal limitation on photosynthesis in the field grown plants, which results in a low intercellular CO_2_ concentration. This will change the discrimination against the heavier C isotope since RuBisCO will capture more ^13^CO_2_ resulting in slightly higher δ^13^C values (von Caemmerer, 1992).

We observed variation in the δ^13^C values. Populations on the Moroccan mainland often have higher values than the populations on the Canary Islands; but there are many exceptions in populations on the more arid Islands of Fuerteventura and La Graciosa as well as on the more humid Islands of Gran Canaria, Tenerife and La Gomera. There does not seem to be a clear pattern correlated with the general climatic conditions on the Islands. Therefore, the observed variation is likely due to plasticity or local, microclimatic differences. In any case, there are no indications of populations exhibiting C_4_-like photosynthesis based on the δ^13^C values.

*Salsola divaricata* plants from all populations grown for several years under salt stress in the greenhouse showed clear morphological differences in comparison to the control plants. They had more succulent leaves, did not grow to the same height as their counterparts in the control group and most individuals in the salt group flowered earlier. In three of the six tested populations, La Gomera (182), Fuerteventura (191), and Gran Canaria (198), this long-term salt stress was reflected in a significantly higher δ^13^C value. The three other populations from Lanzarote, La Graciosa and Fuerteventura had slightly, but not significantly higher, values for the salt treatment compared to the control group. This suggests *Salsola divaricata* can adapt to salt stress by increasing water storage and decreasing stomatal conductance and it explains some of the δ^13^C value variation found in the wild as salinity certainly varies among sites.

### C_3_-C_4_ intermediate CO_2_ compensation points for *S. divaricata* agg. supports a stable C_2_-state

The CO_2_ compensation points of the *S. divaricata* agg. varied but they are well within the range of other C_2_ species (Sage *et al*., 2014 and ref. therein) and they were significantly lower than the values of C_3_ species *S. webbii* and significantly higher than those of C_4_ species *S. oppositifolia*. There is no evidence for constantly lower Γ values in populations that originated from more arid regions, although individuals of *S. verticillata* and *S. deschaseauxiana* mostly showed somewhat lower values, many individuals from similar climatic environments (e.g., *S. gymnomaschala* or individuals of *S. divaricata* from Fuerteventura or Lanzarote) showed higher values which skews a possible relation of climate and lower Γ values. Sage *et al*. (2014) suggested a Γ value below 10 µmol CO_2_ mol^-1^ characterizes the transition from C_2_ Type I to C_2_ Type II and represents an evolutionary progression towards C_4_. An accessory C_4_ metabolic cycle is present in C_2_ Type II plants and was so far only observed in several species of *Flaveria and Mollugo verticillata* (Edwards & Ku, 1987; Sage et al., 2018). None of the Γ measurements of 53 individuals of the *Salsola divaricata* agg. showed an indication of an accessory C_4_ metabolic cycle and a significantly higher carboxylation by PEPC, therefore indicating a stable C_2_-state for *S. divaricata* agg.

So far, the analyzed traits seem to be independent between populations and islands and show substantial variation. On Lanzarote for example, populations of *S. divaricata* have comparably high Γ values and medium to low carbon isotope values compared to other populations.

### PEPC activity has little or no correlation with Γ values or carbon isotope values in *S. divaricata* agg

The PEPC activity of the *S. divaricata* agg. was similar to the C_3_ species *S. webbii*. Even though there was a high standard deviation within C_4_ *S. oppositifolia*, none of the C_2_ individuals comes even close to a C_4_-like value. The activity of PEPC for *S. verticillata* and *S. deschaseauxiana* was slightly lower than those for *S. divaricata*; therefore, their slightly lower Γ values and slightly higher carbon isotope values are not due to increased CO_2_ fixation by an optimized PEPC. Thus, S. *verticillata* and *S. deschaseauxiana* do not seem to be closer to a more C_4_-like state than *S. divaricata* or *S. gymnomaschala*.

The PEPC activity in the *S. divaricata* agg. is highly variable and showed only a weak correlation with carbon isotope values and CO_2_ compensation points, as discussed above for *S. verticillata*. We would expect a higher PEPC activity in the salt treatment or the more arid locations e.g., Lanzarote or Fuerteventura, however that was not the case. Although there were statistically significant differences between islands, it is not possible to predict a value for PEPC activity given the location since there is generally a large overlap of values. The differences between populations could not be explained by the climatic context of the parent plant in the wild and they were all treated the same in the greenhouse. We could only detect a very weak correlation between PEPC activity, CO_2_ compensation points and carbon isotope values, supporting a rather independent distribution of the traits and their high phenotypic variation.

The independent variation in three photosynthetic traits (carbon isotope value, CO_2_ compensation point and PEPC activity) does not support a directed selection towards C_2_ Type II or C_4_-like photosynthesis in any of the populations studied, not even those growing in semi-desert conditions. In contrast, our trait observations are more in line with the results of hybridization experiments. Summarizing the physiology of advanced generations of interspecific C_3_ and C_4_ hybrids, Brown & Bouton (1993) stated that the ‘correlation among photosynthetic traits were low’ which indicates ‘a high degree of independence, both genetic and physiological, among C_4_ traits.’ Like in our study, the hybridization experiment of C_3_ and C_4_ *Atriplex* (Oakley *et al*., 2014) showed a distinct range of Γ values (in nine F_2_ individuals from 25-45 µmol CO_2_ mol^-1^; and one C_4_-like value). Brown & Bouton (1993) hypothesize that some of the photosynthetic traits measured can be more C_4_-like in a still functioning C_3_ metabolism, such as Γ, while others just mirror the physiological C_3_ default condition, such as enzyme activity. This together with the phylogenomic results and photosynthetic gene tree analyses (see below) supports the hypothesis of a hybrid origin of the *Salsola divaricata* agg. and subsequent stabilizing selection forming an evolutionary stable C_2_ lineage.

### First phylogenomic evidence of a hybrid origin of the *S. divaricata* agg

The Salsoleae species tree showed a high level of conflict including the *S. divaricata* agg, despite high bootstrap support. The assessment of hybridization events using PhyloNet and HyDe showed multiple hybridization events within Salsoleae in which the *Salsola divaricata* agg. was likely involved. The incomplete sampling of Salsoleae does not allow us to identify the exact parental lineages; but a hybrid origin of the *Salsola divaricata* agg. from C_3_ and C_4_ parental lineages was highly supported.

The network that was recovered most often showed an ancient hybridization event involving an ancestor of C_4_ *Salsola soda* and the C_3_ proto-kranz *Salsola montana* giving rise to the *Salsola divaricata* agg. clade. This is the first phylogenomic evidence that a C_3_-C_4_ intermediate species is the result of (ancient) hybridization between a C_4_ and a C_3_ lineage. The second reticulation event later indicates a hybridization event involving the ancestor of *Salsola divaricata* agg. and the C_4_ *Salsola soda* giving rise to the C_4_ clade formed by *Salsola oppositifolia, Halogeton, Hammada*, and *Haloxylon*. This might indicate that this relative old C_4_ lineage (Salsoleae ∼ 21–22 MYA; Morales-Briones et al., 2021), could have inherited C_4_ or antecedent traits from a parental lineage giving them an adaptive advantage that might have influenced their photosynthesis pathway, e.g., adaptations in photosynthesis or photorespiration associated enzymes or Kranz-cells. This is in line with recent findings in the genus *Flaveria* (Morales-Briones and Kadereit, 2022) where ancient hybridization was found to be a step towards C_4_ evolution and likely promoted the rapid acquisition of C_4_ traits. We also found hybridization events in the two clades in the other networks with similar parental lineages, supporting the evidence that these lineages were involved in hybridization events. This could indicate either an incomplete reproductive barrier in the early lineages and recurrent backcrossing or incidences of horizontal gene transfer. We must keep in mind though that our sampling is limited and that other lineages not sampled in the C_4_ clade might also play a role (see Schüssler *et al*., 2017 for a plastid tree of the Salsoloideae).

### Phylogenetic analysis of photosynthetic genes supports the hybrid origin hypothesis and suggests candidate genes for further study

A closer look of 50 gene trees from genes or gene families involved in photosynthesis or photorespiration revealed the same overall pattern, namely incongruence regarding the position of the *Salsola divaricata* agg., which in most cases either groups with the C_3_ *S. montana* (also the case in the plastid dataset) or with the C_4_ species. This pattern could be explained by hybridization of parental C_3_ and C_4_ lineages giving rise to an allopolyploid *S. divaricata* agg. with an intermediate expression of photosynthetic traits. The Kranz-like salsoloid leaf type of the *S. divaricata* agg. (Schüssler *et al*., 2017: Fig. 7), for example, might occur as in the case of the gene SHORTROOT, a C_4_ associated transcription factor which influences the development of Kranz cells (Kelly *et al*., 2017, Slewinski *et al*. 2014), with expression of the parental C_4_ copy. Studying transcriptome profiles of Salsoleae and Camphorosmeae leaves, respectively, Lauterbach *et al*. (2017a) and Siadjeu *et al*. (2021) revealed a high expression of SHORTROOT in C_2_ species. In the case of PEPC, two copies were recovered from our transcriptomes and for both the *S. divaricata* agg. group with the C_3_ species supporting the C_3_-like PEPC activity data found in this study.

## Conclusions

The phylogenetic signal in our trees, gene trees and networks indicate hybridization events gave rise to the *Salsola divaricata* agg. Furthermore, the distinct variation of three photosynthetic traits within a C_3_-C_4_ intermediate range without indication of selection towards a more C_4_-like expression in this relatively old lineage supports the interpretation of an evolutionarily stable C_2_ lineage of hybrid origin. Overall, the photosynthetic traits seem plastic and genetically fixed stress reactions are not prominent among the populations studied. The *Salsola divaricata* agg. is well adapted to a broad range of climatic conditions including mesic to arid coastal habitats and to salinity. Part of the success of the species aggregate might be the photorespiratory pump acquired through hybridization with a C_4_ lineage in the past. However, a broader sampled phylogenomic study is needed to further narrow down the parental lineages. We propose that reticulation events might play an important role in the multiple convergent acquisition of complex traits such as C_2_ or C_4_ photosynthesis as a possible fast track.

## Conflict of Interest

The authors declare that the research was conducted in the absence of any commercial or financial relationships that could be construed as a potential conflict of interest.

## Author Contributions

GK designed the study. DT and GE (carbon isotope values) measured the trait data and DT analyzed them. DT, DMB and YY conducted the phylogenomic analyses. DMB prepared the figures and DT the map. DT, GK and DMB wrote the first draft. All authors contributed to the discussion of the results and the final version of the manuscript.

## Funding

Funding for this project came from the German Science Foundation (DFG grant KA1816/9-1) and the University of Minnesota.

## Acknowledgements

We thank the collectors of seeds of the *Salsola divaricata* agg. for their great help to start this project, in particular R. Barone, V. Boehlke, H. Freitag, J. Gil González, F. Hernández, M. Olangua Corral, S. Scholz and E. Voznesenskaya. We are grateful to C. Wild for cultivating and maintaining the plants for this project. For conducting a part of the carbon isotope measurements, we thank M. Maus (Geology Department University of Mainz). The Minnesota Supercomputing Institute provided access to computational resources.

## Data Availability Statement

Transcriptome data generated for this study can be found in the NCBI Sequence Read Archive (SRA) (see **Table 1** for SRA accession numbers).

The following Supporting Information is available for this article:

**Fig. S1:** Carbon isotope values (δ13C[‰]) of different populations (P) of the *Salsola divaricata* agg. on the Canary Islands and Morocco.

**Fig. S2:** CO2 compensation points (Γ in µmol CO2 mol-1) of populations (P) of the *Salsola divaricata* agg.

**Fig. S3:** a) Cladogram of Salsoloideae inferred from the Maximum likelihood analyses of the concatenated 991-nuclear gene supermatrix. b) Cladogram of Salsoloideae inferred from the ASTRAL analyses of 991 nuclear genes trees.

**Fig. S4:** A) RAxML cladogram of Salsoloideae inferred from the concatenated 991-nuclear gene supermatrix. B) ASTRAL Cladogram of Salsoloideae inferred from 991 nuclear genes trees.

**Fig S5:** Maximum likelihood phylogeny of Salsoloideae s.l. inferred from the IQ-TREE analysis of complete and partial plastomes.

**Fig. S6:** Maximum pseudo-likelihood species network inferred with PhyloNet with up to five hybridization events.

**Fig. S7:** HyDe boxplot of the distribution of the admixture parameters (γ) from the 298 significant tests.

**Fig. S8:** Maximum likelihood cladograms of the 50 photosynthetic gene trees. Bootstrap support (BS) values are shown above the branches.

**Table S1:** Populations (named by living collection number in the Botanical Garden Mainz; see Fig. 1) included in the carbon isotope measurements, CO2 compensation point measurements and PEPC activity measurements, including sampling location and collector.

**Table S2:** Gene and character occupancy of the concatenated matrix.

**Table S3:** HyDe results from the 298 significant tests.

**Table S4:** Fifty trees of photosynthesis pathway-associated genes characteristic.

**Materials and Methods S1:** More detailed description of Plant material, methods of CO_2_ compensation point, PEPC activity measurement, Transcriptome processing and nuclear phylogenetic analyses, Assessment of hybridization, Plastome assembly and phylogenetic analysis, and Analysis of photosynthetic gene trees.

## Notes

### Competing Interest Statement

The authors have declared no competing interest.

### Summary of Updates

We rephrased the general premises and did some rewording in the discussion and conclusion. We added a correlation analysis and updated some of the figures for readability. We added Fig.7. We moved details from the supplemental material to the main text. Additionally, we have revised the entire manuscript to streamline the text.

