## Supporting Information for "On the hybrid origin of the C_2_ *Salsola divaricata* agg. (Amaranthaceae) from C_3_ and C_4_ parental lineages"

Article acceptance date:

The following Supporting Information is available for this article:

**Fig. S1:** Carbon isotope values (δ13C(‰)) of different populations (P) of the *Salsola divaricata* agg. on the Canary Islands and Morocco
**Fig. S2:** CO2 compensation points (Γ in µmol CO2 mol-1) of populations (P) of the *Salsola divaricata* agg.
**Fig. S3:** a) Cladogram of Salsoloideae inferred from the Maximum likelihood analyses of the concatenated 991-nuclear gene supermatrix. b) Cladogram of Salsoloideae inferred from the ASTRAL analyses of 991 nuclear genes trees.
**Fig. S4:** A) RAxML cladogram of Salsoloideae inferred from the concatenated 991-nuclear gene supermatrix. B) ASTRAL Cladogram of Salsoloideae inferred from 991 nuclear genes trees.
**Fig S5:** Maximum likelihood phylogeny of Salsoloideae s.l. inferred from the IQ-TREE analysis of complete and partial plastomes.
**Fig. S6:** Maximum pseudo-likelihood species network inferred with PhyloNet with up to five hybridization events.
**Fig. S7:** HyDe boxplot of the distribution of the admixture parameters (γ) from the 298 significant tests.
**Fig. S8:** Maximum likelihood cladograms of the 50 photosynthetic gene trees. Bootstrap support (BS) values are shown above the branches.

**Table S1:** Populations (named by living collection number in the Botanical Garden Mainz; see Fig. 1) included in the carbon isotope measurements, CO2 compensation point measurements and PEPC activity measurements, including sampling location and collector.
**Table S2:** Gene and character occupancy of the concatenated matrix.
**Table S3:** HyDe results from the 298 significant tests.
**Table S4:** Fifty trees of photosynthesis pathway-associated genes characteristic.

**Materials and Methods S1:** More detailed description of Plant material, methods of $CO_2$ compensation point, PEPC activity measurement, Transcriptome processing and nuclear phylogenetic analyses, Assessment of hybridization, Plastome assembly and phylogenetic analysis, and Analysis of photosynthetic gene trees.

**References Materials and Methods S1**

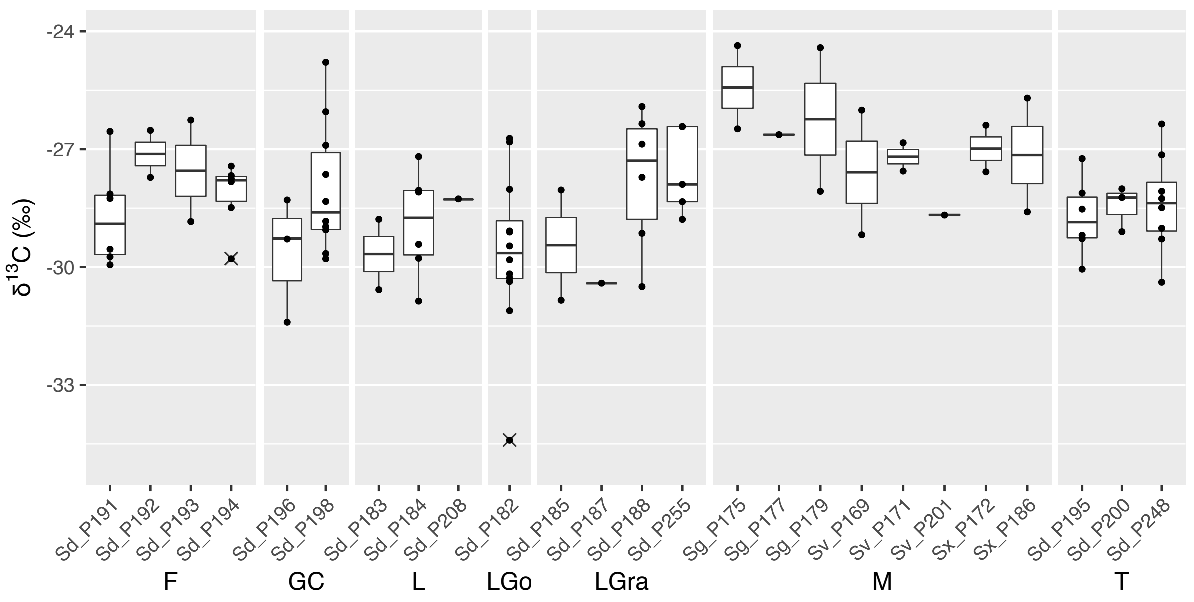


**Fig. S1:** Carbon isotope values (δ13C[‰]) of different populations (P) of the *Salsola divaricata* agg. on the Canary Islands and Morocco (only control treatment; Sd = *Salsola divaricata*, Sg = *S. gymnomaschala*, Sv = *S. verticillata*, Sx = *S. deschaseauxiana*; F = Fuerteventura, GC = Gran Canaria, L = Lanzarote, LGo = La Gomera, LGra = La Graciosa, M = Morocco, T = Tenerife). Due to very different sample sizes further statistical tests including all populations were inconclusive.

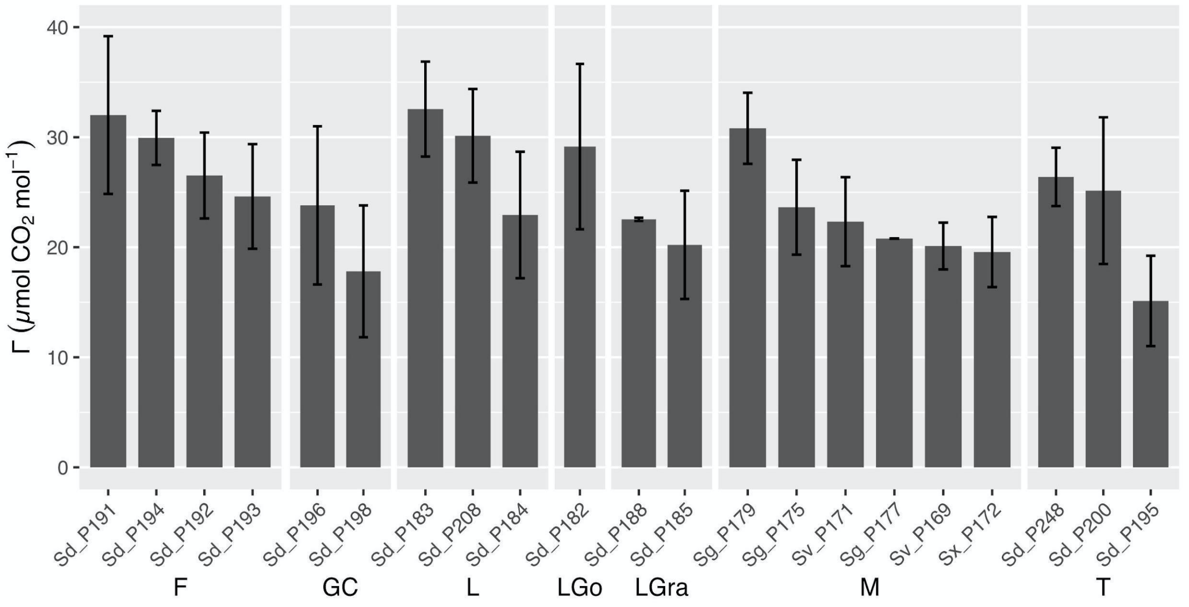


**Fig. S2:** CO2 compensation points (Γ in µmol CO2 mol-1) of populations (P) of the *Salsola divaricata* agg. sorted by island (Sd = *Salsola divaricata*, Sg = *S. gymnomaschala*, Sv = *S. verticillata,* Sx = *S. deschaseauxiana*; F = Fuerteventura, GC = Gran Canaria, L = Lanzarote, Go = La Gomera, LGra = La Graciosa, M = Morocco, T = Tenerife).

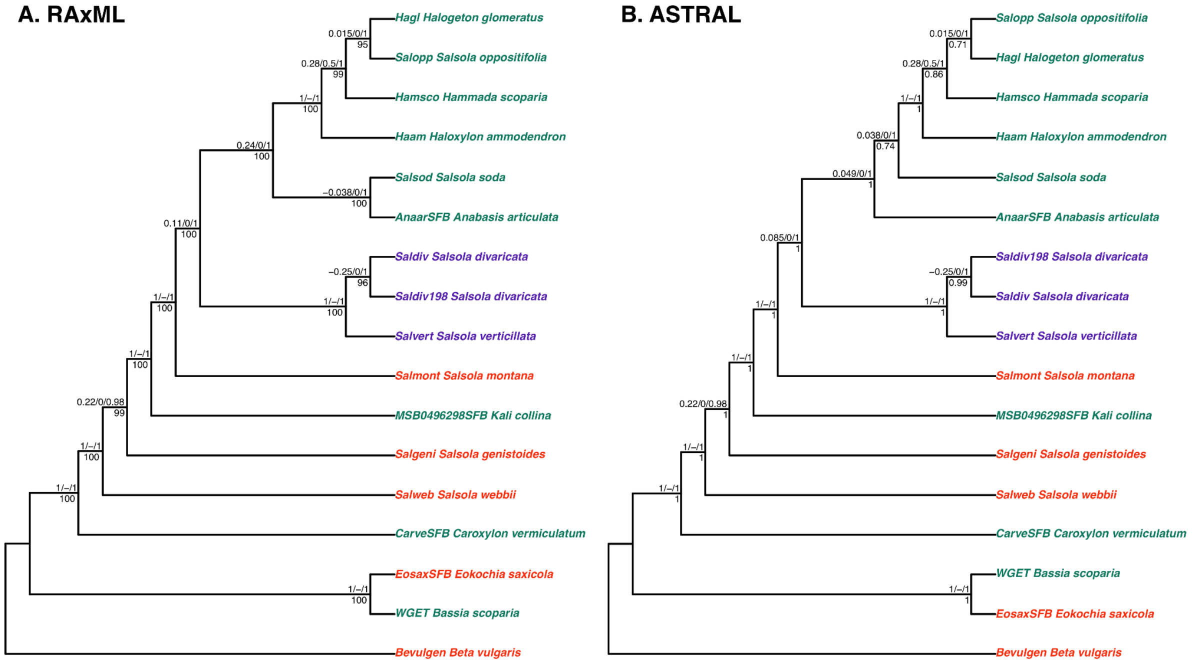


**Fig. S3:** a) Cladogram of Salsoloideae inferred from the Maximum likelihood analyses of the concatenated 991-nuclear gene supermatrix. Quartet Sampling scores (QS) and Bootstrap support (BS) values are shown above and below branches, respectively. b) Cladogram of Salsoloideae inferred from the ASTRAL analyses of 991 nuclear genes trees. Quartet Sampling scores (QS) and Local Posterior Probabilities (LPP) are shown above and below branches, respectively. QS scores: Quartet concordance/Quartet differential/Quartet informativeness.

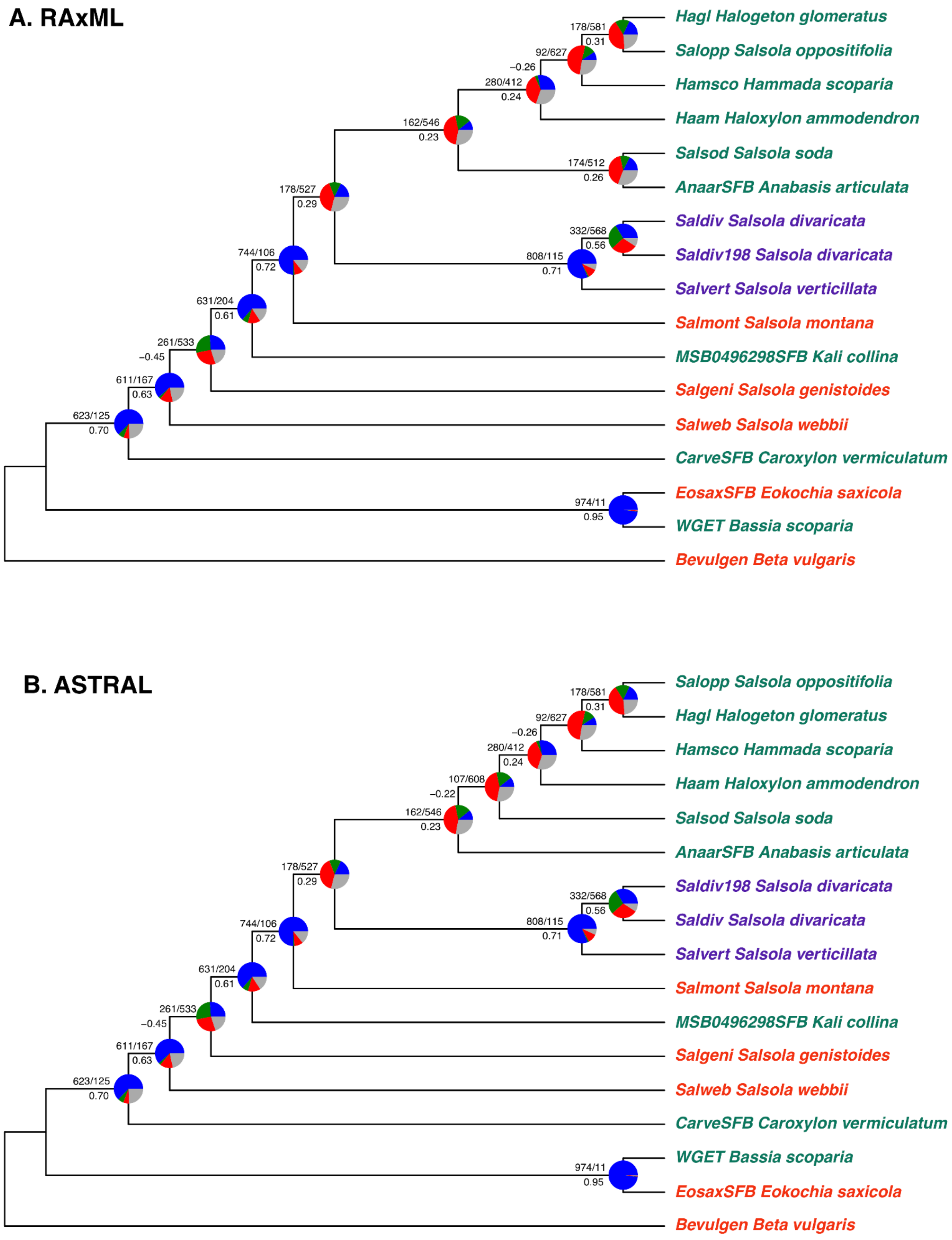


**Fig. S4:** A) RAxML cladogram of Salsoloideae inferred from the concatenated 991-nuclear gene supermatrix. B) ASTRAL Cladogram of Salsoloideae inferred from 991 nuclear genes trees. Numbers above branches indicate the number of gene trees concordant/conflicting with that node. Numbers below the branches are the Internode Certainty All (ICA) score. Pie charts represent the proportion of gene trees that support that clade (blue), the proportion that support the main alternative bifurcation (green), the proportion that support the remaining alternatives (red), and the proportion (conflict or support) that have < 50% bootstrap support (gray).

cpDNA

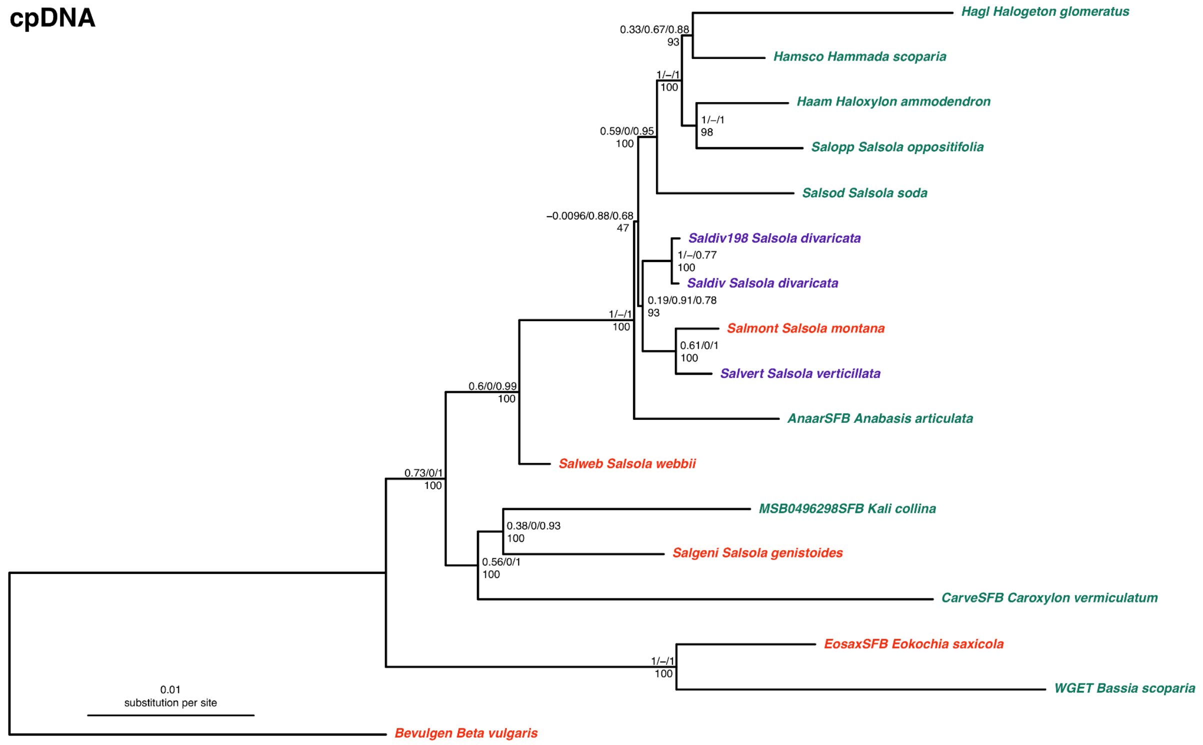


**Fig S5:** Maximum likelihood phylogeny of Salsoloideae s.l. inferred from the IQ-TREE analysis of complete and partial plastomes. Quartet Sampling (QS) scores are shown above the branches. QS scores: Quartet concordance/Quartet differential/Quartet informativeness. Bootstrap support (BS) values are shown below the branches. Branch lengths are proportional to substitutions per site (scale bar on the bottom).

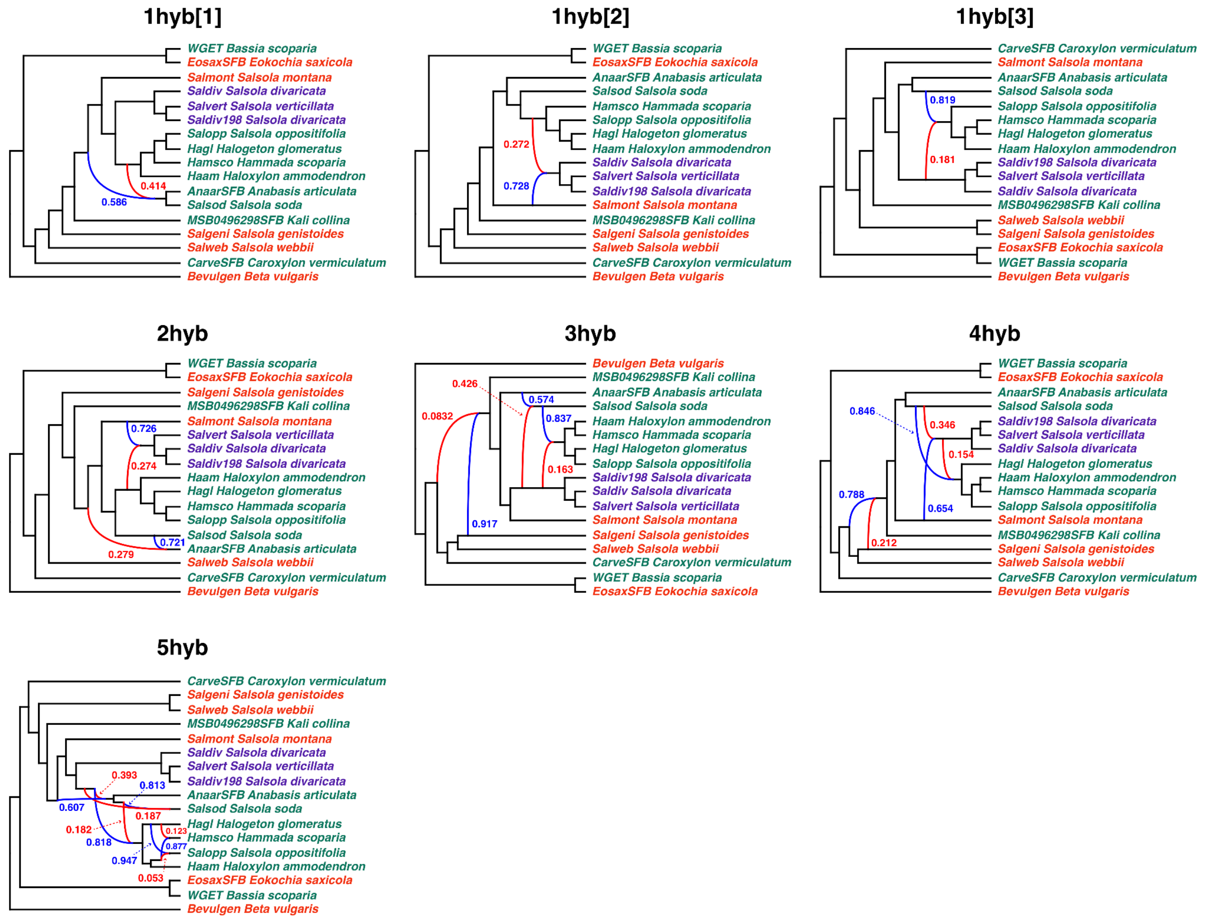


**Fig. S6:** Maximum pseudo-likelihood species network inferred with PhyloNet with up to five hybridization events. The top three networks are shown for the search with one hybridization event. Red and blue curved branches indicate the minor and major edges, respectively of hybrid nodes. Numbers next to curved branches indicate inheritance probabilities for each hybrid node.

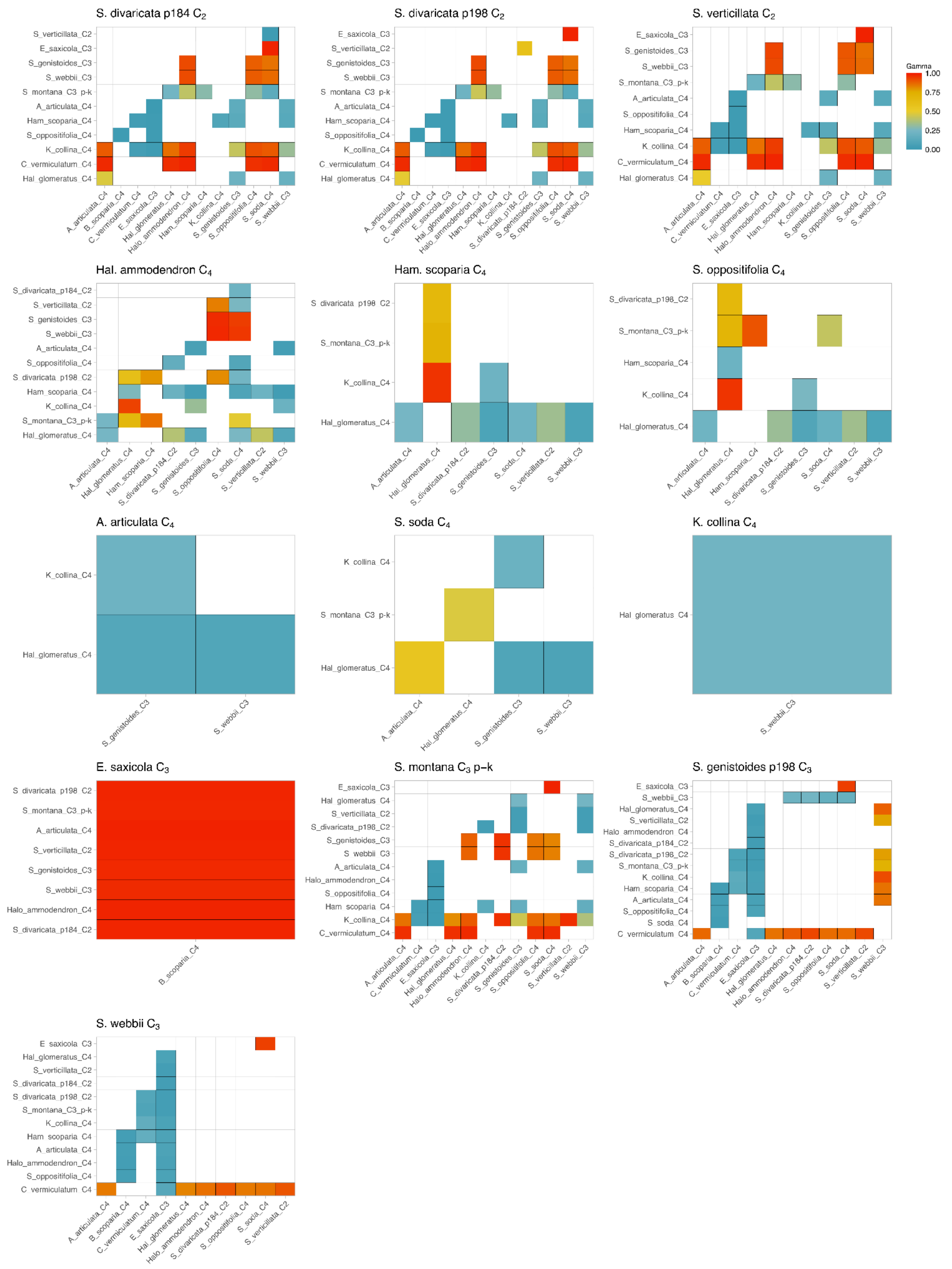


**Fig. S7:** HyDe significant hybridization tests (298). Species on the x-axis are the parental lineage 1 (P1) and species on the y-axis represent the parental lineage 2 (P2). Only colored boxes denote possible combinations of P1 and P2 to produce the hybrid species. The color scale represents the level of hybridization parameter ($\gamma$) for each hybridization event. Recent 50:50 hybrids would be represented by a $\gamma$ ~0.5. Levels of the hybridization parameter approaching 0 indicate a major hybrid contribution from P1, and levels approaching 1 indicate a major hybrid contribution from P2, with both cases representing ancient hybridization.

**Fig. S8.** Maximum likelihood cladogram of the 50 photosynthetic gene trees. Bootstrap support (BS) values are shown above the branches.

**PEP carboxylase kinase (PEPC–K1) – AT1G08650**

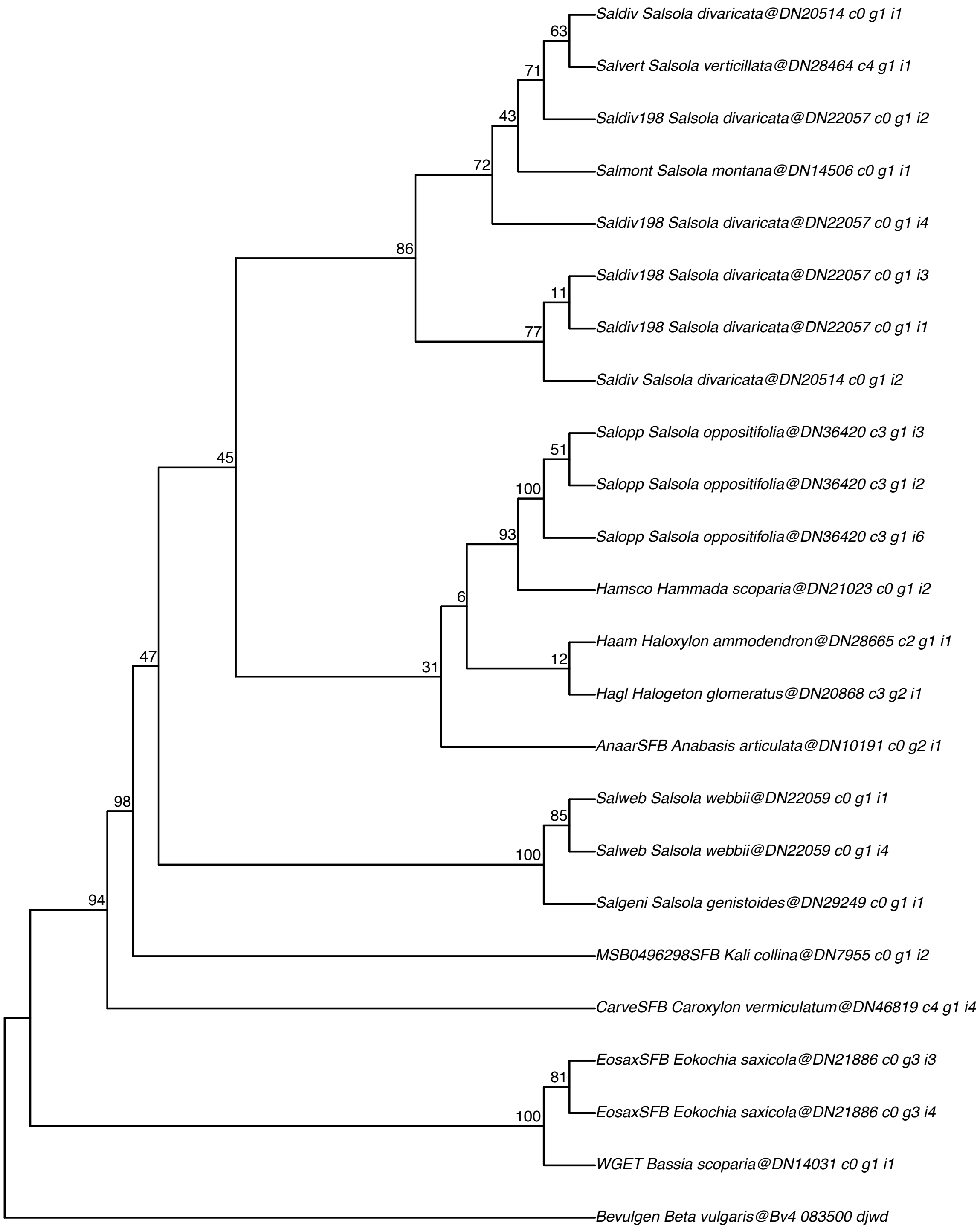

#### Alanine aminotransferase (Ala–AT1)–AT1G17290

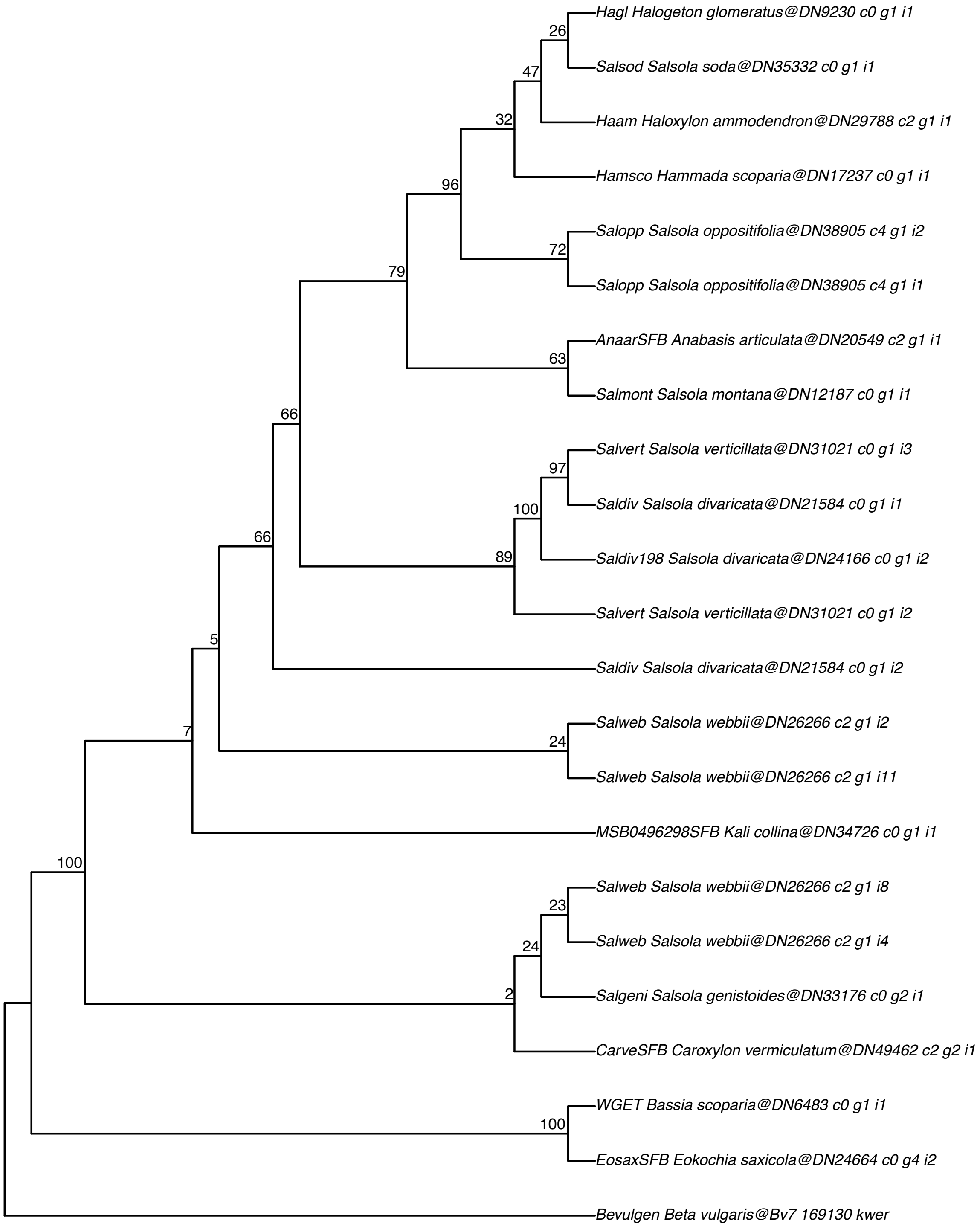

#### Beta carbonic anhydrase (BCA3)–AT1G23730

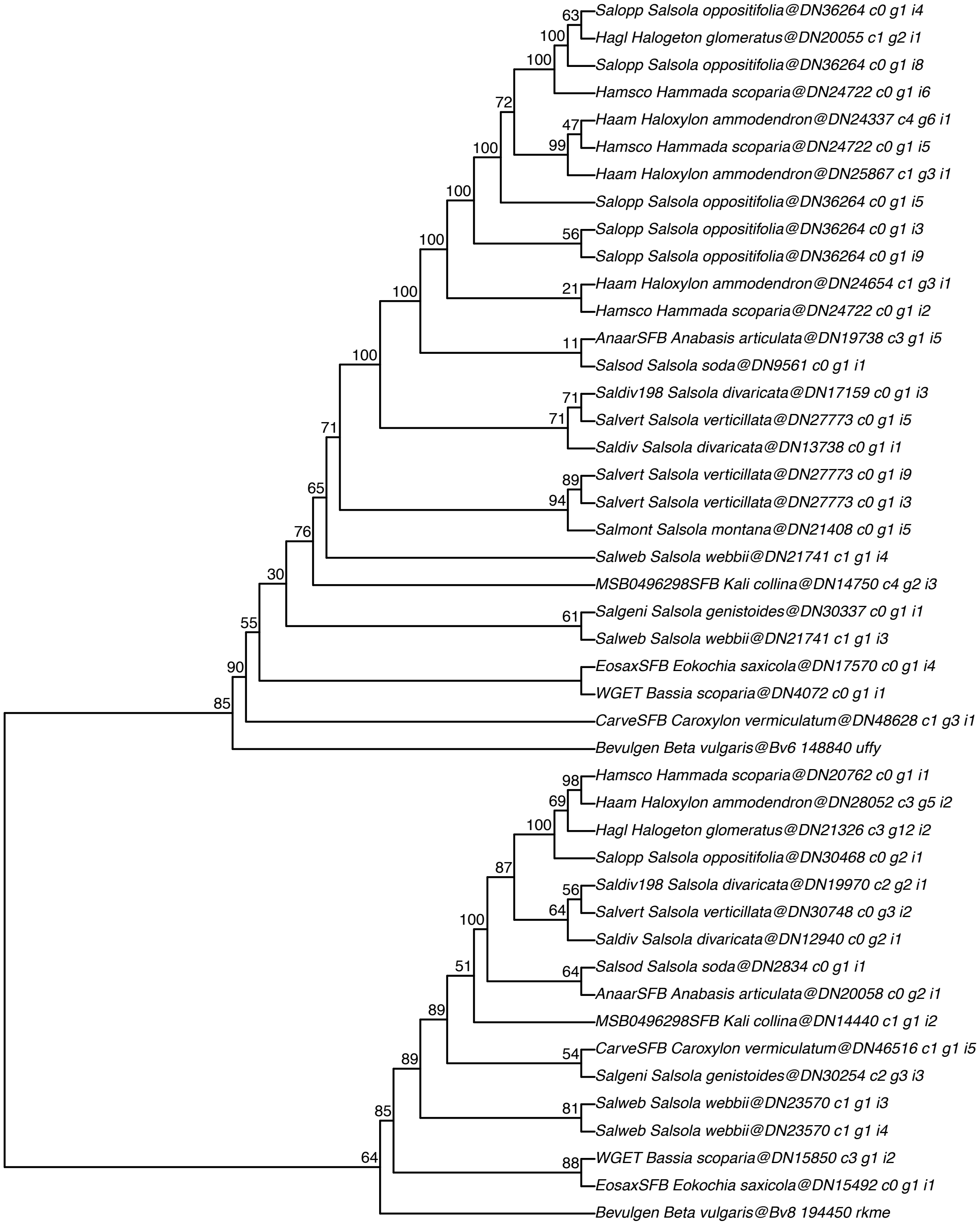

#### Sodium:hydrogen antiporter (NHD2)–AT1G49810

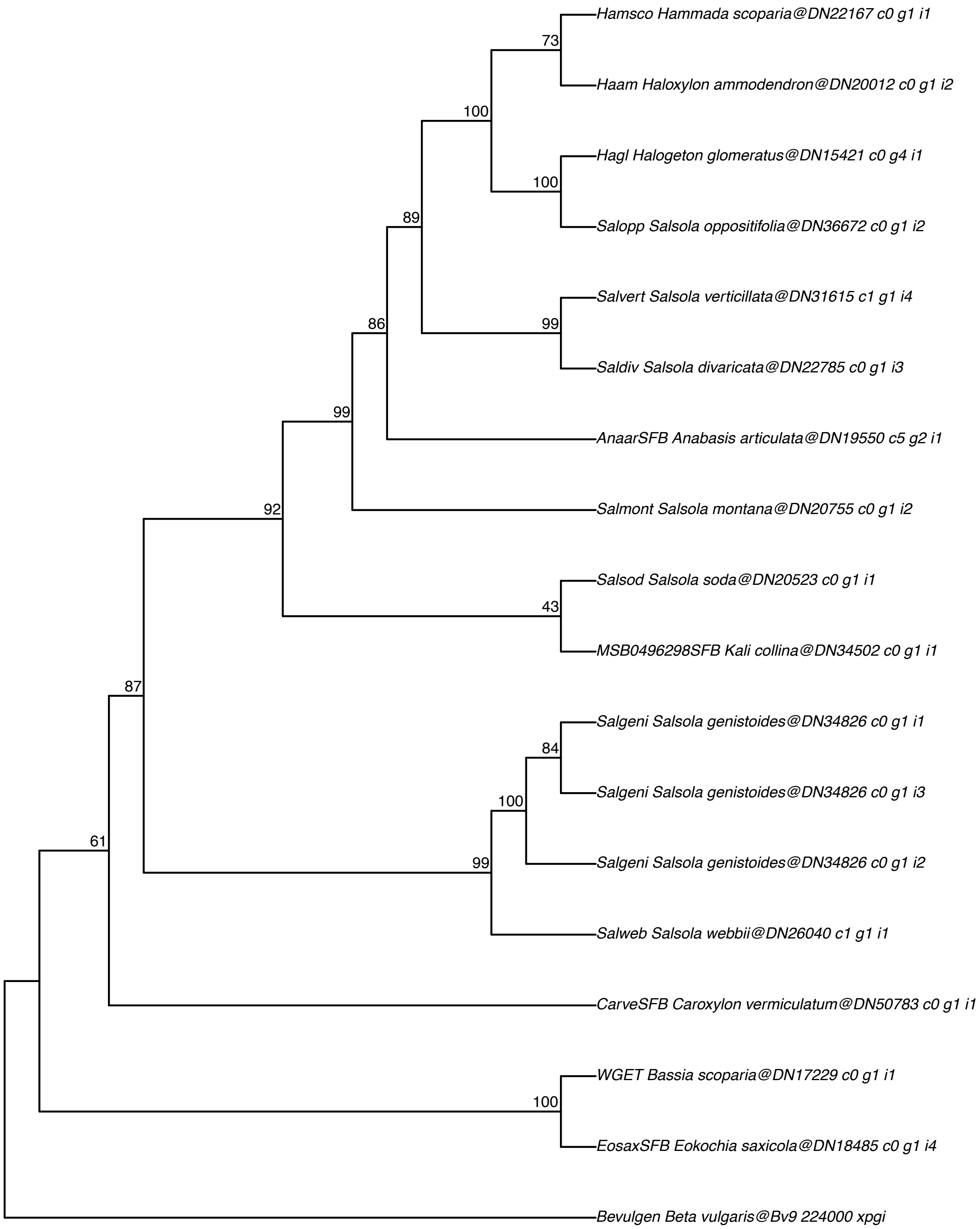

#### PEP carboxylase (PEPC1)–AT1G53310

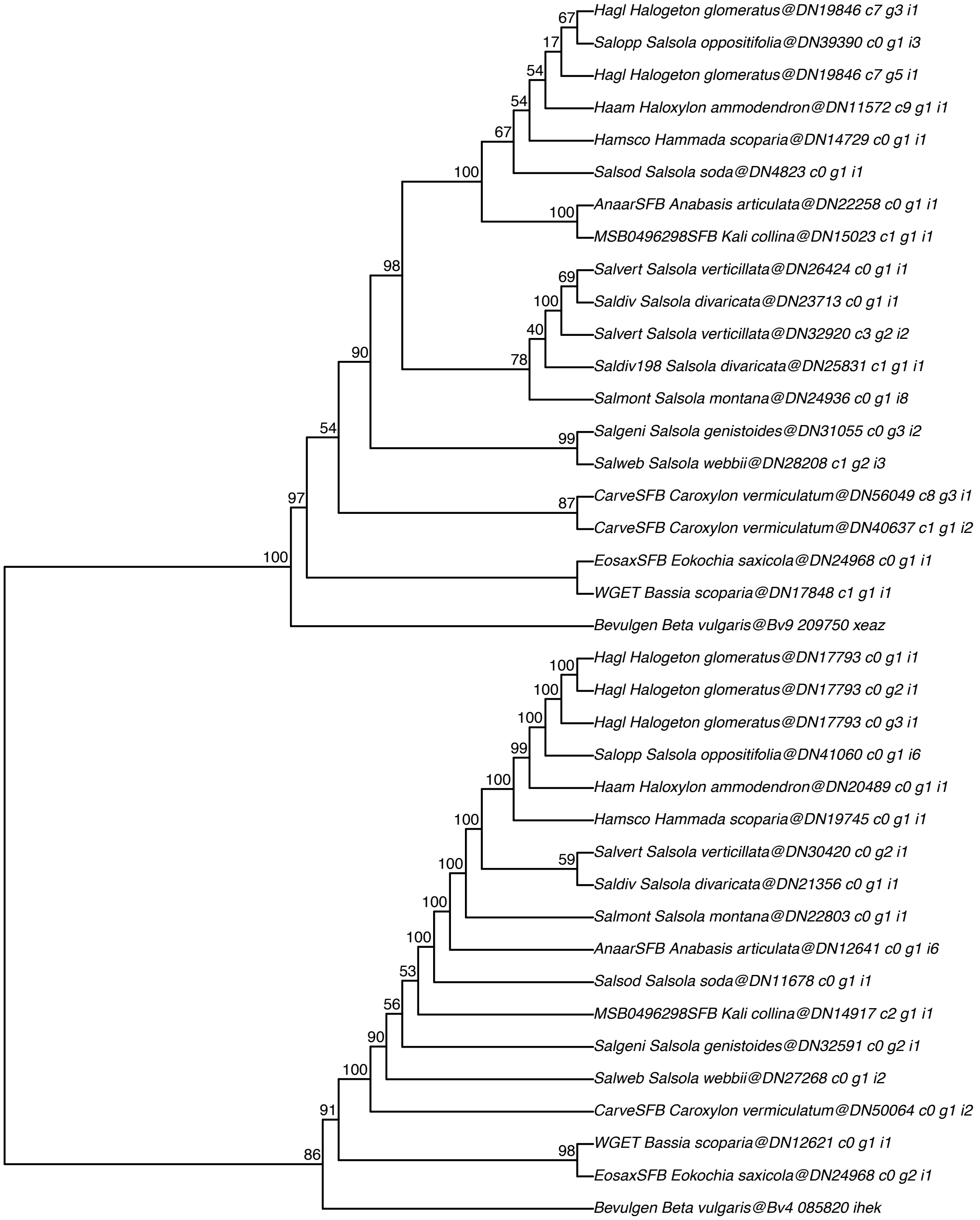

#### NADP–dependent malic enzyme (NADP–ME4)–AT1G79750

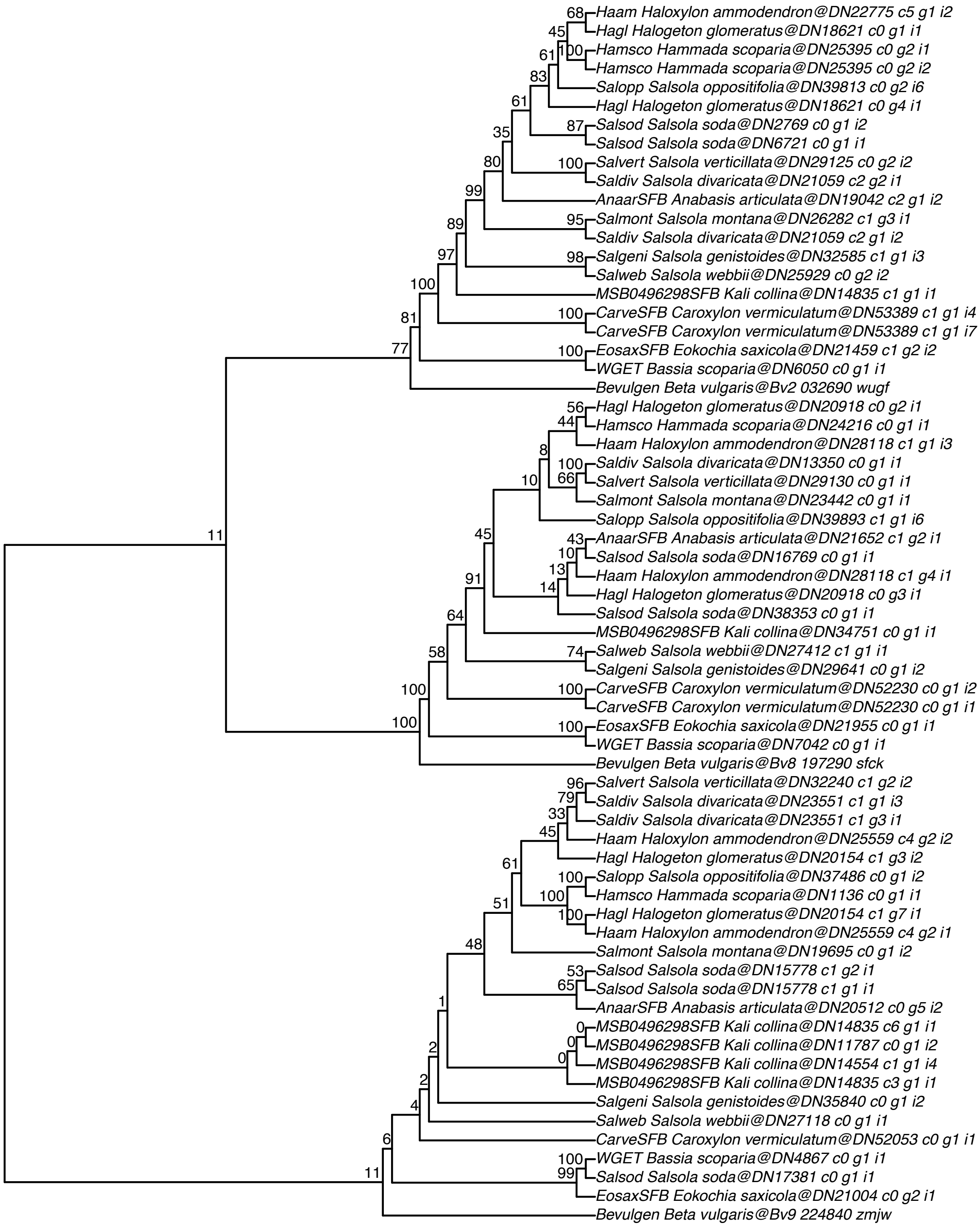

#### BEL1–like homeo domain 7–AT2G16400

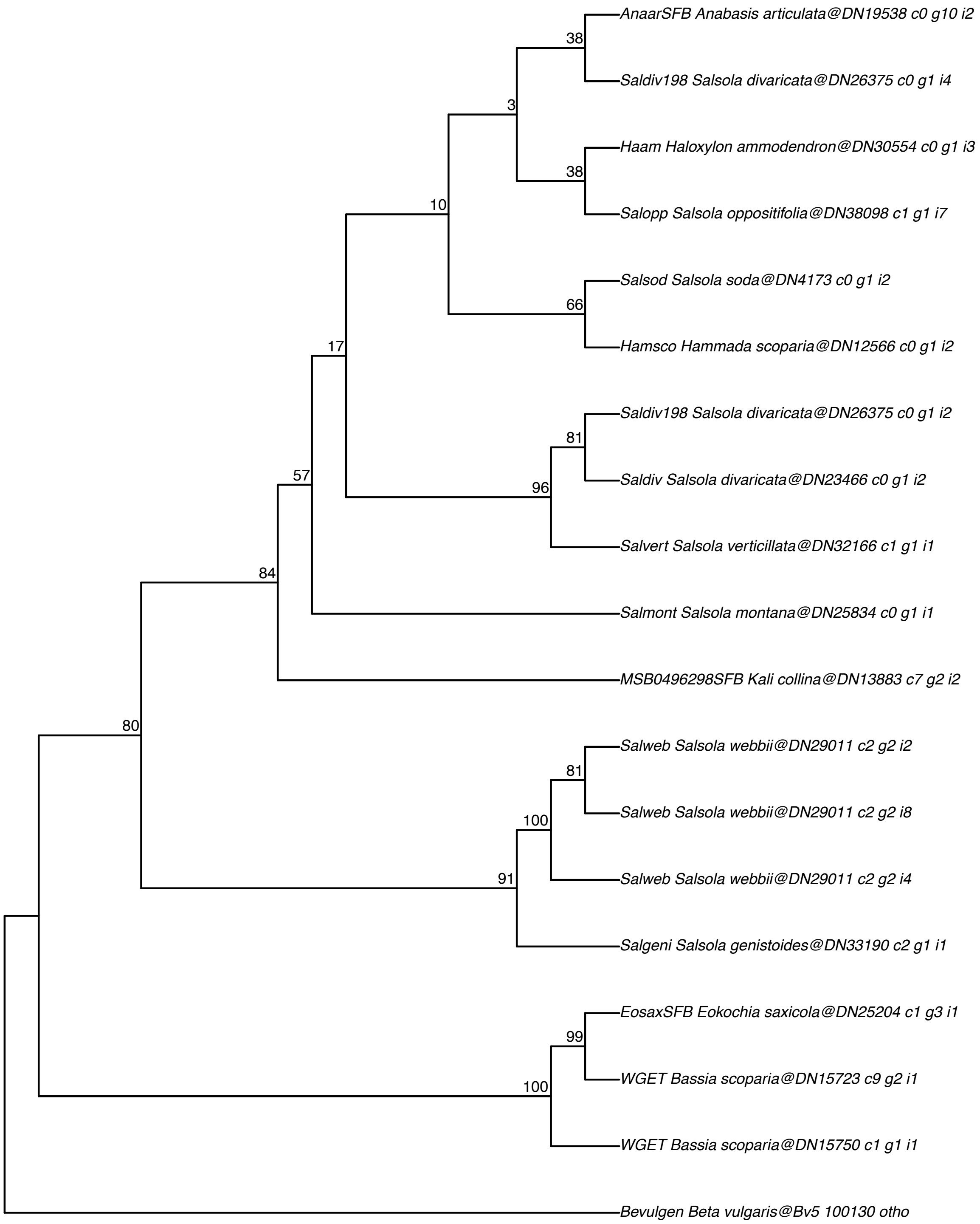

#### NADP–dependent malate enzyme (NADP–ME1)–AT2G19900

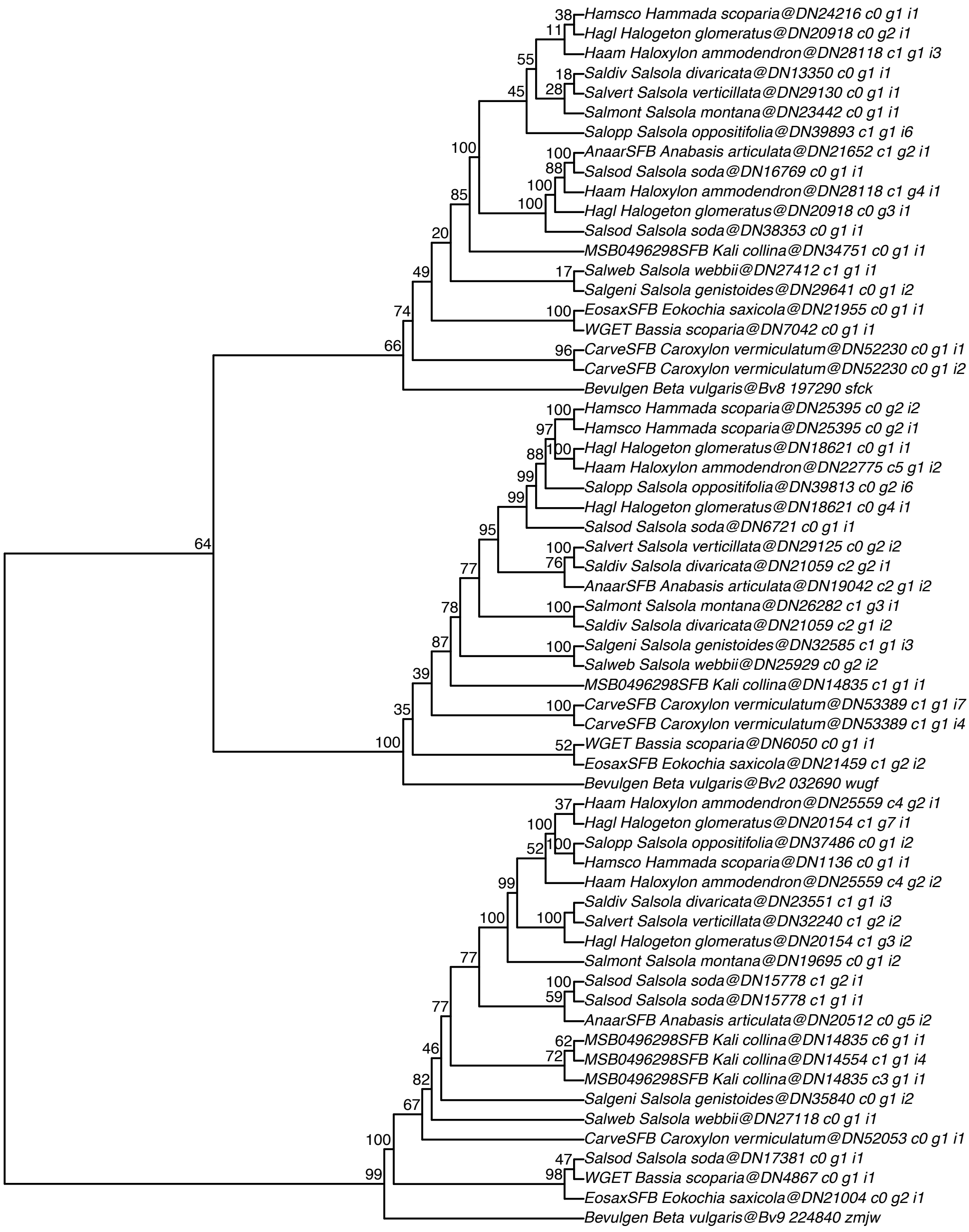

#### Bile acid:sodium symporter family protein (BASS2)–AT2G26900

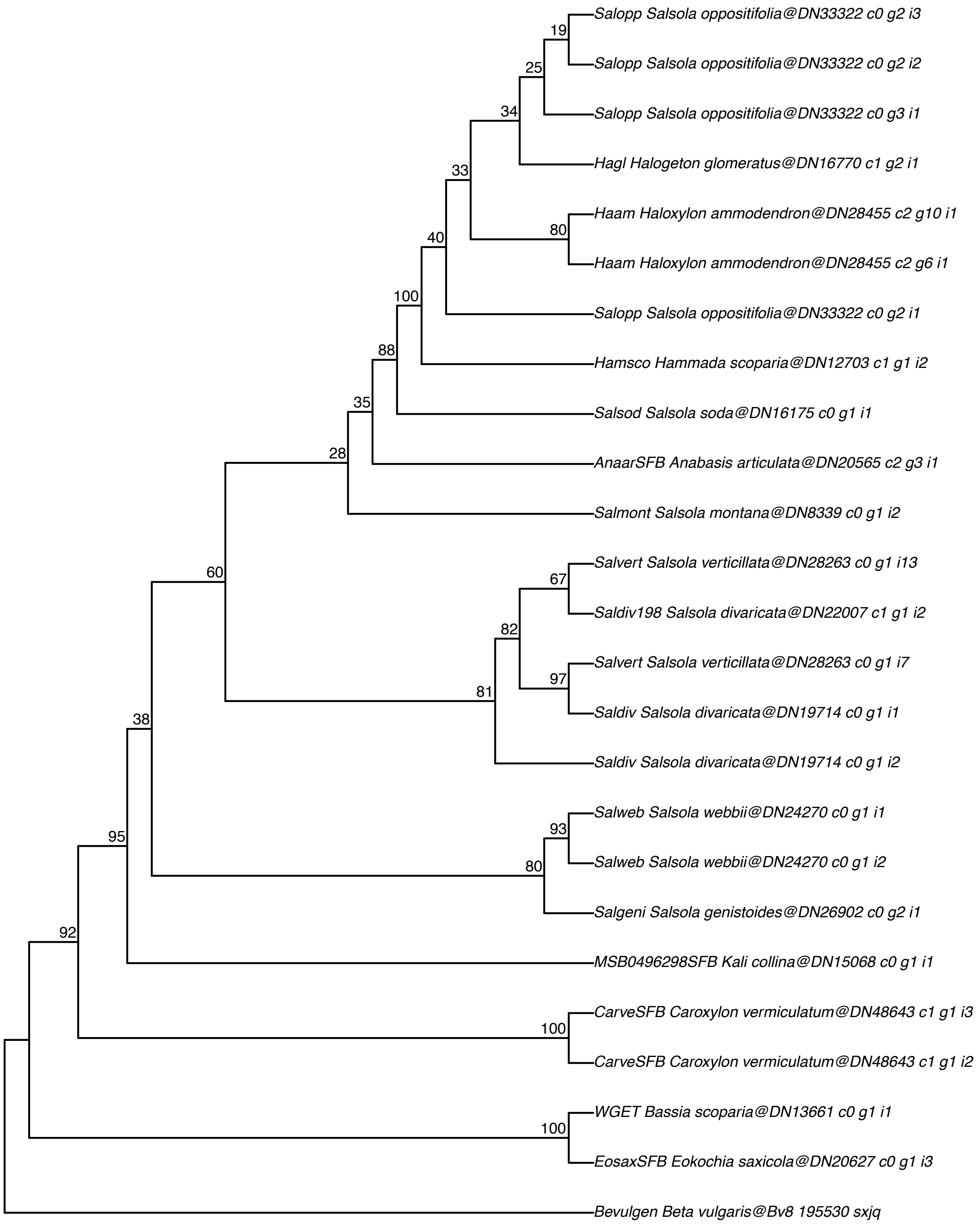

**PEP carboxylase (PEPC2)–AT2G42600**

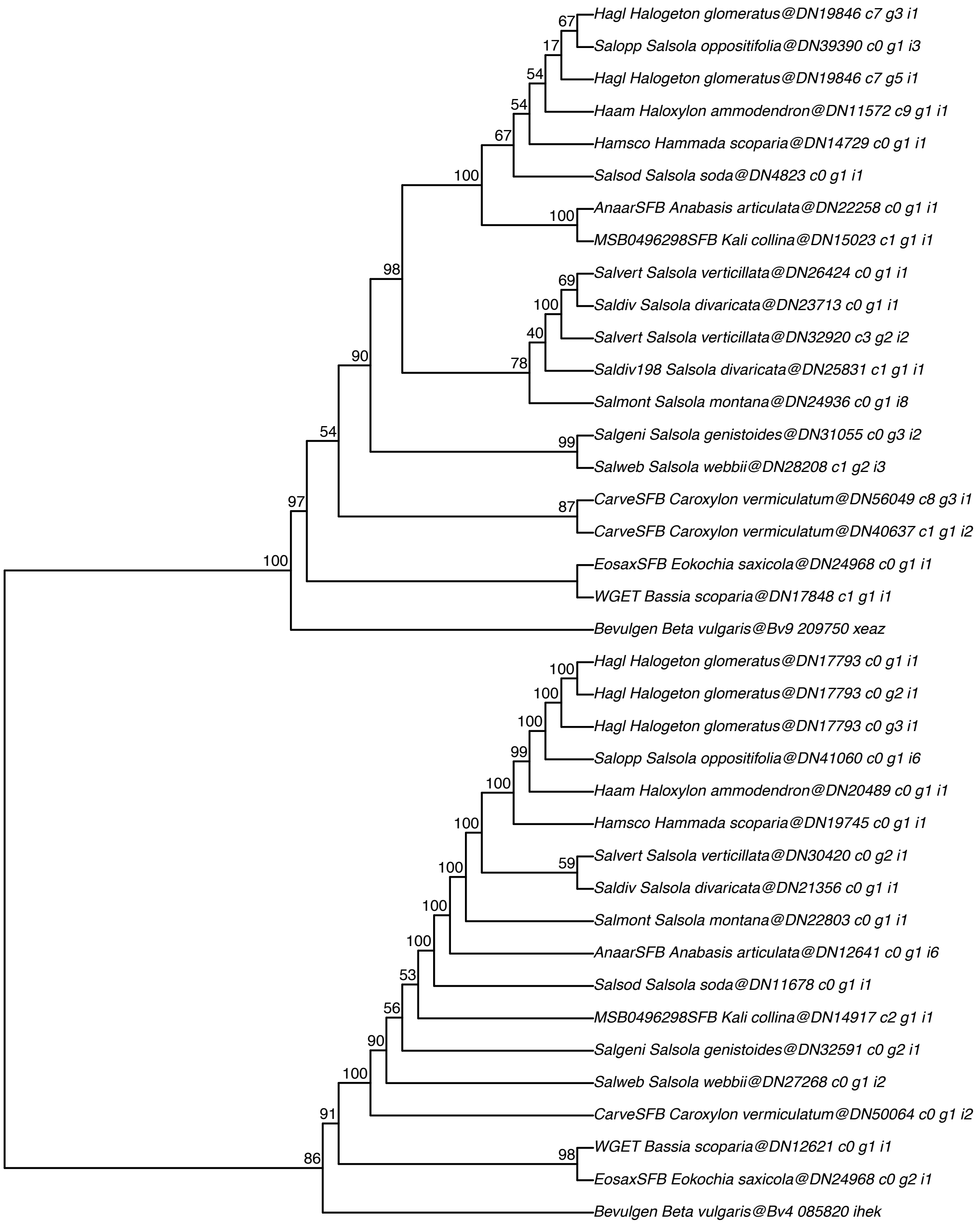

**PPdK−related protein (PPdK−RP2)−AT3G01200**

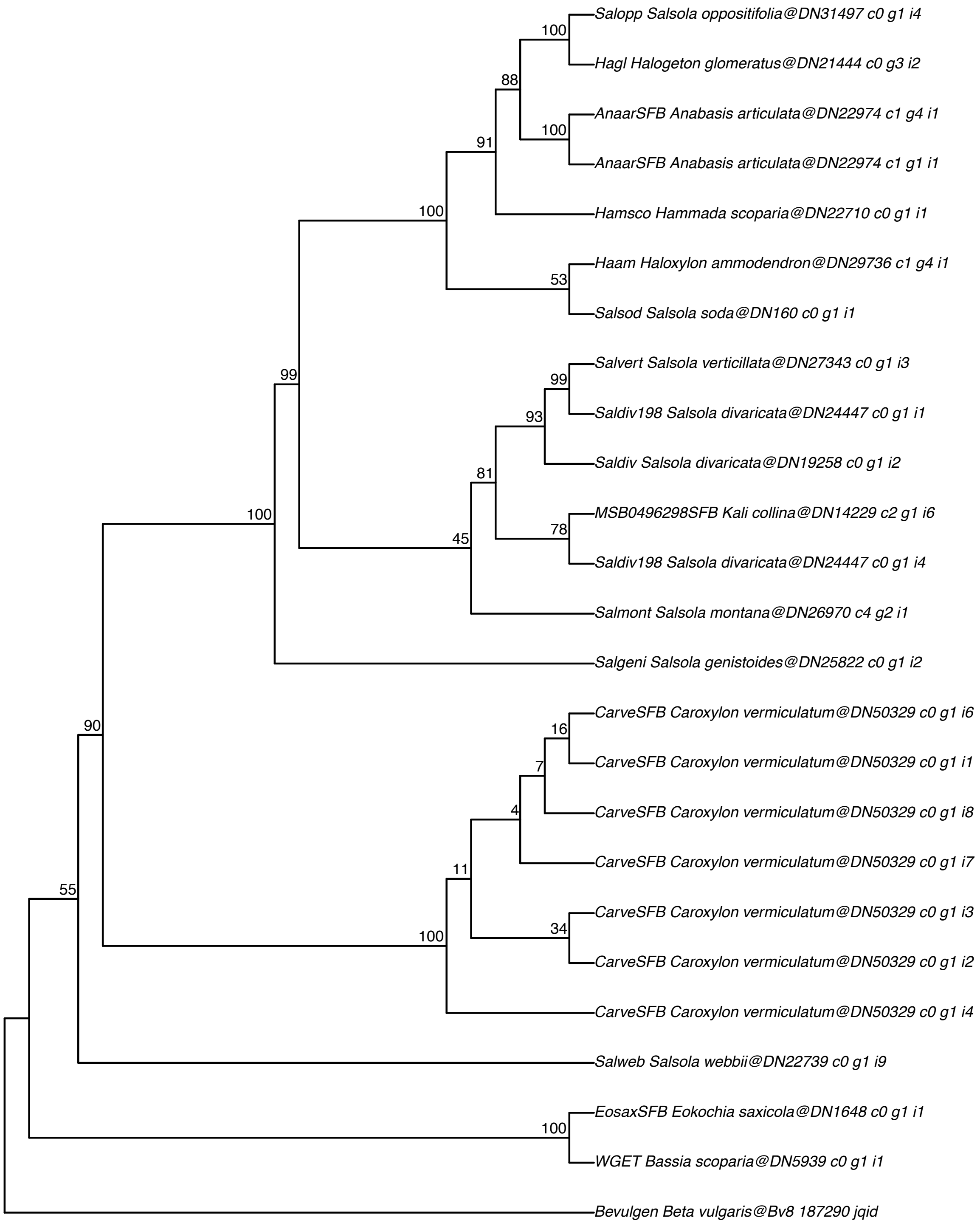

#### PEP/phosphate translocator (PPT2)–AT3G01550

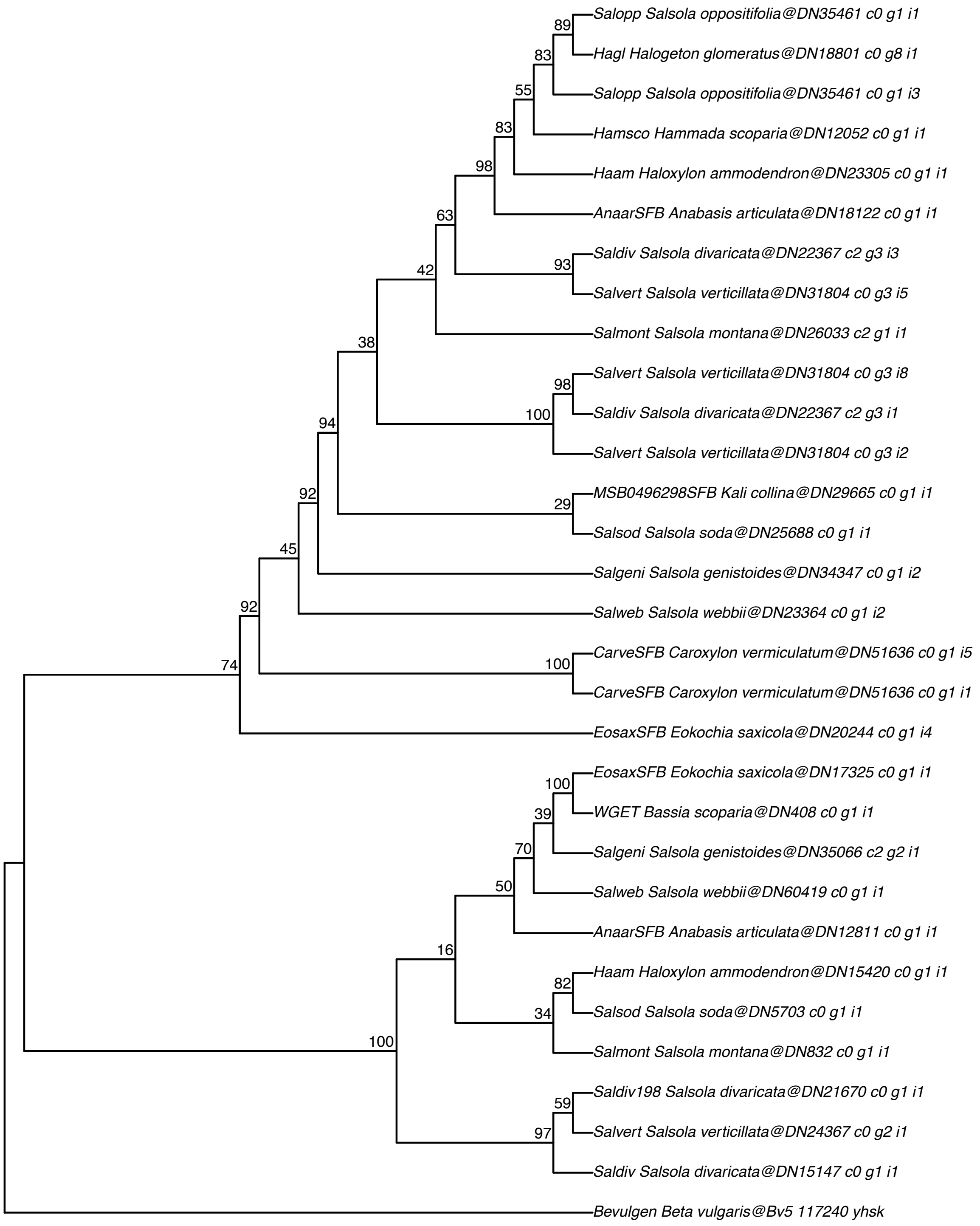

#### PEP carboxylase kinase (PEPC−K2)−AT3G04530

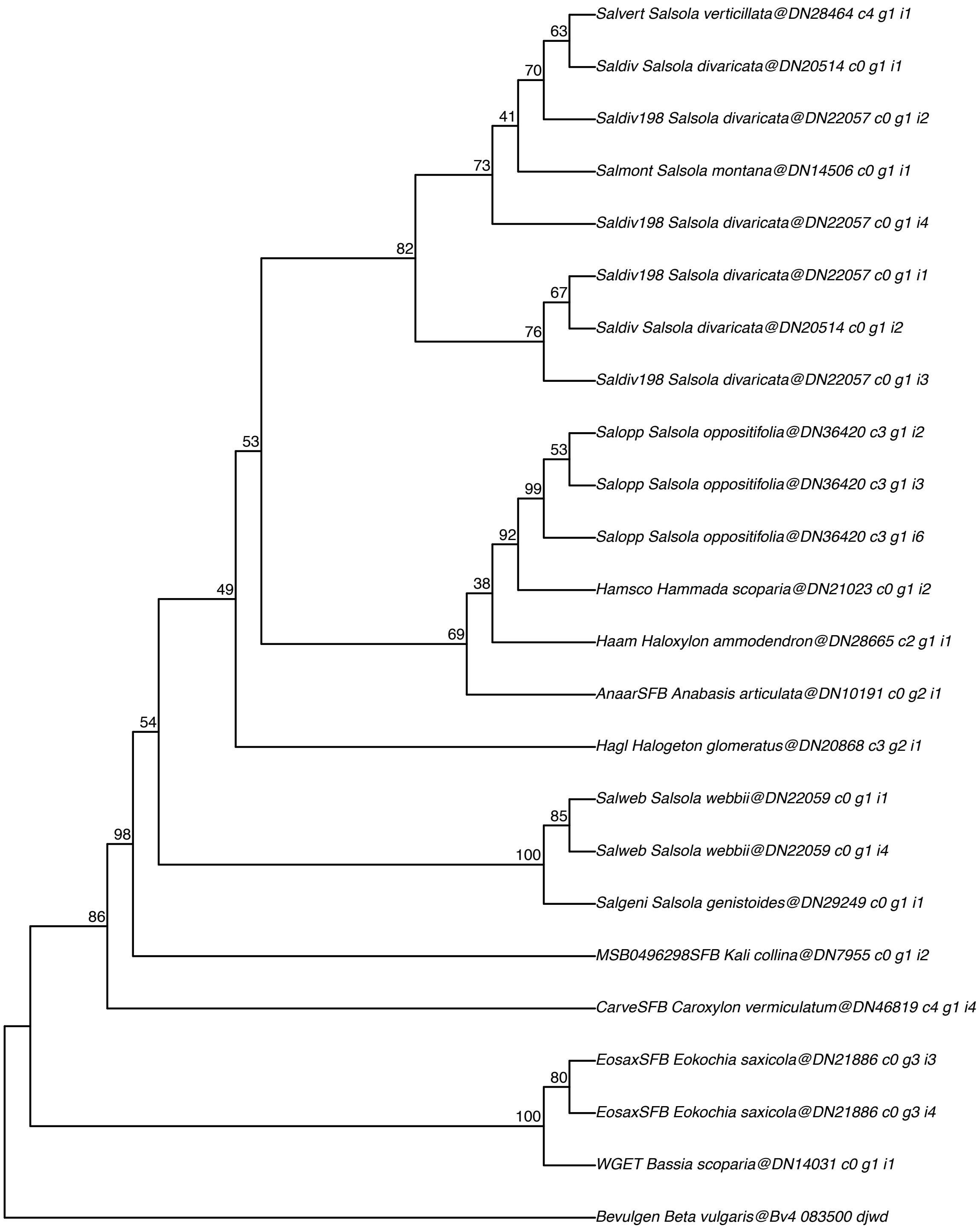

**PEP carboxylase–related protein (PEPC–RP)–AT3G42628**

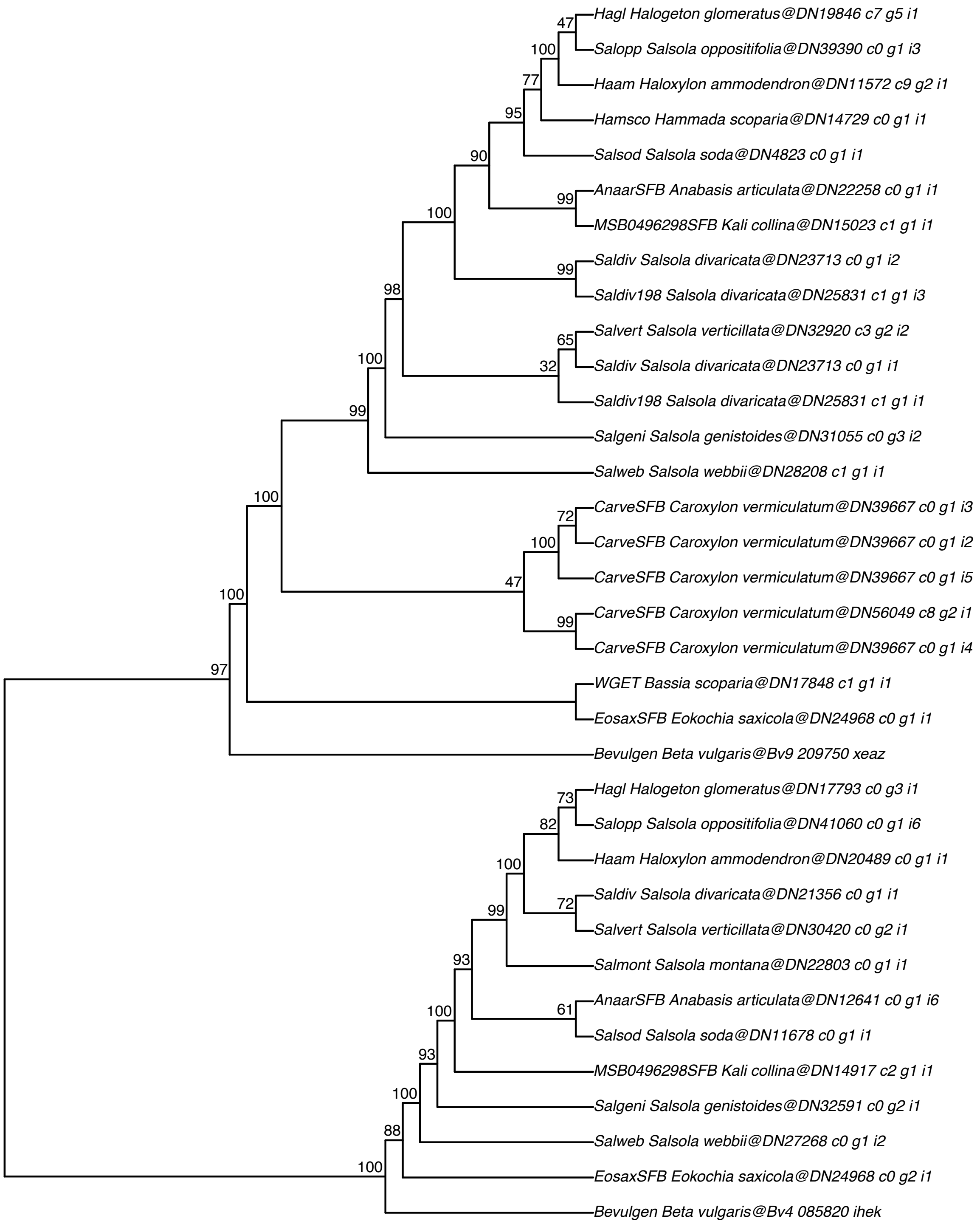

**Bile acid:sodium symporter family protein (BASS4)–AT3G56160**

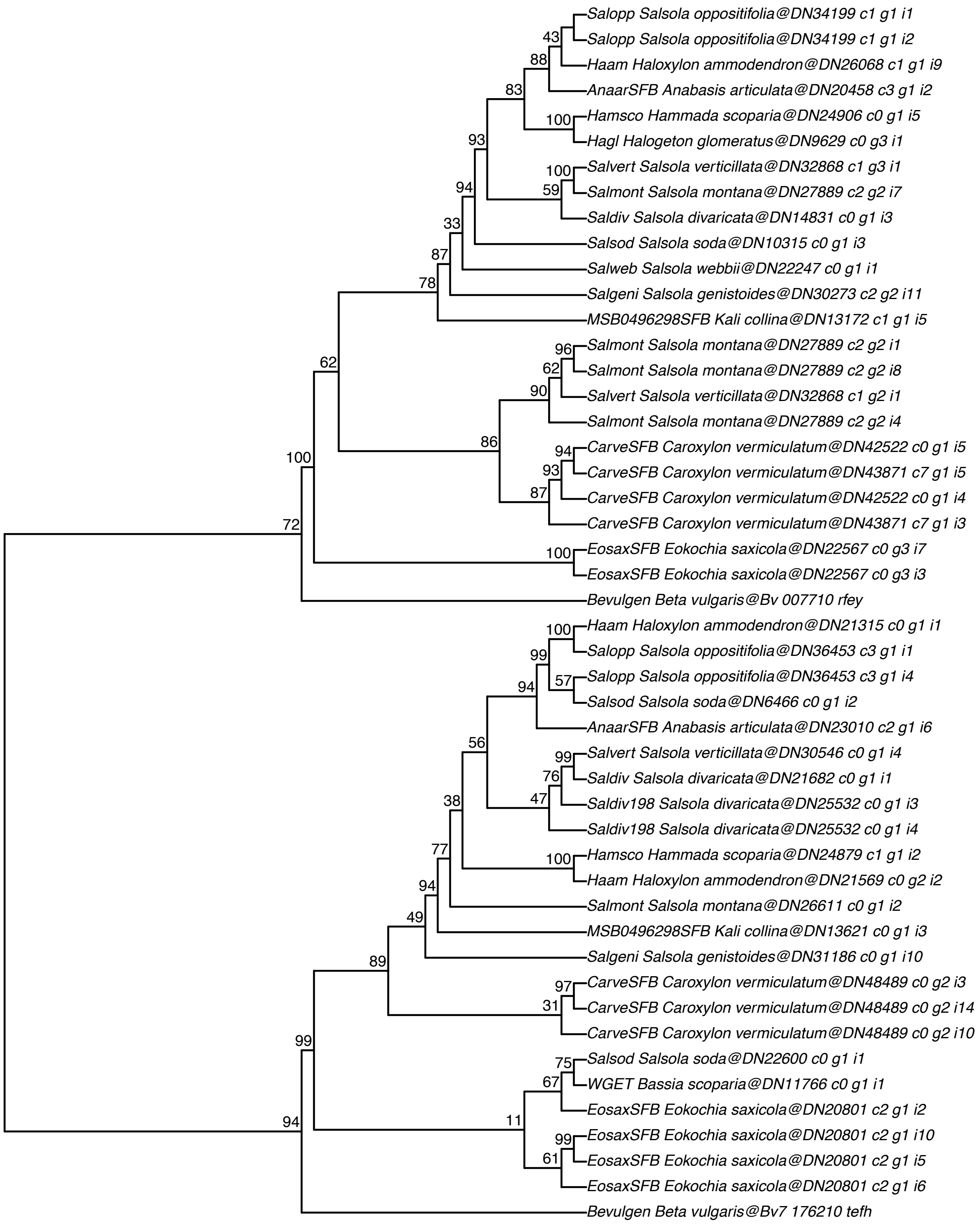

#### Pyruvate orthophosphat dikinase (PPdK)–AT4G15530

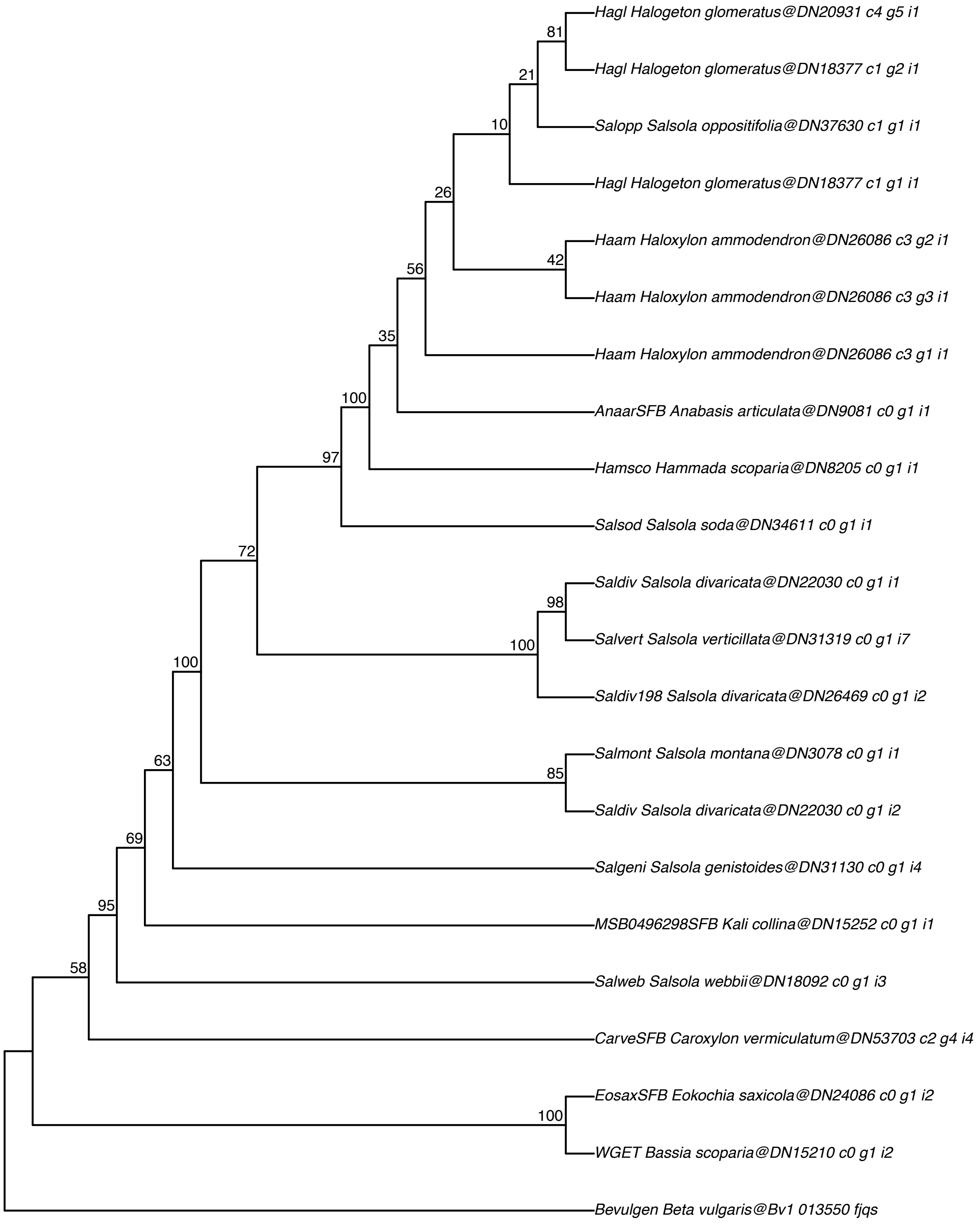

#### PPdK–related protein (PPdK–RP1)–AT4G21210

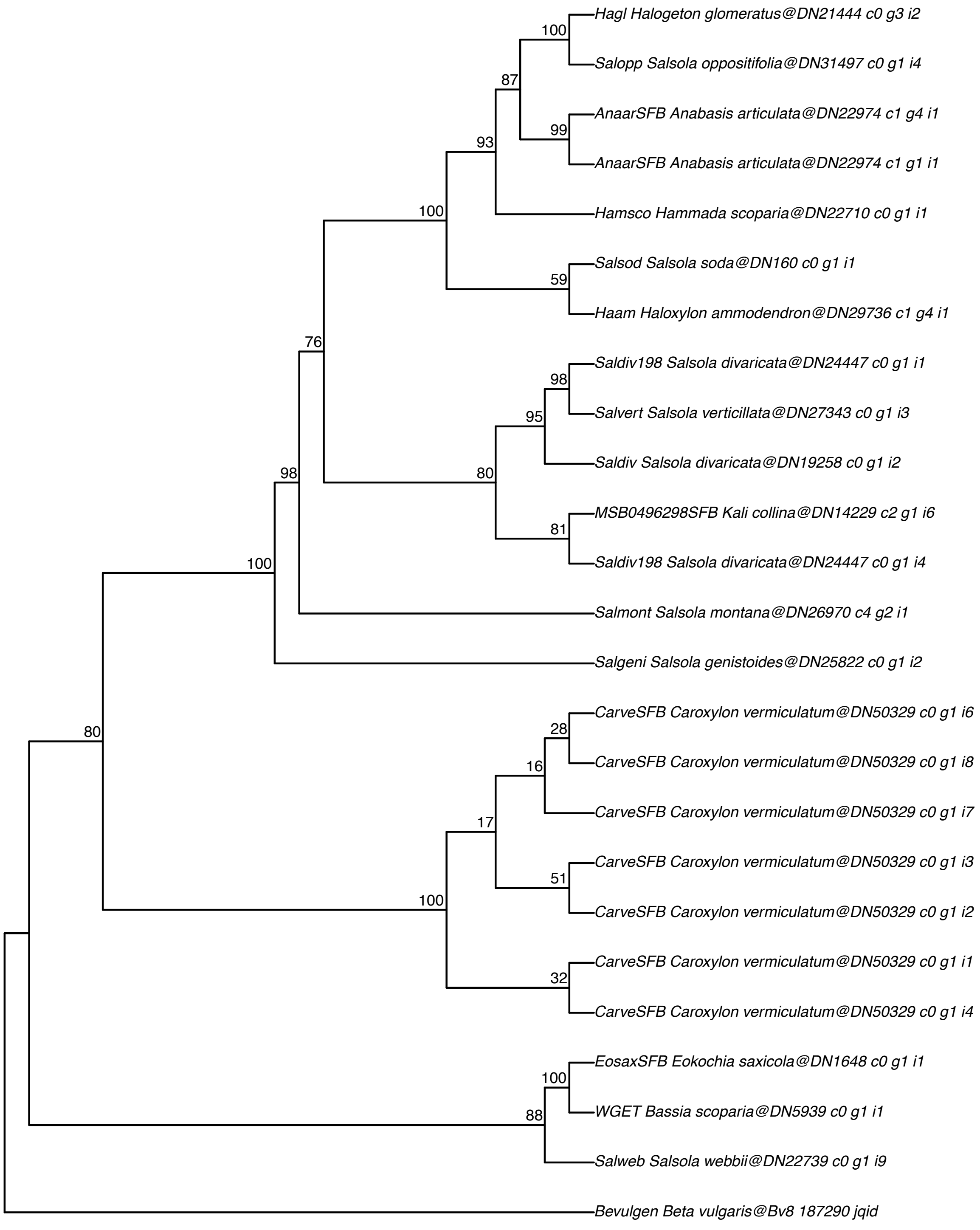

#### Aspartat aminotransferase (Asp–AT5)–AT4G31990

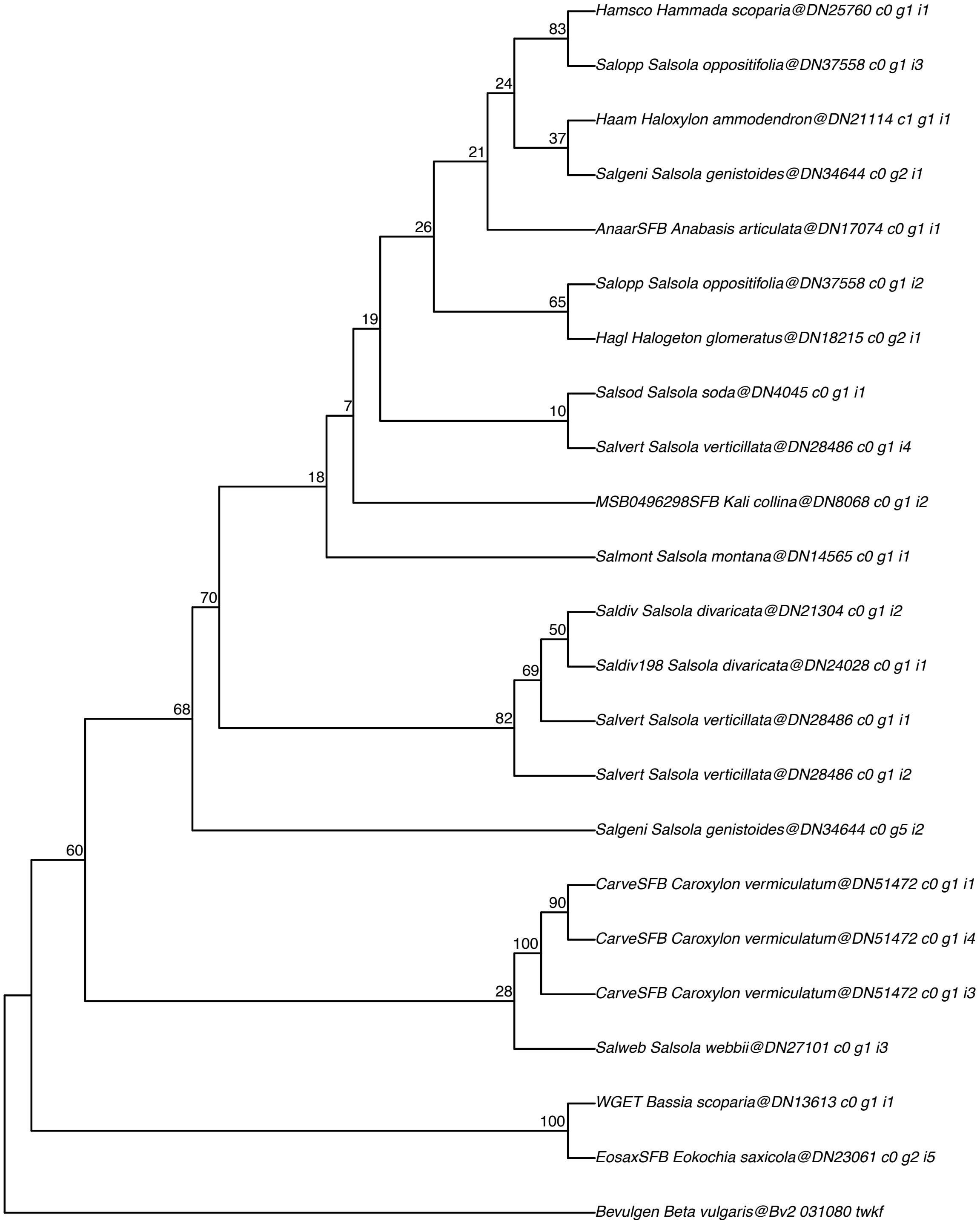

#### SHORTROOT–AT4G37650

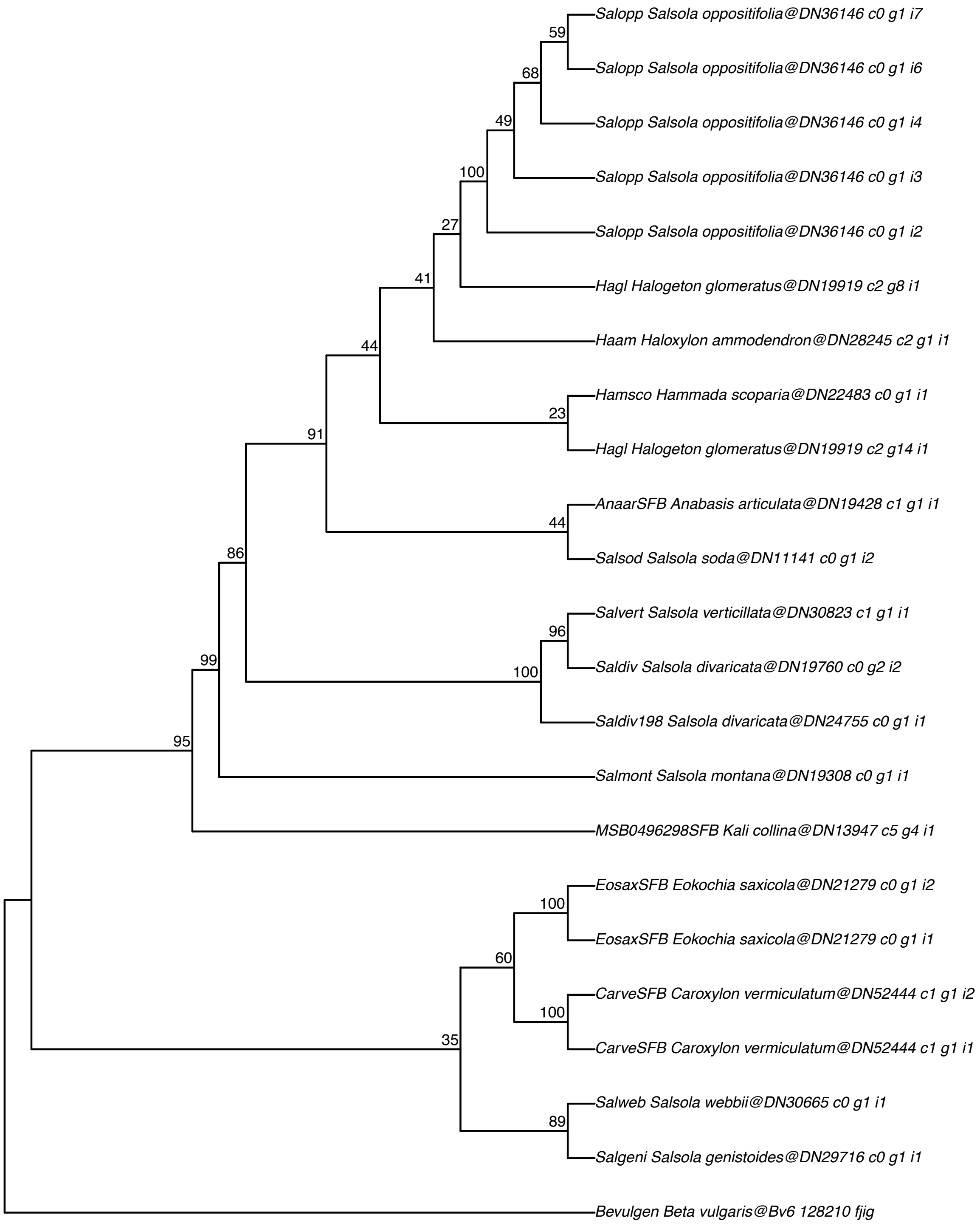

**Pyrophosphatase (PPase6)–AT5G09650**

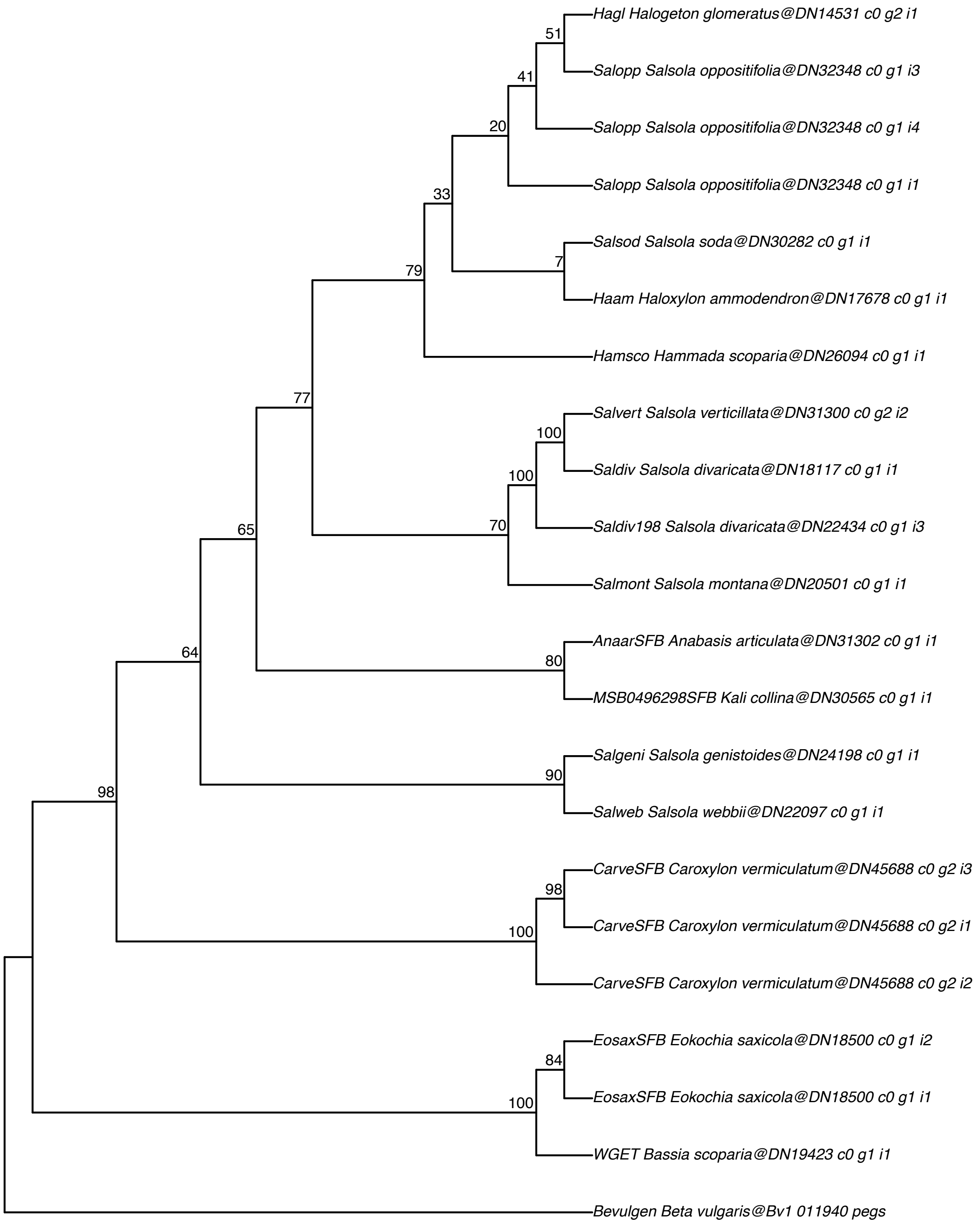

#### NADP–dependent malic enzyme (NADP–ME2)–AT5G11670

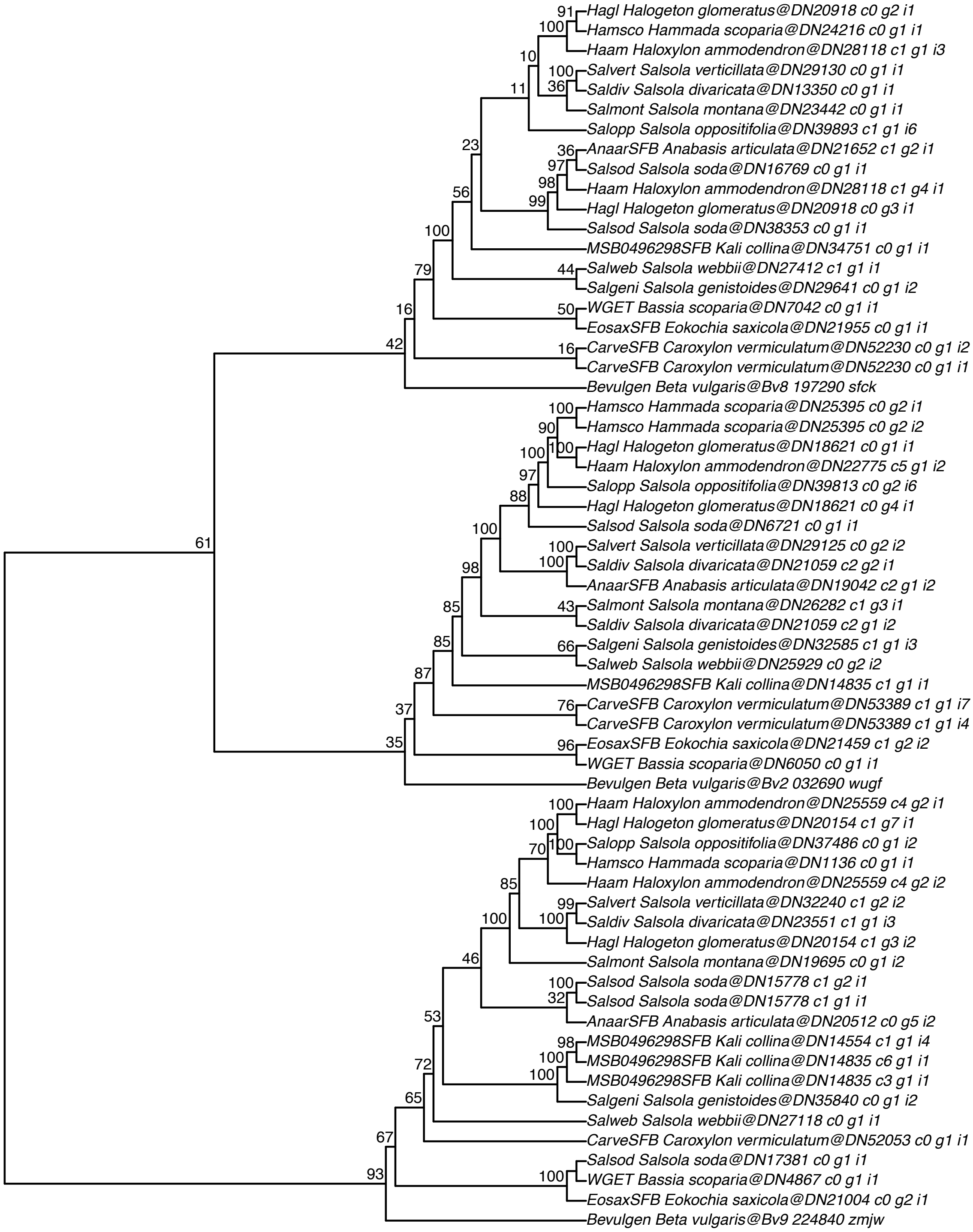

#### Dicarboxylate transporter (Dit1)–AT5G12860

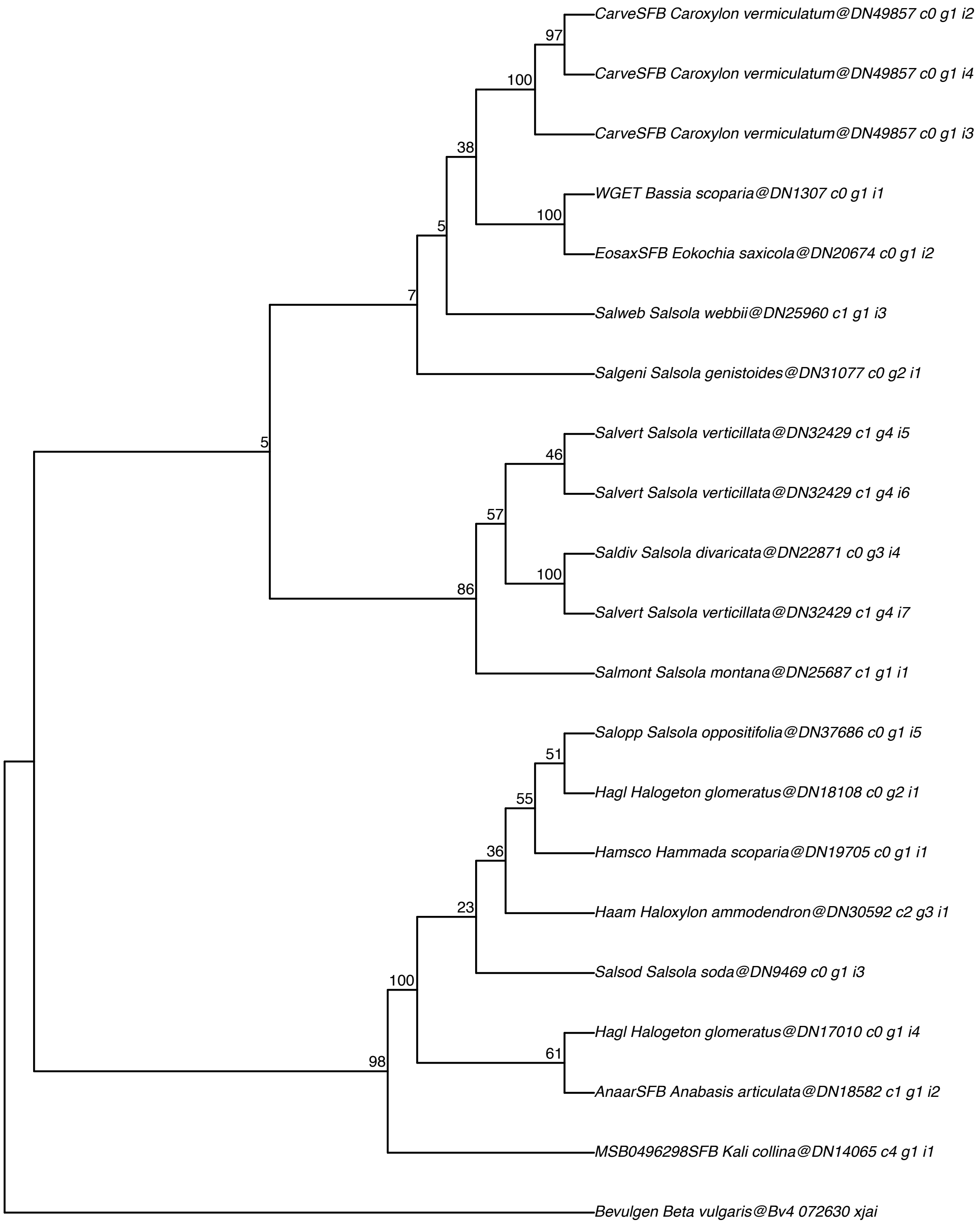

#### PEP/phosphate translocator (PPT1/CUE1)−AT5G33320

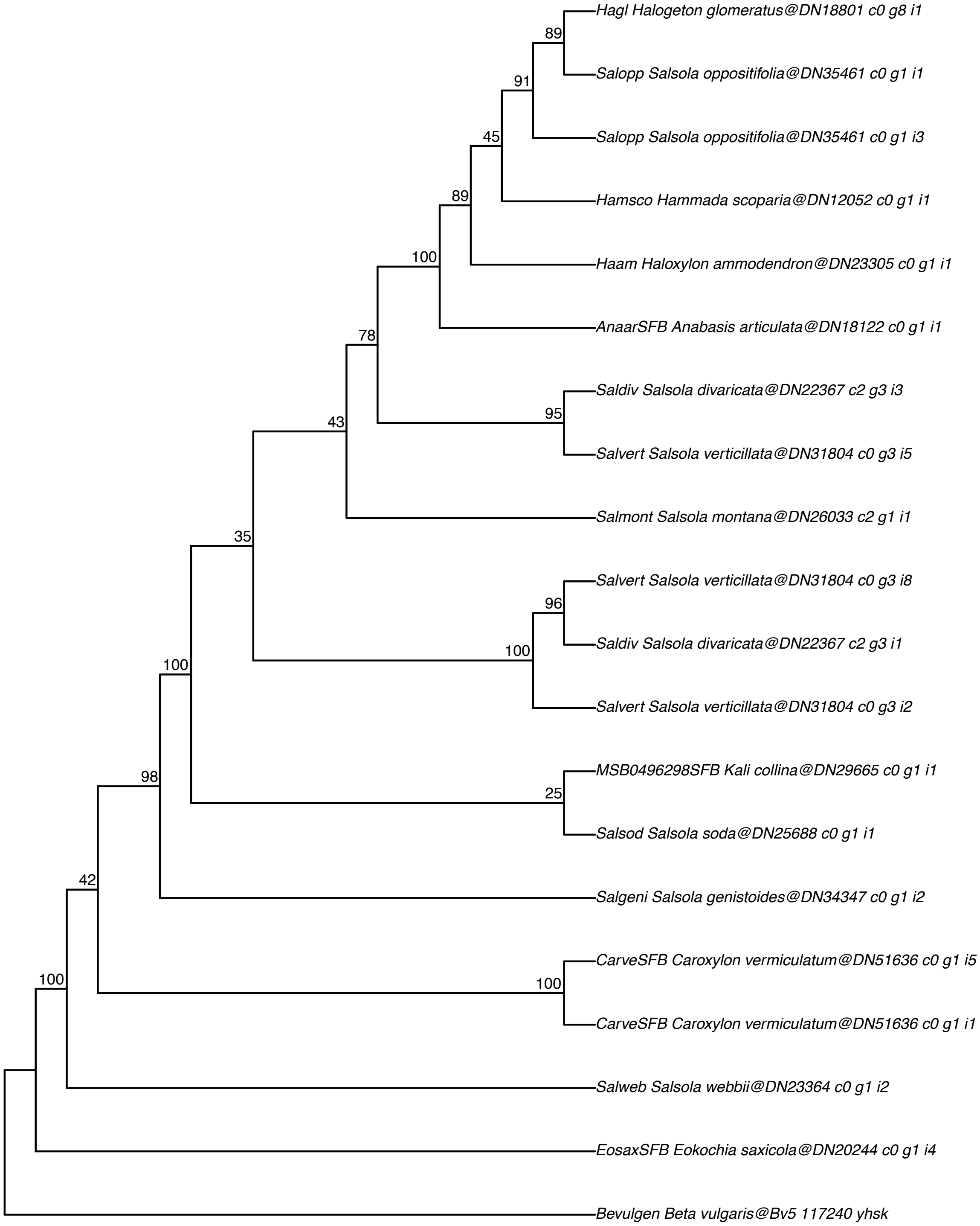

#### Triose phosphate translocator (TPT)–AT5G46110

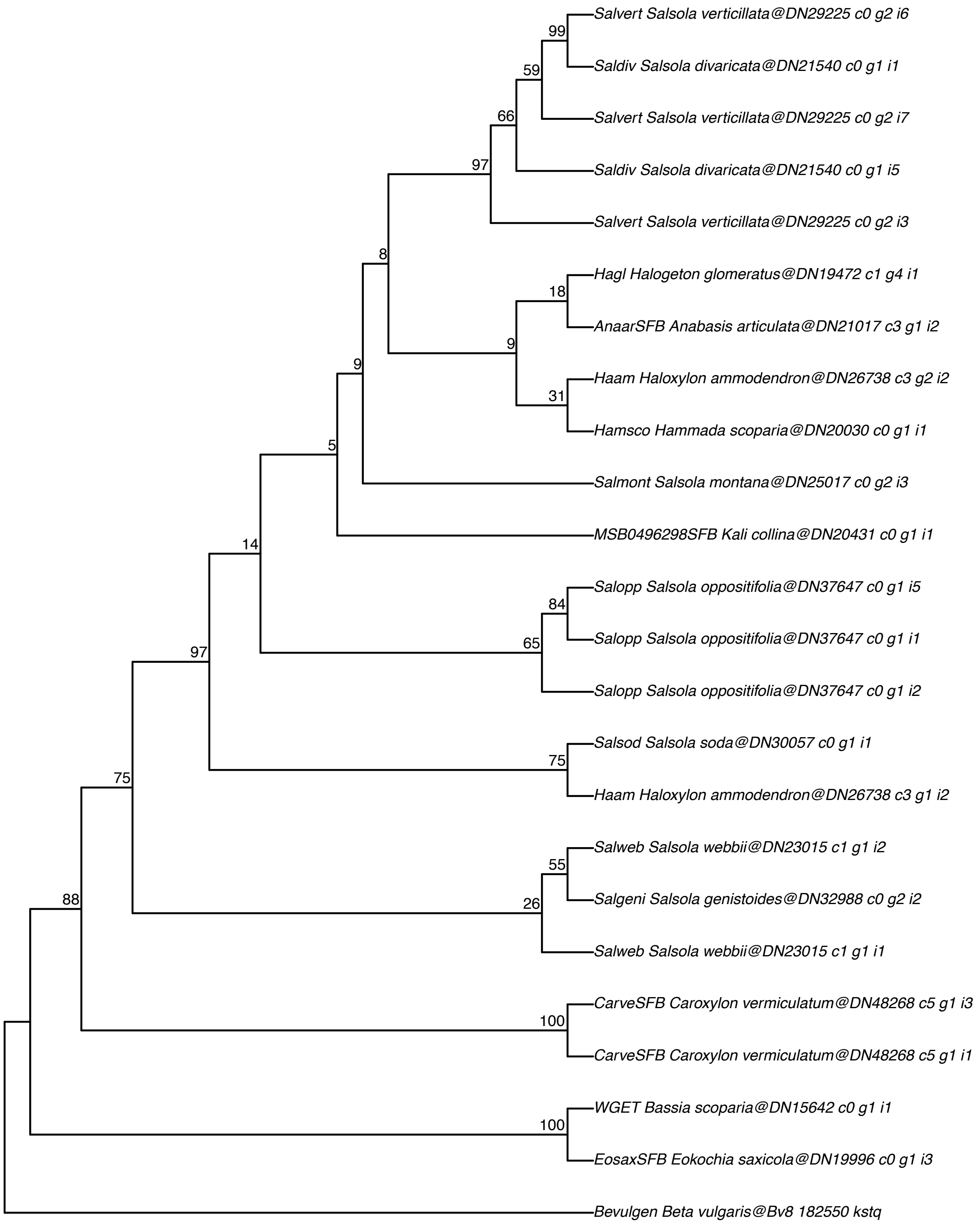

#### Adenosine monophosphate kinase (AMK2)–AT5G47840

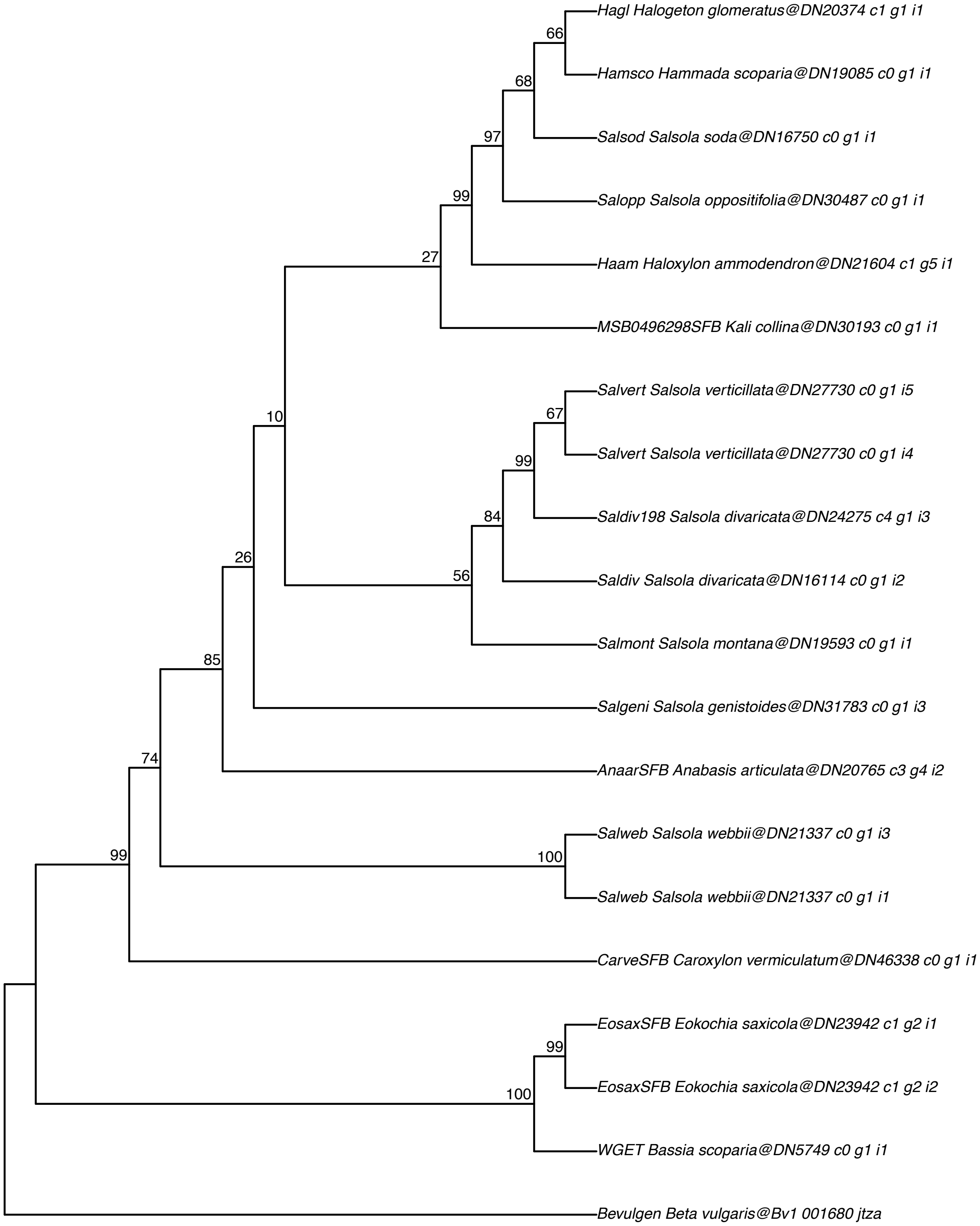

#### NADP–dependent malate dehydrogenase (NADP–MDH)–AT5G58330

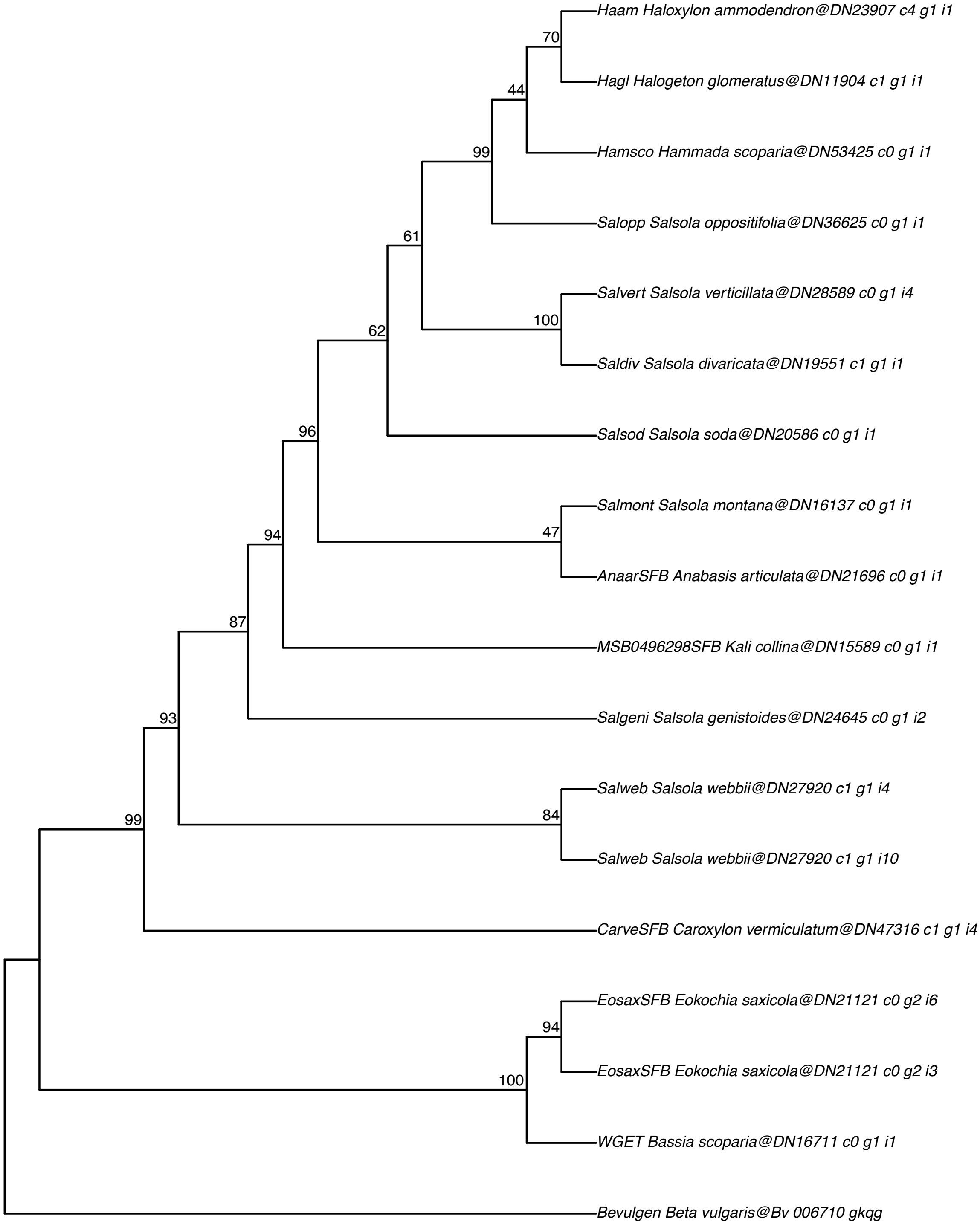

#### Dicarboxylate transporter (Dit2)–AT5G64280

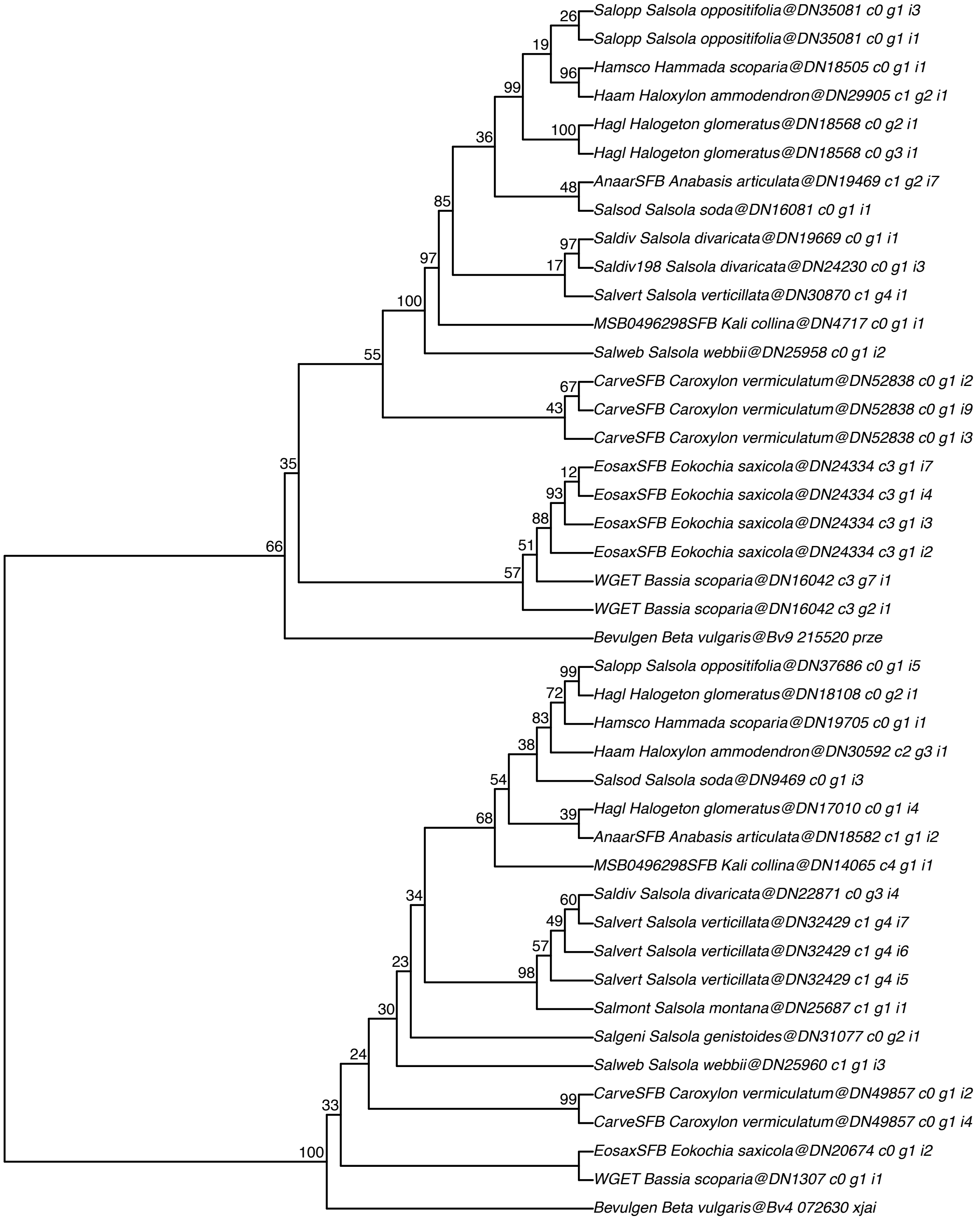

### Isocitrate lyase–AT3G21720

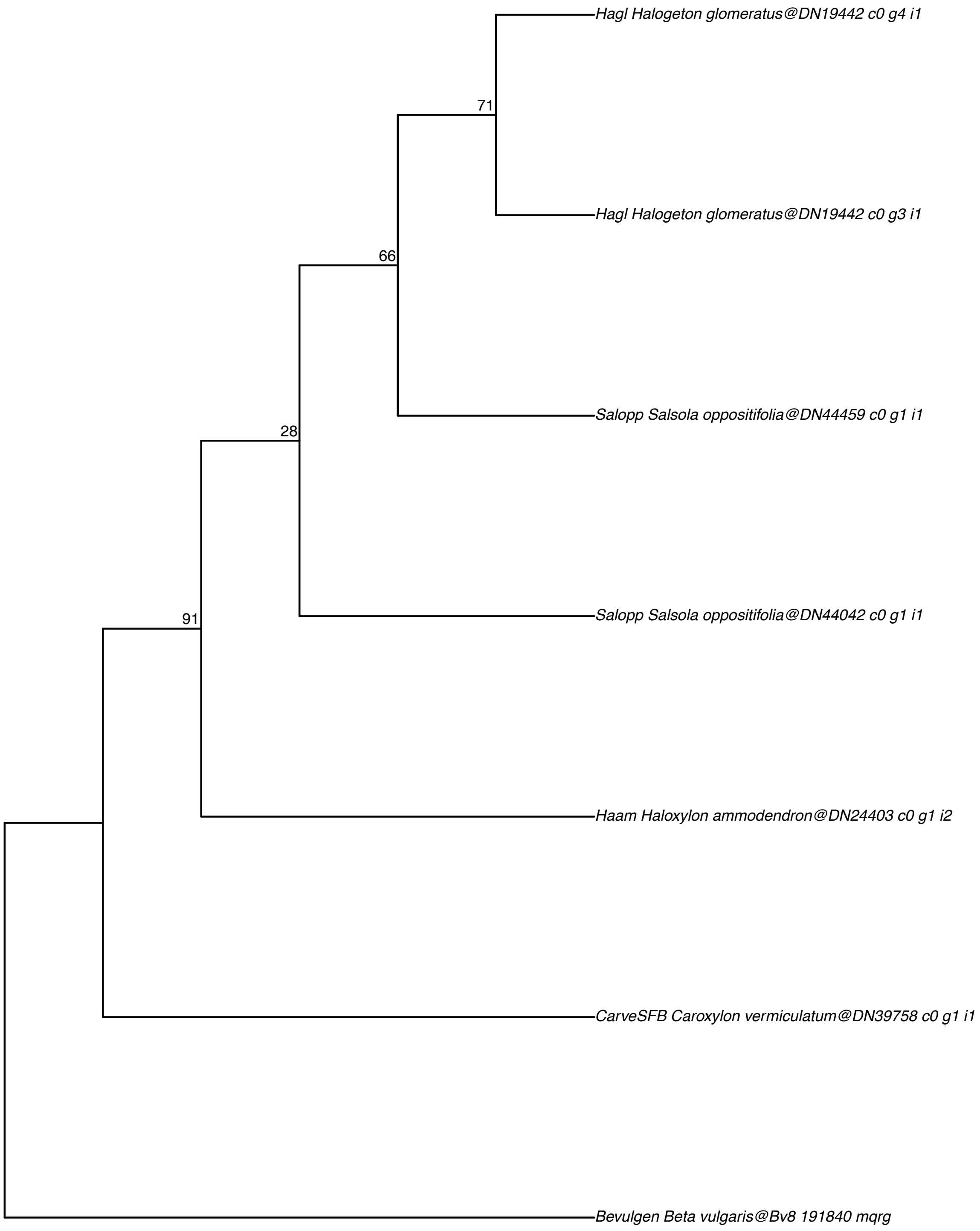

### PEP carboxykinase (PEP−CK1)−AT4G37870

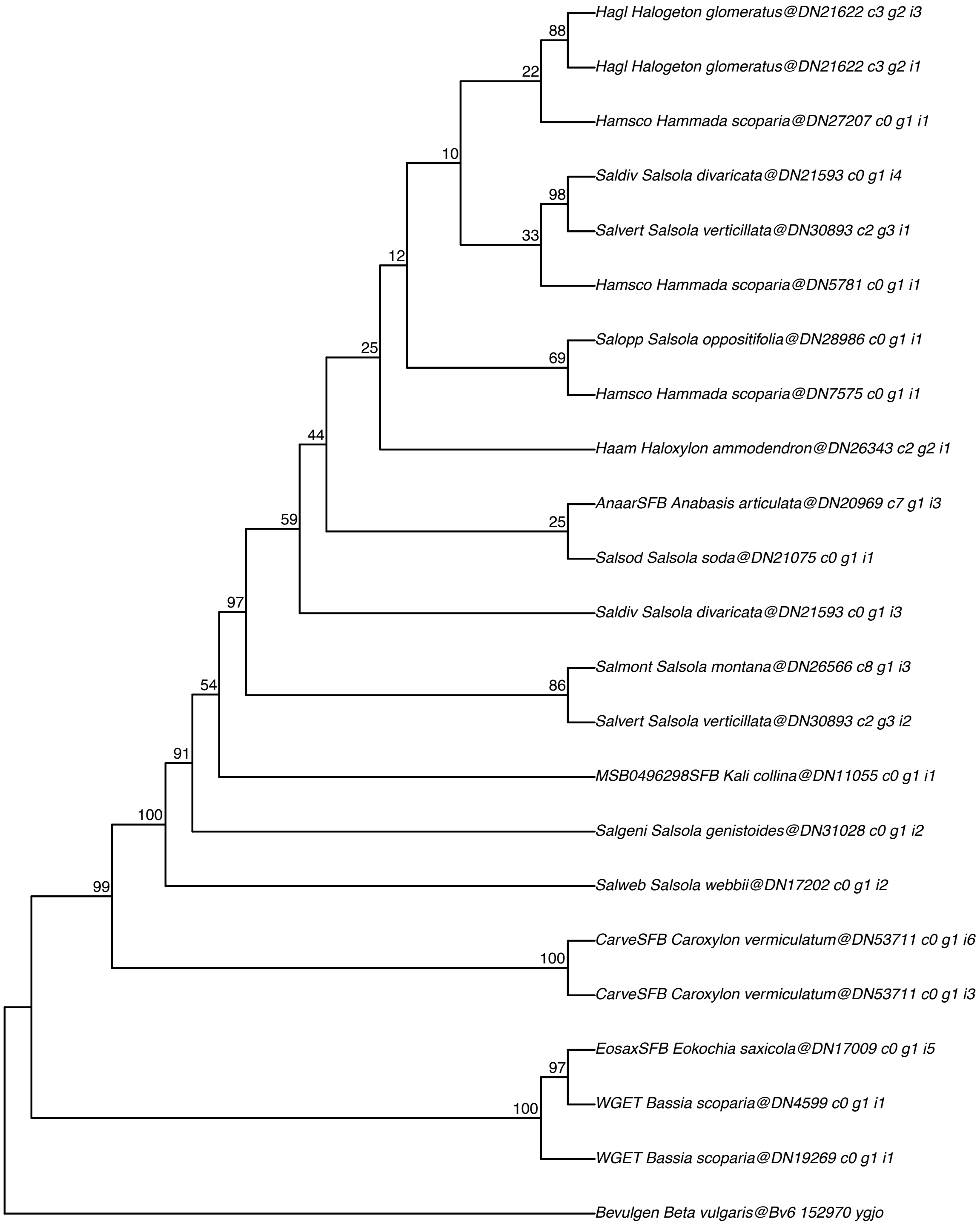

#### Malate synthase–AT5G03860

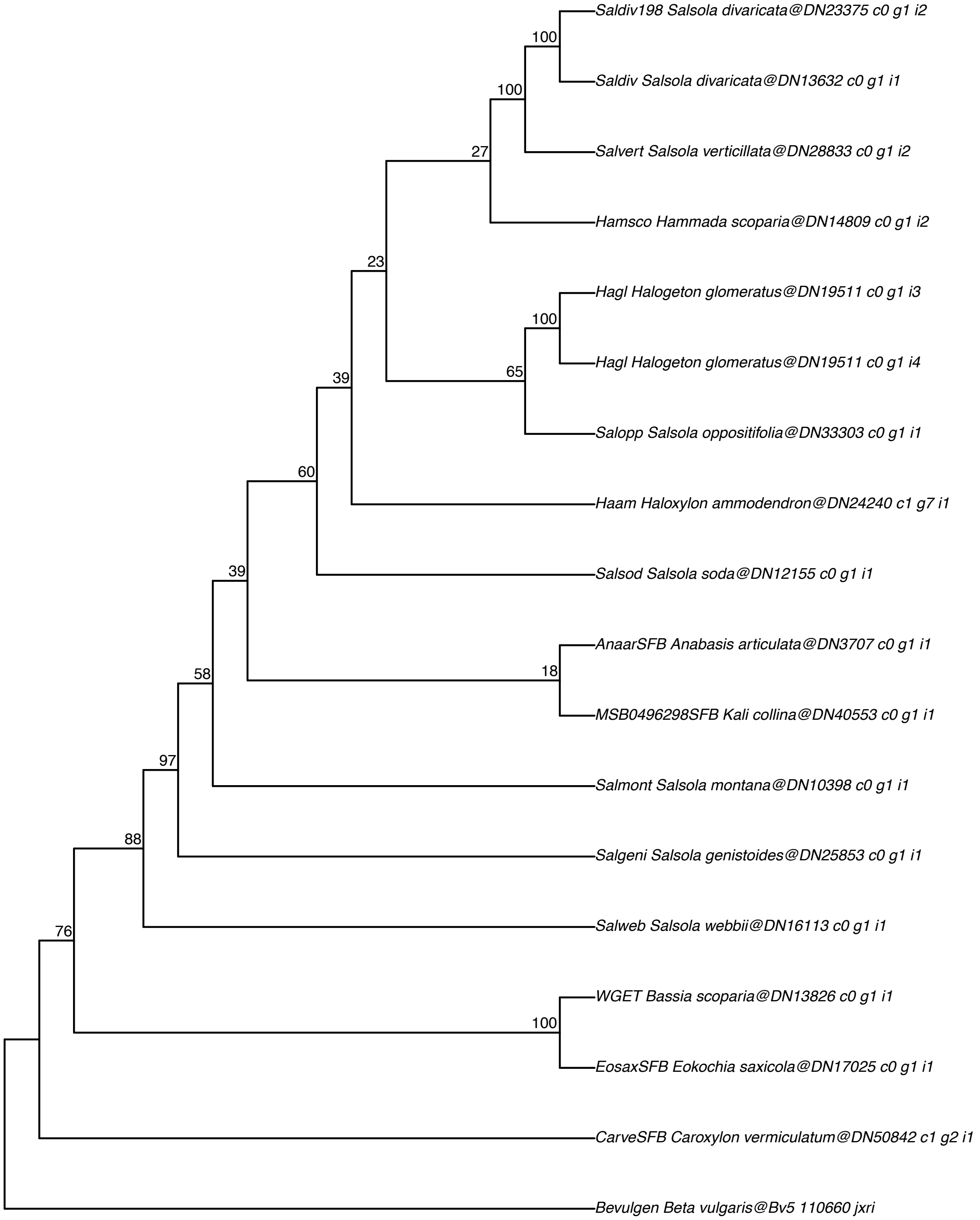

#### PEP carboxykinase (PEP−CK2)−AT5G65690

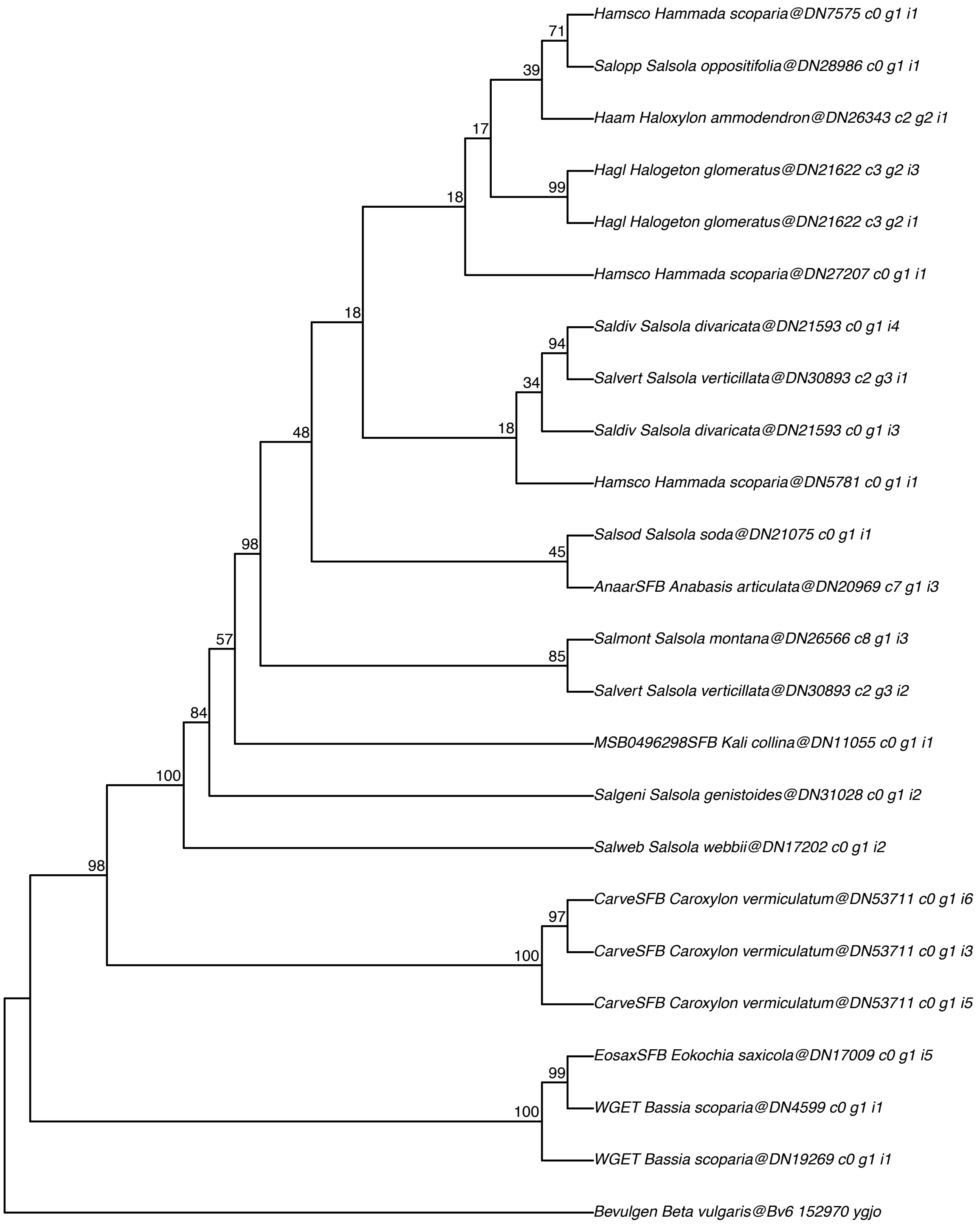

#### Glycine decarboxylase T−protein (GDC T−protein)−AT1G11860

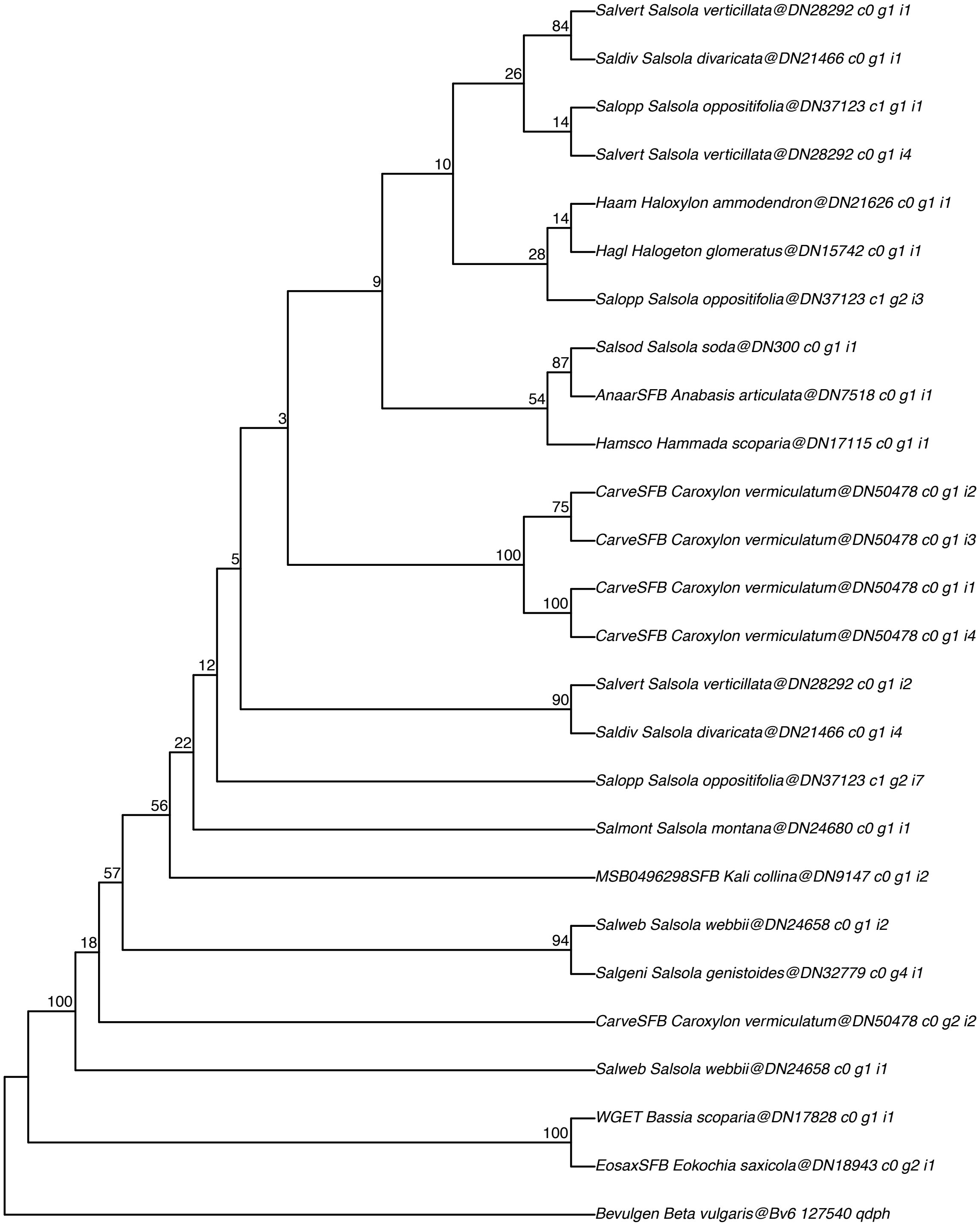

#### Glutamate:glyoxylate aminotransferase (GGT1)–AT1G23310

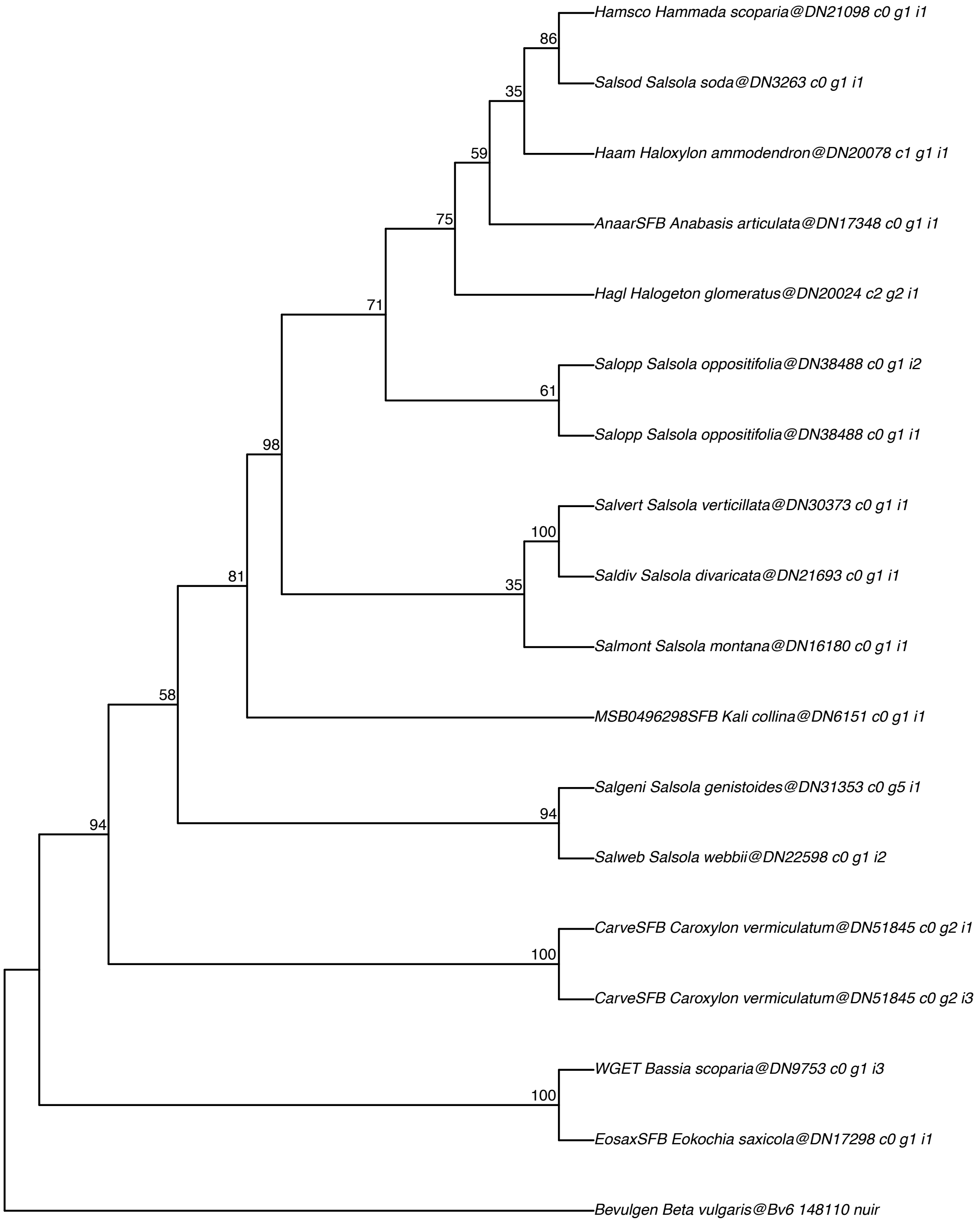

#### Glycine cleavage H−protein (GDC H−protein)−AT1G32470

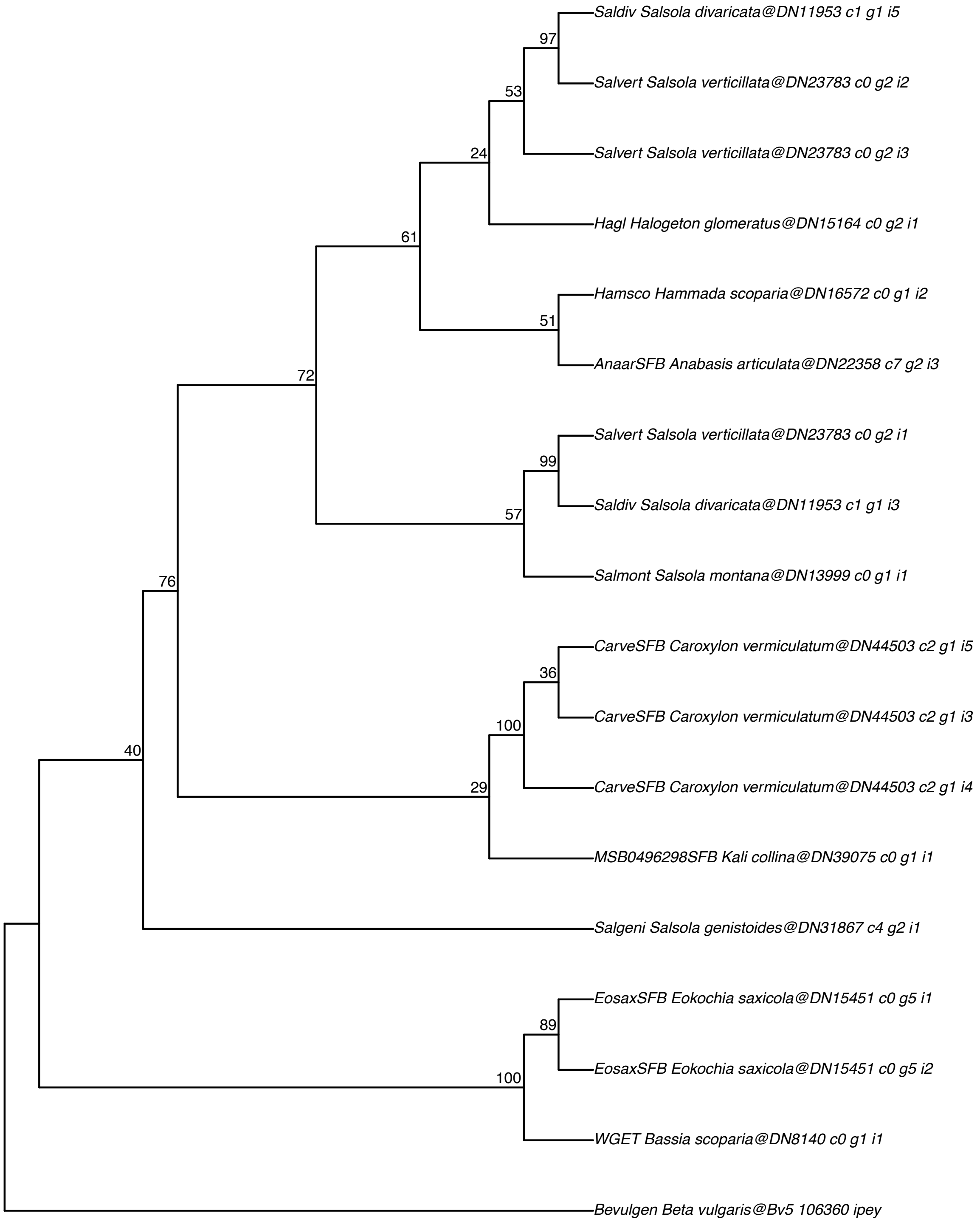

#### Glycine decarboxylase L–protein (GDC L–protein)–AT1G48030

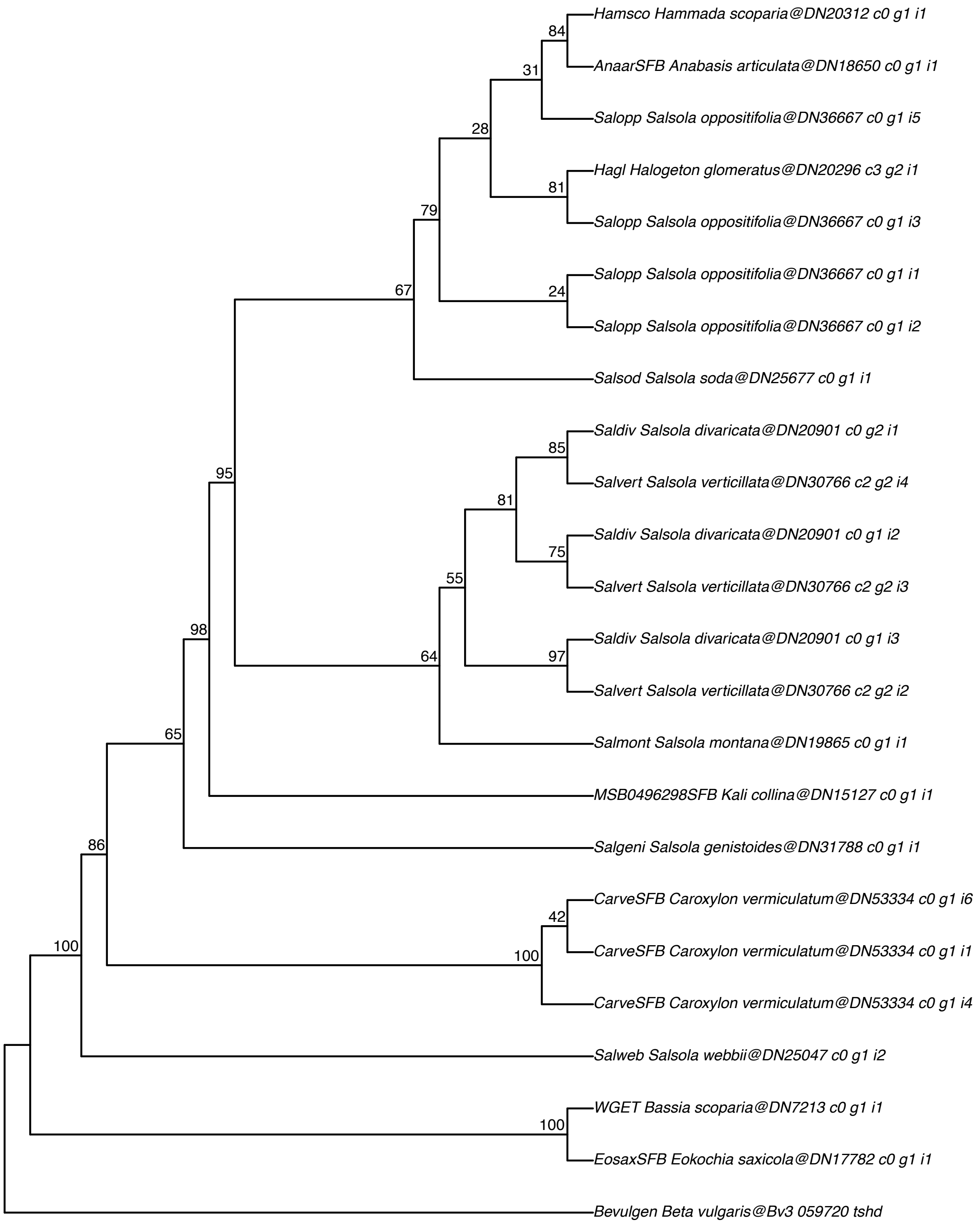

#### NADH–dependent hydroxypyruvate reductase (HPR1)–AT1G68010

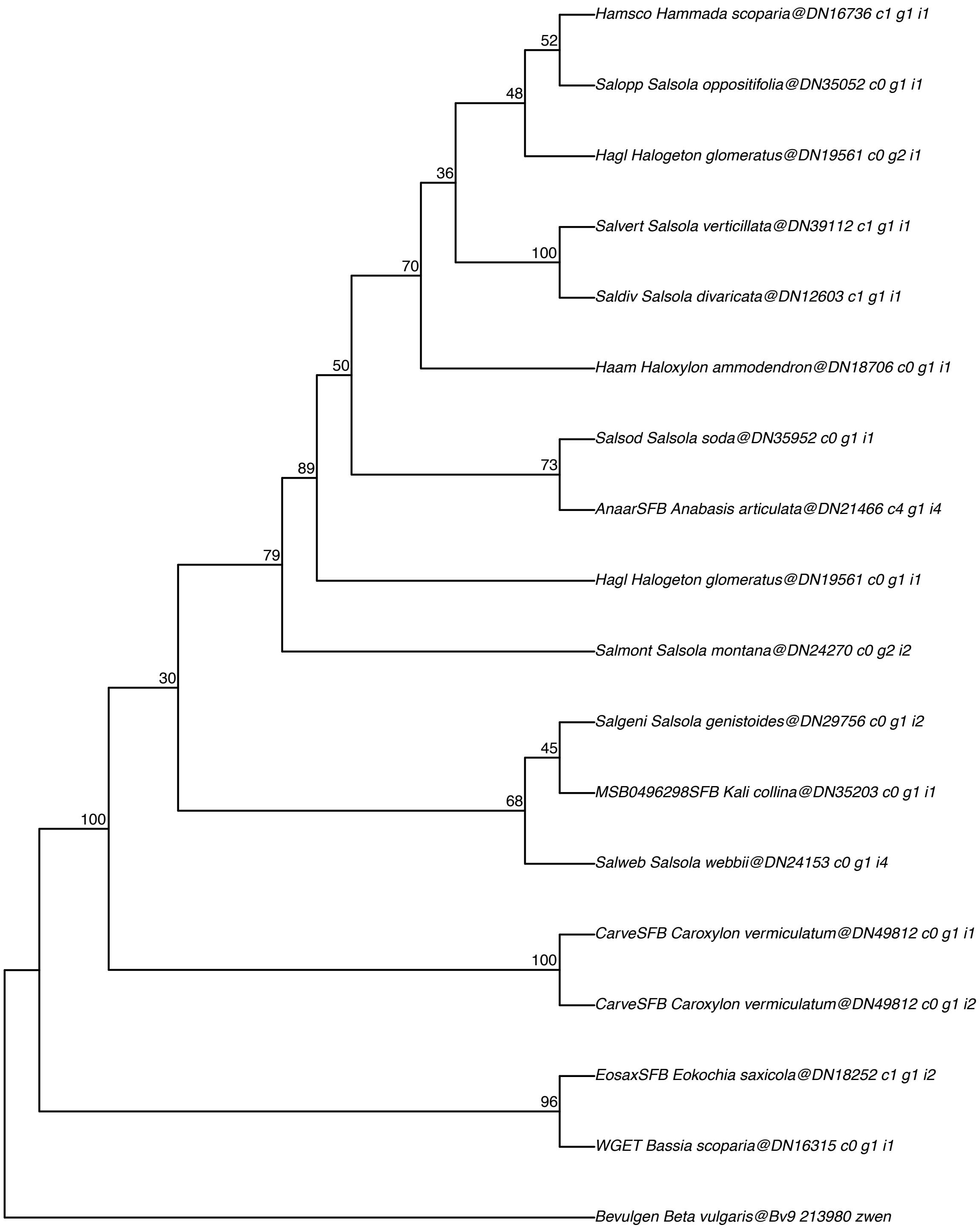

#### Glycerate 3–kinase (GLYK)–AT1G80380

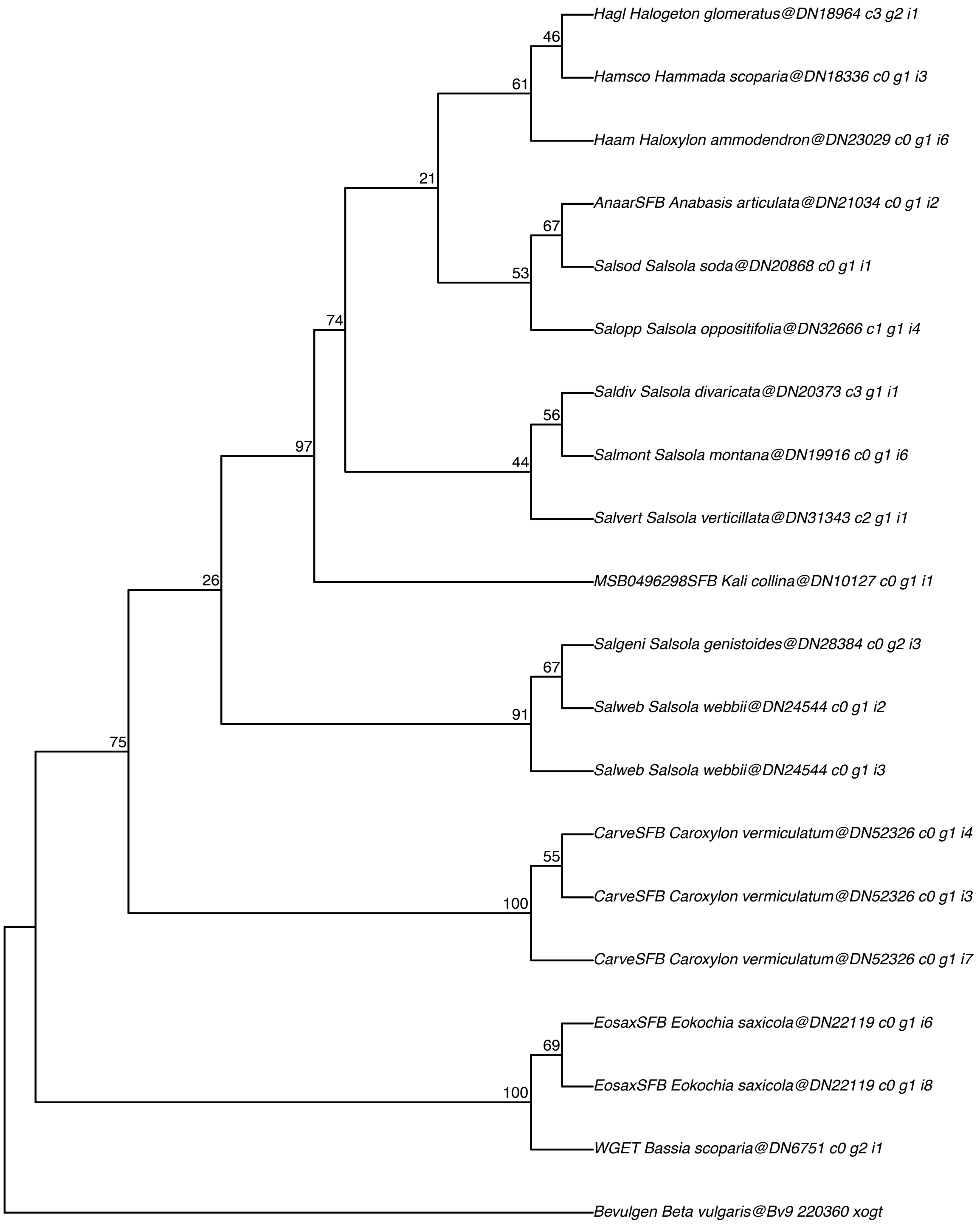

#### Serine:glyoxylate aminotransferase (AGT1/SGT1)–AT2G13360

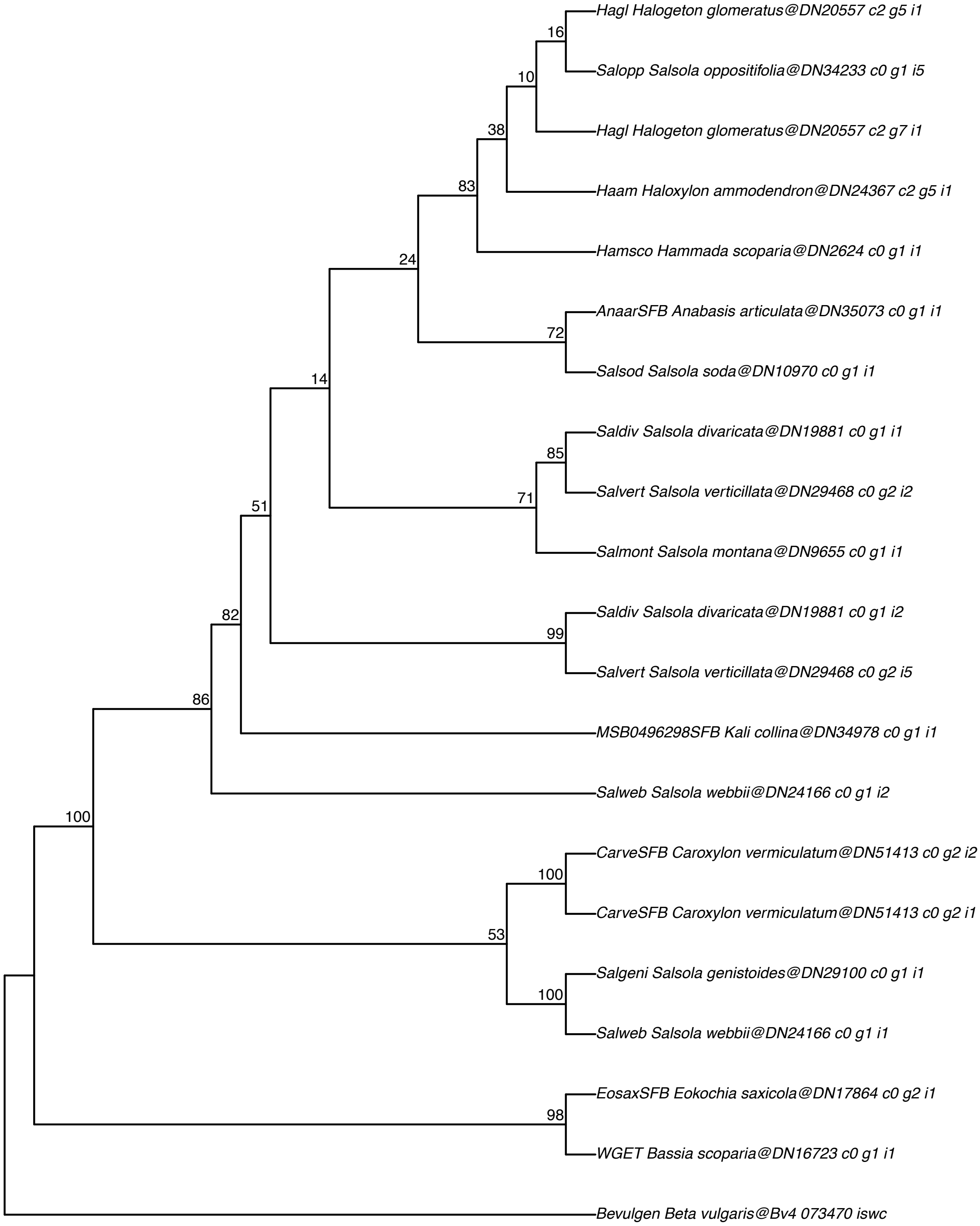

#### Glycine decarboxylase P–protein (GDC P–protein)–AT2G26080

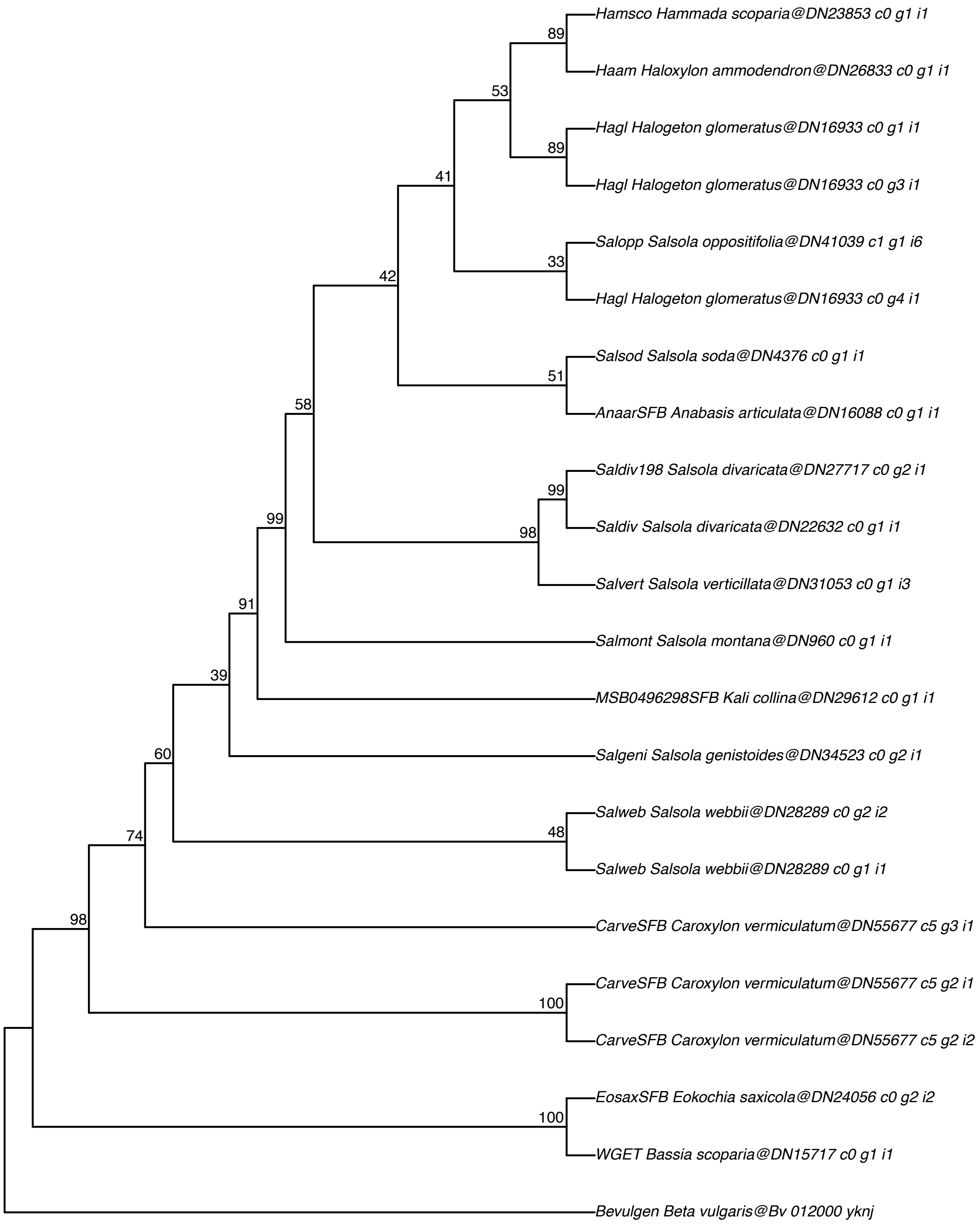

#### Glycine cleavage H–protein (GDC H–protein)–AT2G35120

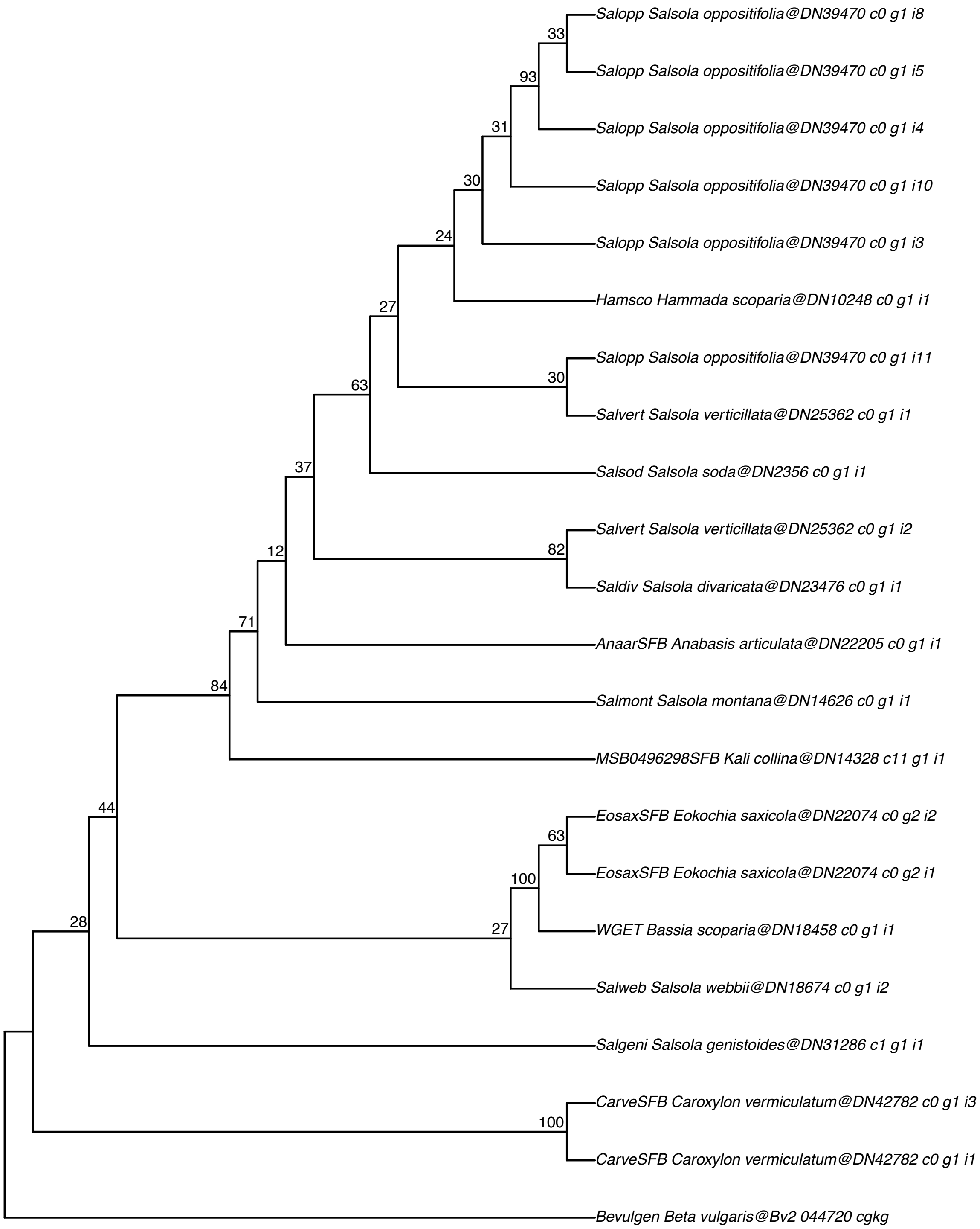

#### Serine:glyoxylate aminotransferase (AGT3/SGT3)–AT2G38400

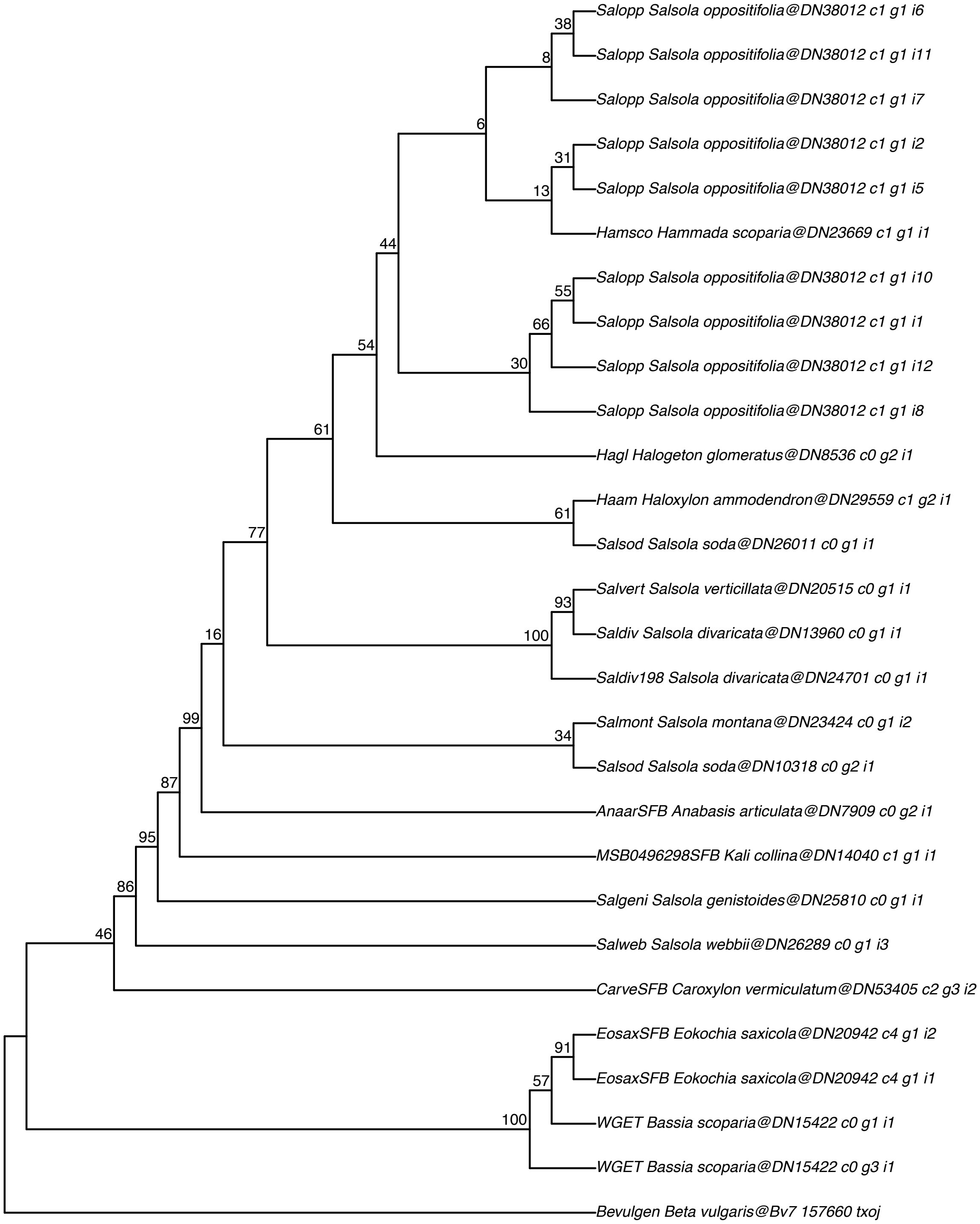

#### Serine:glyoxylate aminotransferase (AGT/SGT)–AT3G08860

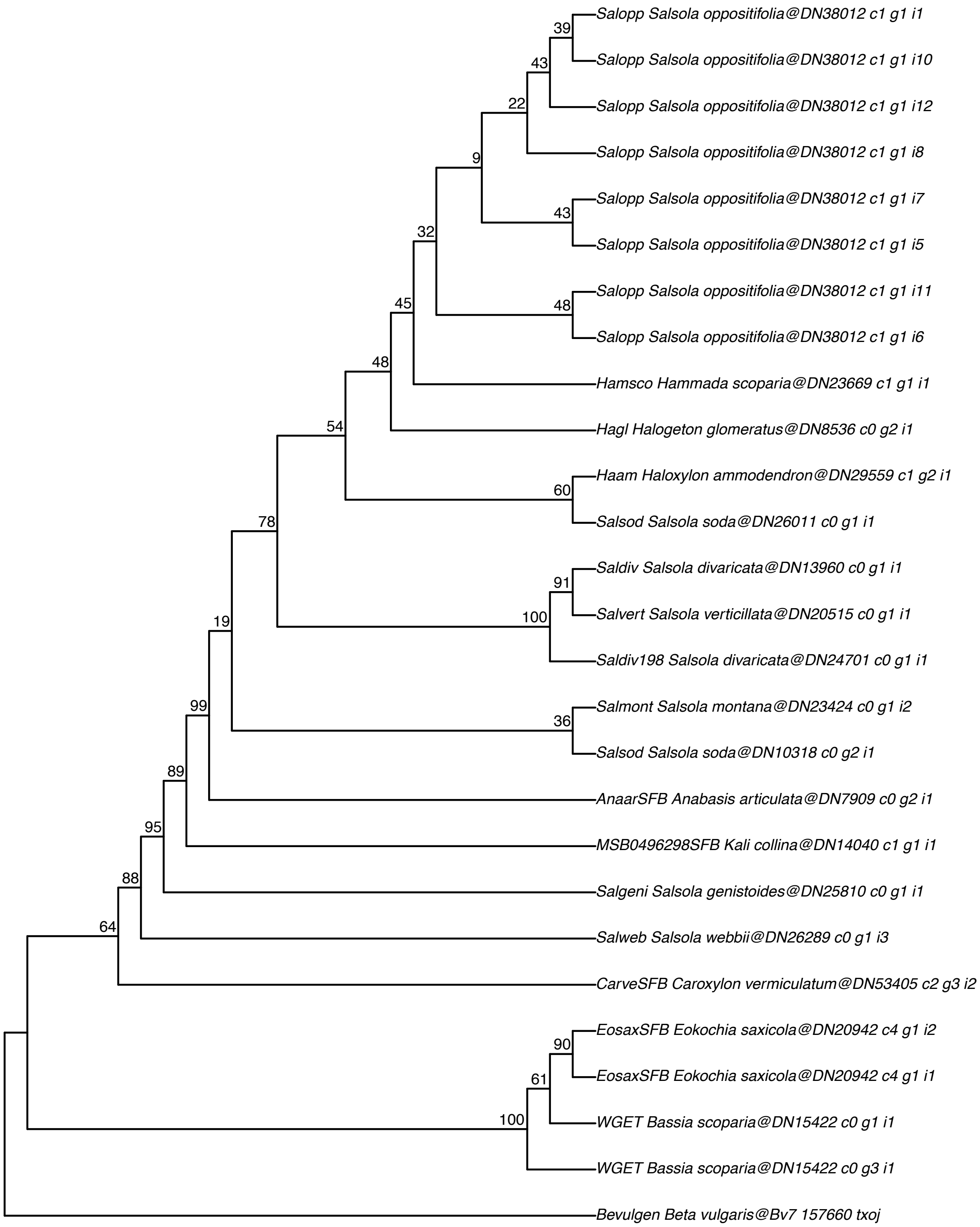

#### Glycolate oxidase (GOX2)–AT3G14415

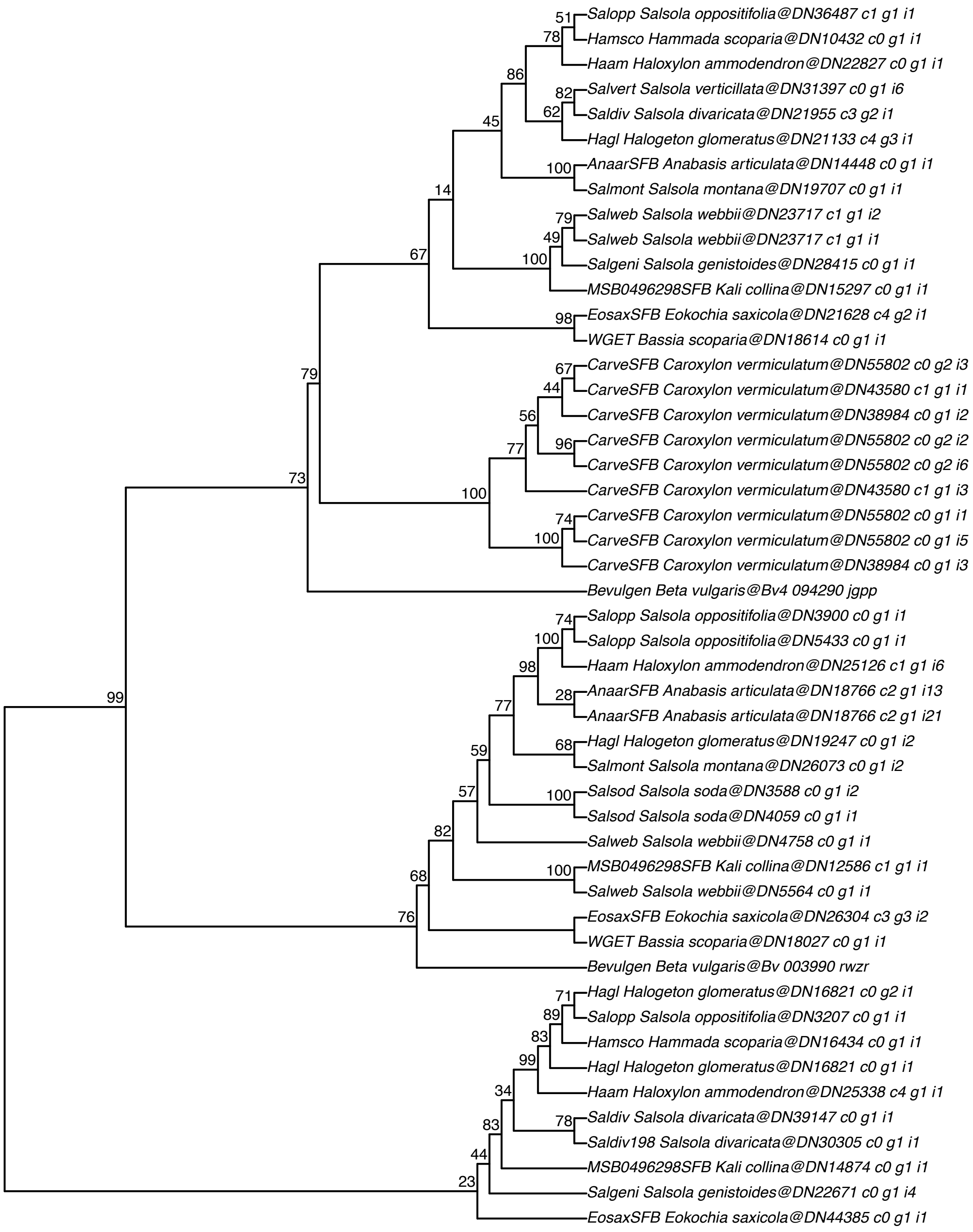

#### Glycolate oxidase (GOX1)–AT3G14420

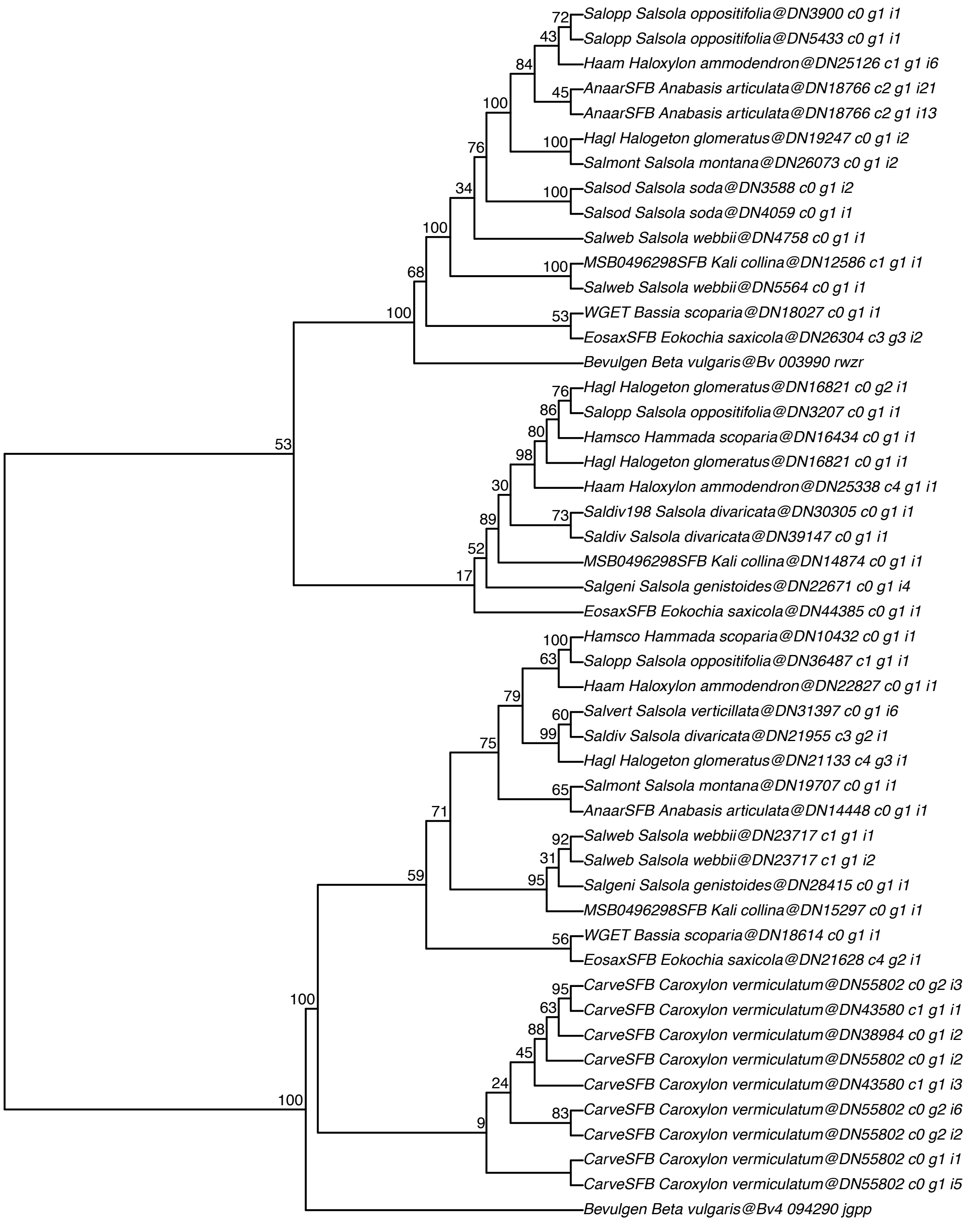

#### Glycolate oxidase (GOX3)–AT4G18360

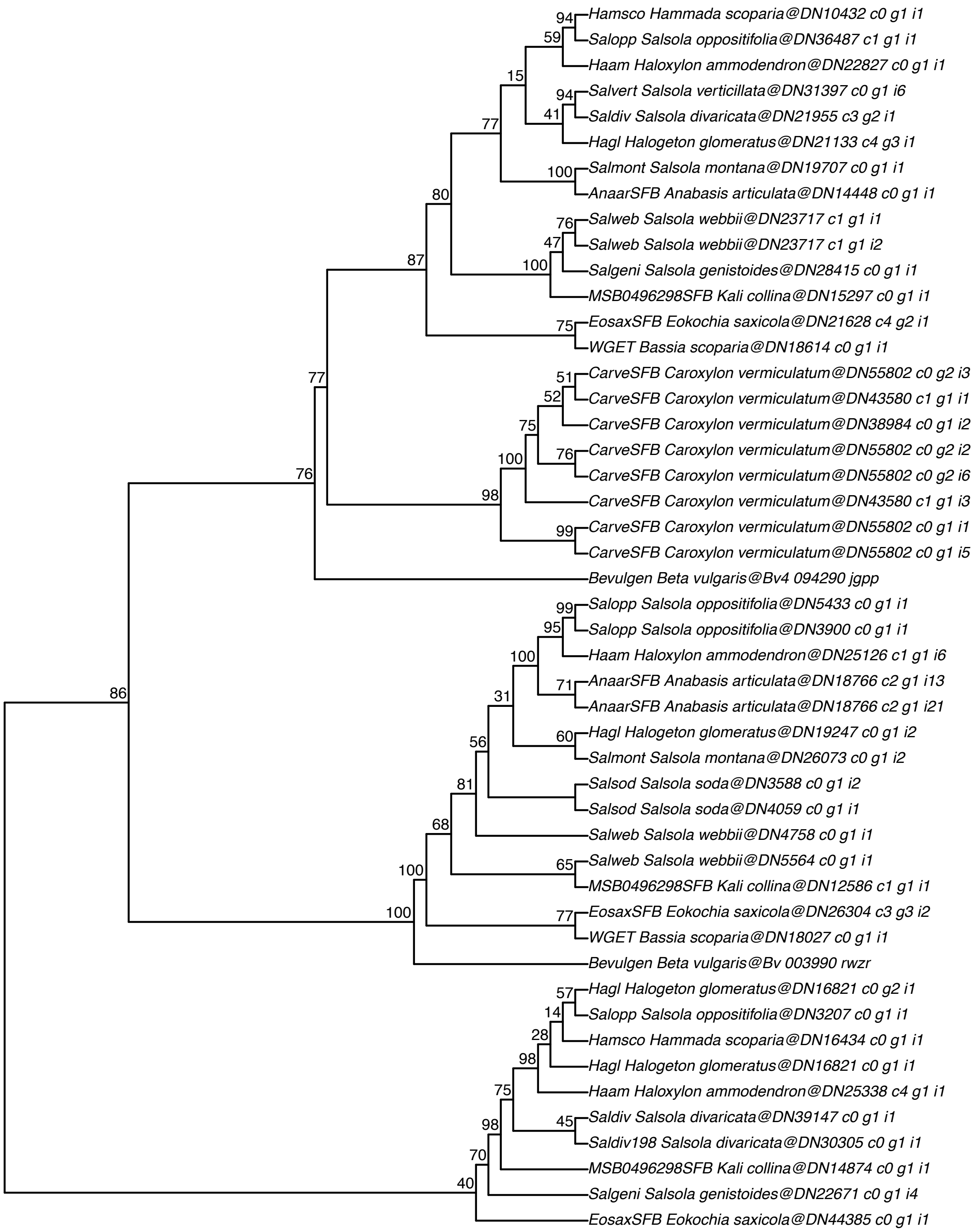

#### Serine hydroxymethyltransferase (SHMT1)–AT4G37930

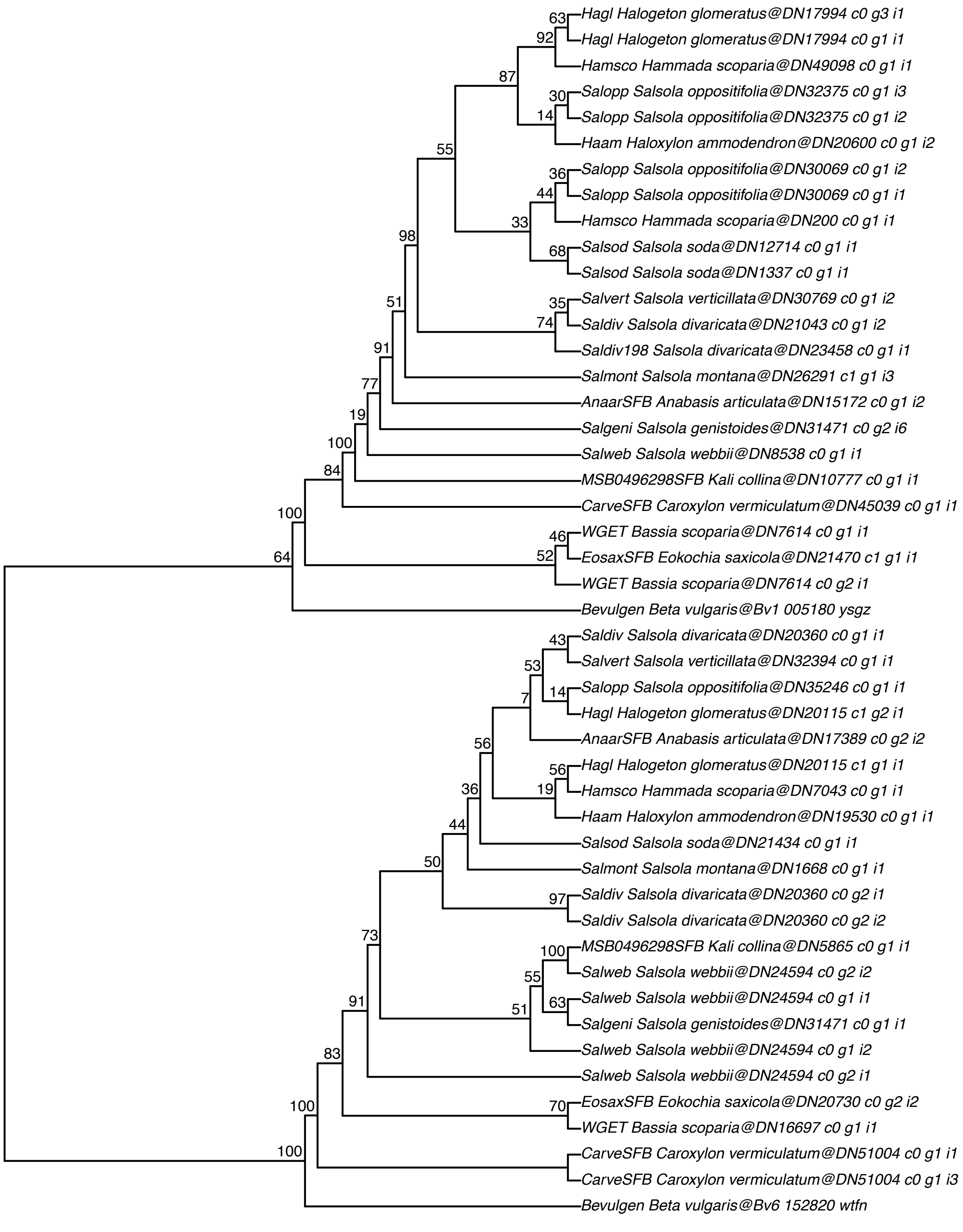

#### Serine:glyoxylate aminotransferase (AGT2/SGT2)–AT4G39660

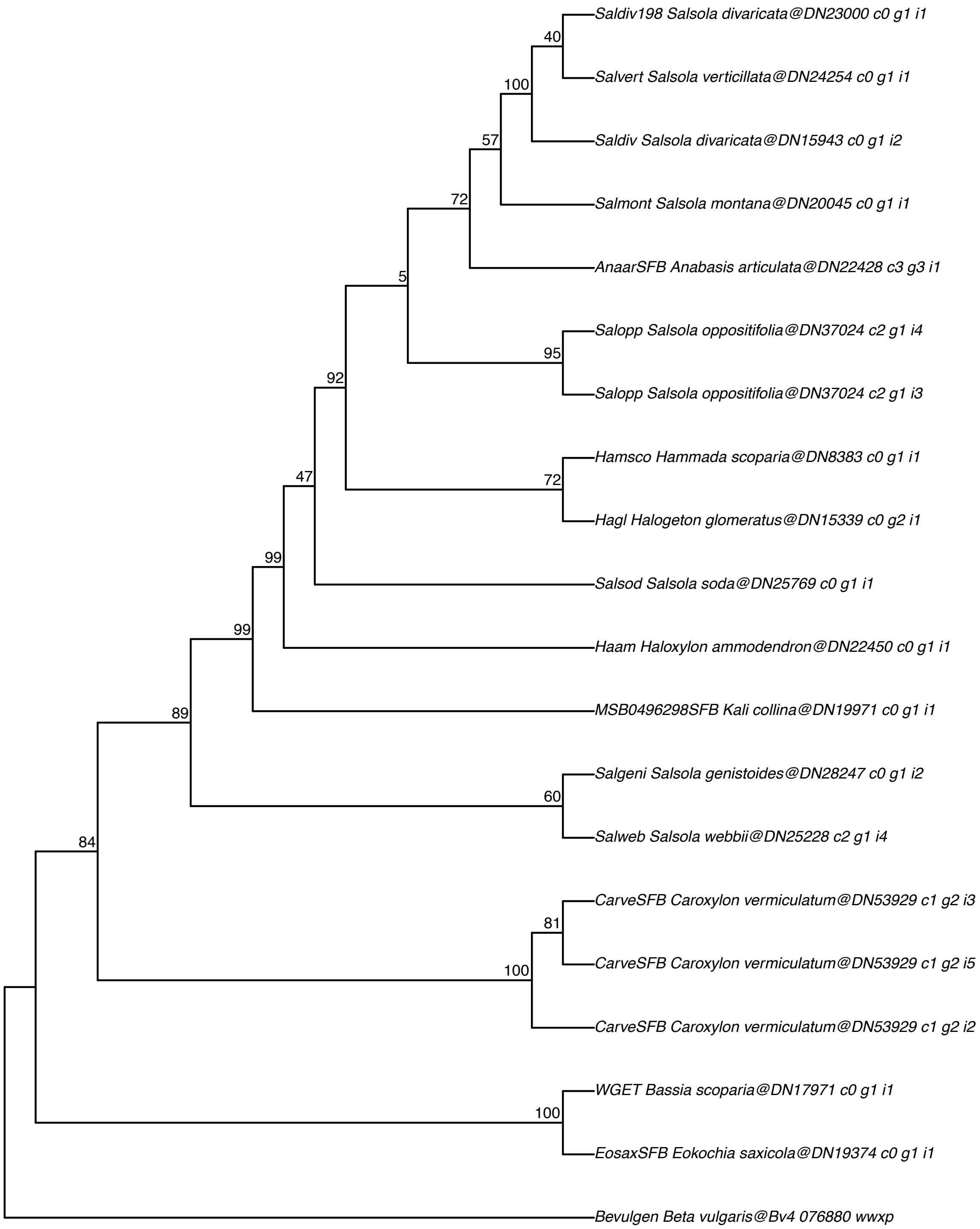

#### 2–phosphoglycolate phosphatase (PGP1/PGLP1)–AT5G36700

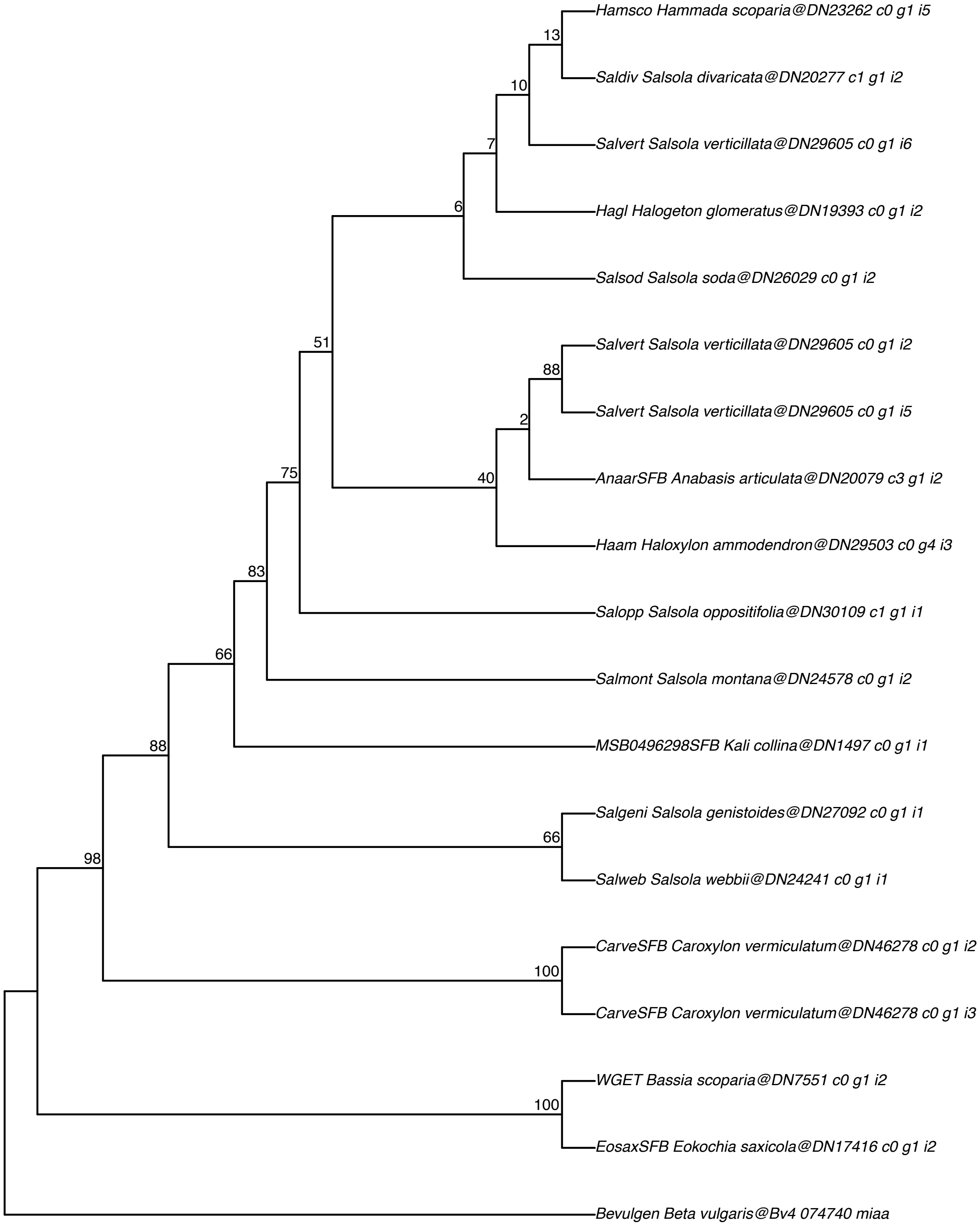

#### RuBisCO small subunit 1A−AT1G67090

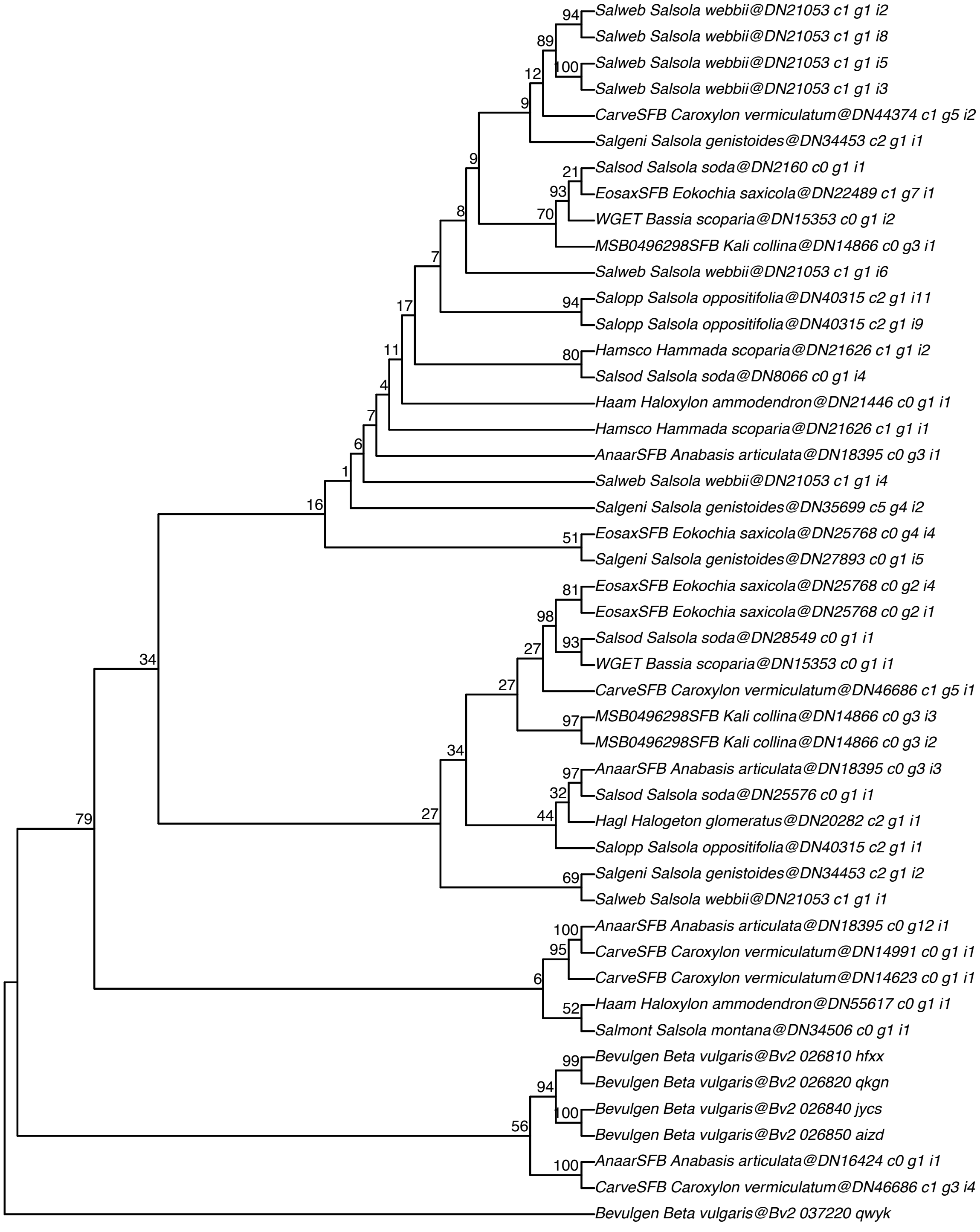

#### RuBisCO small subunit 3b–AT5G38410

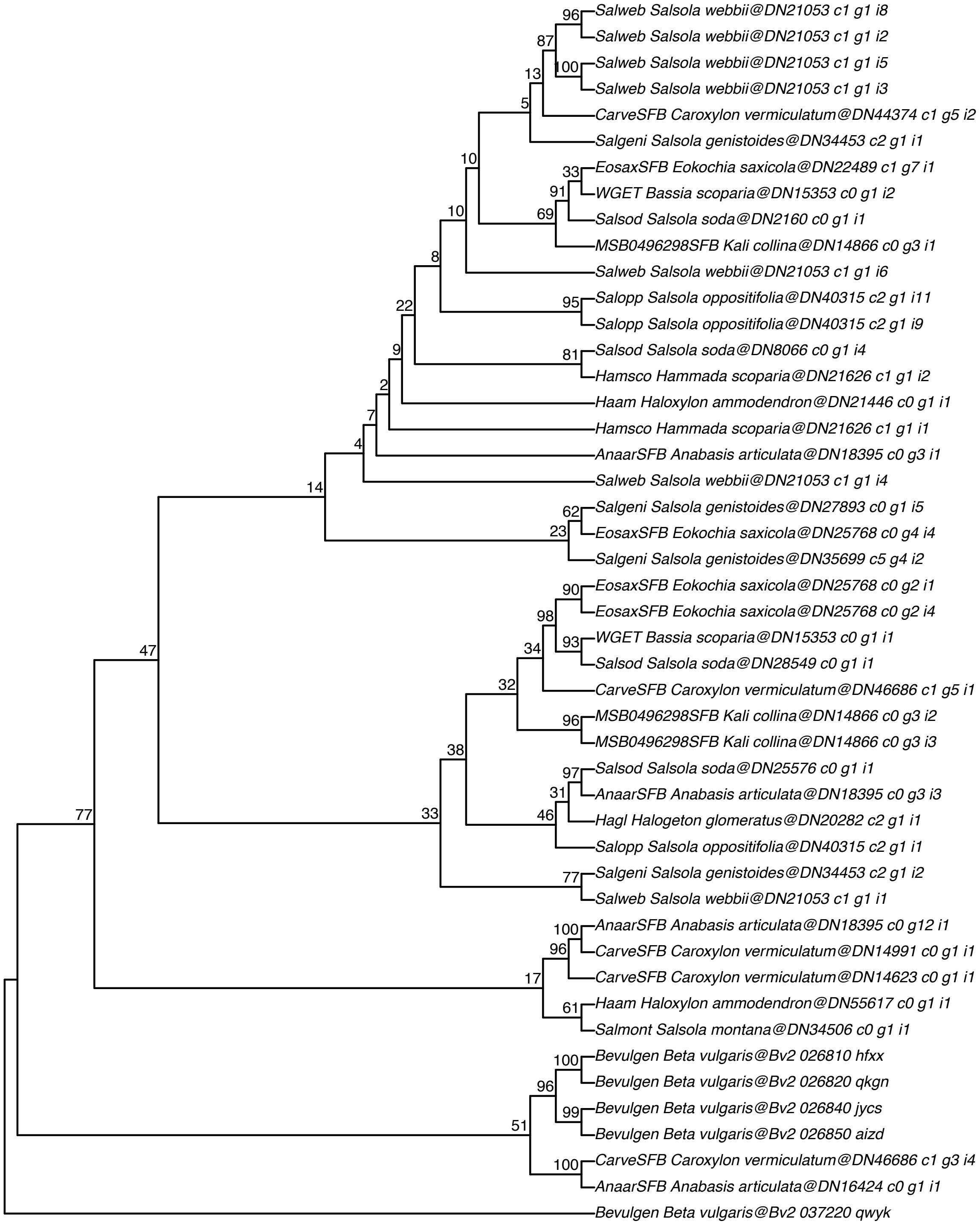

**Table S1**: Populations (named by living collection number in the Botanical Garden Mainz; roman numbers indicate different mother plants, see Fig. 1) included in the carbon isotope measurements, $CO_2$ compensation point measurements and PEPC activity measurements, including sampling location and collector. Vouchers are available at the Herbarium MJG in Mainz (https://mjg.jacq.org) and the Botanical Garden Munich (Living collection (LS)).

| LS No. | *Salsola* species | Location | Lat. N | Lon. E | Collector |
|---|---|---|---|---|---|
| 169 | *S. verticillata* Schousb. | SW Morocco, Isle de Mogador | 31.496319 | -9.785481 | H. Freitag |
| 171 | *S. verticillata* Schousb. | SW Morocco, S Essaouira | 31.466781 | -9.759214 | H. Freitag |
| 172 | *S. deschaseauxiana* Litard. & Maire | SW Morocco, SW Tamri | 30.659489 | -9.884175 | H. Freitag |
| 175 | *S. gymnomaschala* Maire | SW Morocco, SSW El Ouatia | 28.172481 | -11.869994 | H. Freitag |
| 176 | *S. gymnomaschala* Maire | SW Morocco, Merlift | 29.633703 | -10.006381 | H. Freitag |
| 177 | *S. gymnomaschala* Maire | SW Morocco, SW Guelmim | 28.900467 | -10.179222 | H. Freitag |
| 179 | *S. gymnomaschala* Maire | SW Morocco, SW La Ouatia | 28.446833 | -11.377308 | H. Freitag |
| 183 | *S. divaricata* Moq. | Lanzarote, Órzola | 29.206912 | -13.429706 | J. Gil González |
| 184 | *S. divaricata* Moq. | Lanzarote, Punta Mujeres | 29.146182 | -13.451436 | J. Gil González |
| 185 | *S. divaricata* Moq. | La Graciosa, Montana Amarilla | 29.22114 | -13.536032 | J. Gil González |
| 186 | *S. deschaseauxiana* Litard. & Maire | SW Marokko, N Agadir, Taghazout | 30.546858 | -9.716233 | H. Freitag |
| 187 | *S. divaricata* Moq. | La Graciosa, Playa Francesa | 29.220718 | -13.532952 | J. Gil González |
| 188 | *S. divaricata* Moq. | La Graciosa, Playa Francesa | 29.221187 | -13.533717 | J. Gil González |
| 191 | *S. divaricata* Moq. | Fuerteventura, Matas Blancas | 28.169453 | -14.204983 | S. Scholz |
| 192 | *S. divaricata* Moq. | Fuerteventura, Jandia, Costa Calma | 28.150678 | -14.201202 | S. Scholz |
| 193 | *S. divaricata* Moq. | Fuerteventura, Jandia, La Pared | 28.211006 | -14.225353 | S. Scholz |
| 194 | *S. divaricata* Moq. | Fuerteventura, Jandia, El Rayon | 28.131483 | -14.267744 | S. Scholz |
| 201 | *S. verticillata* Schousb. | SW Morocco, Isle de Mogador | 31.496319 | -9.785481 | H. Freitag |
| 208 | *S. divaricata* Moq. | Lanzarote, Tras Tenesar | 29.079891 | -13.710841 | J. Gil González |
| 182 I-IV | *S. divaricata* Moq. | La Gomera | 28.099444 | -17.349444 | V. Boehlke |
| 195 I-IV | *S. divaricata* Moq. | Tenerife, Punta de Abona (Arico) | 28.155156 | -16.43525 | R. Barone & F. Hernández |
| 196 III & V | *S. divaricata* Moq. | Gran Canaria, La Palmita (Agaete) | 28.098711 | -15.707096 | M. Olangua Corral |
| 198 I-III | *S. divaricata* Moq. | Gran Canaria, Cuesta Ramón, Jinamar | 28.046296 | -15.42001 | M. Olangua Corral |
| 200 I-II | *S. divaricata* Moq. | Tenerife, Charco del Viento (La Guancha) | 28.398921 | -16.671921 | R. Barone & F. Hernández |
| 248 I-IV | *S. divaricata* Moq. | Tenerife, Callao Chico - Punta Blanca (Guia de Isora) | 28.220458 | -16.839101 | R. Barone & F. Hernández |
| 255 I-III | *S. divaricata* Moq. | La Graciosa | 29.221652 | -13.521839 | J. Gil González |
| 67 | *S. webbii* Moq. | Spain | | | E. Voznesenskaya |
| 173 | *S. oppositifolia* Desf. | SW Morocco, 18 km N Agadir | | | H. Freitag |

**Table S2**: Gene and character occupancy of the concatenated matrix. The concatenated matrix comprises 991 genes, 17 taxa and 1427449 aligned columns with an overall matrix occupancy of 0.86.

| Taxon | No. orthologs | No. total characters | % orthologs | % characters |
|---|---|---|---|---|
| WGET_Bassia_scoparia | 991 | 924926 | 1 | 0.65 |
| Salsod_Salsola_soda | 991 | 1307319 | 1 | 0.92 |
| EosaxSFB_Eokochia_saxicola | 991 | 1320616 | 1 | 0.93 |
| Saldiv_Salsola_divaricata | 991 | 1273083 | 1 | 0.89 |
| Salopp_Salsola_oppositifolia | 991 | 1224330 | 1 | 0.86 |
| Bevulgen_Beta_vulgaris | 991 | 1382065 | 1 | 0.97 |
| Salweb_Salsola_webbii | 991 | 1270418 | 1 | 0.89 |
| Salgeni_Salsola_genistoides | 991 | 1270815 | 1 | 0.89 |
| Salvert_Salsola_verticillata | 991 | 1246077 | 1 | 0.87 |
| AnaarSFB_Anabasis_articulata | 991 | 1298566 | 1 | 0.91 |
| Hagl_Halogeton_glomeratus | 991 | 844209 | 1 | 0.59 |
| CarveSFB_Caroxylon_vermiculatum | 991 | 1181147 | 1 | 0.83 |
| Haam_Haloxylon_ammodendron | 991 | 1282382 | 1 | 0.90 |
| MSB0496298SFB_Kali_collina | 991 | 1344590 | 1 | 0.94 |
| Hamsco_Hammada_scoparia | 991 | 1283882 | 1 | 0.90 |
| Salmont_Salsola_montana | 991 | 1259920 | 1 | 0.88 |
| Saldiv198_Salsola_divaricata | 991 | 1243857 | 1 | 0.87 |

**Table S3:** HyDe results from the 298 significant tests.

| P1 | Hybrid | P2 | Zscore | Pvalue | Gamma |
|---|---|---|---|---|---|
| WGET | Salgeni | Salsod | 4.585159791 | 2.27E-06 | 0.034236573 |
| WGET | EosaxSFB | Saldiv | 6.565820377 | 2.60E-11 | 0.979288801 |
| WGET | EosaxSFB | Haam | 5.049458393 | 2.22E-07 | 0.983938636 |
| WGET | EosaxSFB | Salweb | 8.792344413 | 0 | 0.97311096 |
| WGET | EosaxSFB | Salgeni | 8.111523406 | 2.22E-16 | 0.975175644 |
| WGET | EosaxSFB | Salvert | 6.20812026 | 2.69E-10 | 0.980082988 |
| WGET | EosaxSFB | AnaarSFB | 4.192472493 | 1.38E-05 | 0.986655511 |
| WGET | EosaxSFB | Salmont | 7.108810816 | 5.89E-13 | 0.977856624 |
| WGET | EosaxSFB | Saldiv198 | 6.060892165 | 6.79E-10 | 0.980649408 |
| WGET | Saldiv | Salopp | 4.944712458 | 3.82E-07 | 0.017700895 |
| WGET | Salweb | Salopp | 6.35779125 | 1.03E-10 | 0.04736038 |
| WGET | Salgeni | Salopp | 6.727880895 | 8.66E-12 | 0.04781432 |
| WGET | Saldiv198 | Salopp | 4.045980949 | 2.61E-05 | 0.014909091 |
| WGET | Salweb | Haam | 5.141131886 | 1.37E-07 | 0.037221859 |
| WGET | Salweb | AnaarSFB | 6.198357579 | 2.86E-10 | 0.046806395 |
| WGET | Salweb | Hamsco | 5.295481905 | 5.95E-08 | 0.040556492 |
| WGET | Salgeni | AnaarSFB | 6.026082673 | 8.43E-10 | 0.0440196 |
| WGET | Salgeni | Hamsco | 4.276734842 | 9.49E-06 | 0.032103656 |
| Salsod | Saldiv | EosaxSFB | 4.782687601 | 8.66E-07 | 0.984466652 |
| Salsod | Salweb | EosaxSFB | 10.05566386 | 0 | 0.936257102 |
| Salsod | Salgeni | EosaxSFB | 10.03562959 | 0 | 0.940186287 |
| Salsod | Salvert | EosaxSFB | 4.097713258 | 2.09E-05 | 0.986331308 |
| Salsod | Salmont | EosaxSFB | 4.683695492 | 1.41E-06 | 0.984572333 |
| Salsod | Saldiv198 | EosaxSFB | 4.391824089 | 5.62E-06 | 0.98553292 |
| Salsod | Haam | Saldiv | 5.707758148 | 5.74E-09 | 0.214068606 |
| Salsod | Saldiv | Salweb | 20.56101476 | 0 | 0.86579662 |
| Salsod | Saldiv | Salgeni | 20.44585378 | 0 | 0.852539143 |
| Salsod | Saldiv | Salvert | 4.272083258 | 9.69E-06 | 0.017622918 |
| Salsod | Saldiv | CarveSFB | 5.626941175 | 9.20E-09 | 0.972958057 |
| Salsod | Saldiv | MSB0496298SFB | 11.30241179 | 0 | 0.895756733 |
| Salsod | Saldiv | Salmont | 5.770125522 | 3.97E-09 | 0.140606061 |
| Salsod | Haam | Salopp | 5.364206793 | 4.07E-08 | 0.103004292 |
| Salsod | Salopp | Hagl | 7.108360031 | 5.91E-13 | 0.18989547 |
| Salsod | Salopp | Salmont | 5.712762461 | 5.57E-09 | 0.412379421 |
| Salsod | Haam | Salweb | 6.818938197 | 4.61E-12 | 0.951466287 |
| Salsod | Haam | Salgeni | 7.762733223 | 4.22E-15 | 0.940283228 |
| Salsod | Haam | Salvert | 8.116675529 | 2.22E-16 | 0.255539143 |
| Salsod | Haam | Hagl | 11.46221923 | 0 | 0.254019293 |
| Salsod | Haam | Hamsco | 4.412031797 | 5.12E-06 | 0.084210526 |
| Salsod | Haam | Salmont | 9.975122624 | 0 | 0.482443482 |
| Salsod | Haam | Saldiv198 | 7.106554197 | 5.99E-13 | 0.235522736 |
| Salsod | Salgeni | Salweb | 10.97895401 | 0 | 0.182199526 |
| Salsod | Salvert | Salweb | 19.89714122 | 0 | 0.866604186 |
| Salweb | Salsod | Hagl | 7.898006944 | 1.44E-15 | 0.075282906 |
| Salsod | Salweb | CarveSFB | 12.94929644 | 0 | 0.84991511 |
| Salsod | Salmont | Salweb | 26.15863312 | 0 | 0.828288973 |
| Salsod | Saldiv198 | Salweb | 20.11528897 | 0 | 0.864944995 |
| Salsod | Salvert | Salgeni | 19.32321627 | 0 | 0.856534696 |

| | | | | | |
|---|---|---|---|---|---|
| Salgeni | Salsod | Hagl | 8.49688637 | 0 | 0.087392366 |
| Salsod | Salgeni | CarveSFB | 13.22660459 | 0 | 0.864336373 |
| Salgeni | Salsod | MSB0496298SFB | 5.143889026 | 1.35E-07 | 0.201954397 |
| Salsod | Salmont | Salgeni | 24.24491145 | 0 | 0.821674794 |
| Salsod | Saldiv198 | Salgeni | 20.16278284 | 0 | 0.853918443 |
| Salsod | Salvert | CarveSFB | 4.679823578 | 1.44E-06 | 0.976897876 |
| Salsod | Salvert | MSB0496298SFB | 10.8863814 | 0 | 0.896024781 |
| AnaarSFB | Salsod | Hagl | 9.406917582 | 0 | 0.555496548 |
| Salsod | Hamsco | Hagl | 7.297679675 | 1.47E-13 | 0.210931768 |
| Hagl | Salsod | Salmont | 7.347251606 | 1.02E-13 | 0.461708395 |
| Salsod | Salmont | CarveSFB | 6.368236035 | 9.60E-11 | 0.968802888 |
| Salsod | Saldiv198 | CarveSFB | 4.464983008 | 4.01E-06 | 0.978094753 |
| Salsod | Salmont | MSB0496298SFB | 15.50800725 | 0 | 0.855128335 |
| Salsod | Saldiv198 | MSB0496298SFB | 10.71078863 | 0 | 0.898024615 |
| Salsod | Saldiv198 | Salmont | 4.702519236 | 1.29E-06 | 0.120678617 |
| EosaxSFB | Saldiv | Salopp | 7.536049831 | 2.44E-14 | 0.022494367 |
| EosaxSFB | Salweb | Saldiv | 6.898439663 | 2.64E-12 | 0.037937909 |
| EosaxSFB | Salgeni | Saldiv | 5.627704916 | 9.16E-09 | 0.029816005 |
| EosaxSFB | Saldiv | AnaarSFB | 4.390333533 | 5.66E-06 | 0.015019763 |
| EosaxSFB | Saldiv | MSB0496298SFB | 4.121285696 | 1.88E-05 | 0.02176911 |
| EosaxSFB | Saldiv | Hamsco | 5.290127216 | 6.12E-08 | 0.016225388 |
| EosaxSFB | Salweb | Salopp | 11.79304942 | 0 | 0.072469024 |
| EosaxSFB | Salgeni | Salopp | 11.25871137 | 0 | 0.066404184 |
| EosaxSFB | Salvert | Salopp | 5.712194045 | 5.59E-09 | 0.017331601 |
| EosaxSFB | Salmont | Salopp | 6.899261891 | 2.63E-12 | 0.021527518 |
| EosaxSFB | Saldiv198 | Salopp | 6.69499717 | 1.08E-11 | 0.020307784 |
| EosaxSFB | Salweb | Haam | 10.6322501 | 0 | 0.062899603 |
| EosaxSFB | Salgeni | Haam | 8.596411588 | 0 | 0.048691216 |
| EosaxSFB | Salmont | Haam | 4.500454297 | 3.39E-06 | 0.013527575 |
| EosaxSFB | Salweb | Salvert | 6.487875303 | 4.37E-11 | 0.036247459 |
| EosaxSFB | Salweb | AnaarSFB | 12.01563908 | 0 | 0.073686644 |
| EosaxSFB | Salweb | Hagl | 7.751989675 | 4.55E-15 | 0.069515057 |
| EosaxSFB | Salweb | CarveSFB | 8.898134415 | 0 | 0.111950244 |
| EosaxSFB | Salweb | MSB0496298SFB | 11.47774515 | 0 | 0.070553902 |
| EosaxSFB | Salweb | Hamsco | 12.0874961 | 0 | 0.073932441 |
| EosaxSFB | Salweb | Salmont | 7.096533953 | 6.44E-13 | 0.036989867 |
| EosaxSFB | Salweb | Saldiv198 | 7.474130383 | 3.92E-14 | 0.041018551 |
| EosaxSFB | Salgeni | Salvert | 6.501756647 | 3.99E-11 | 0.034423408 |
| EosaxSFB | Salgeni | AnaarSFB | 10.17406337 | 0 | 0.060972851 |
| EosaxSFB | Salgeni | Hagl | 6.791226369 | 5.59E-12 | 0.059137734 |
| EosaxSFB | Salgeni | CarveSFB | 7.929197183 | 1.11E-15 | 0.104659169 |
| EosaxSFB | Salgeni | MSB0496298SFB | 10.0414987 | 0 | 0.061541084 |
| EosaxSFB | Salgeni | Hamsco | 9.580667192 | 0 | 0.057650678 |
| EosaxSFB | Salgeni | Salmont | 5.874878422 | 2.12E-09 | 0.029173076 |
| EosaxSFB | Salgeni | Saldiv198 | 5.955404846 | 1.30E-09 | 0.031607905 |
| EosaxSFB | Salvert | AnaarSFB | 5.005085623 | 2.80E-07 | 0.017174557 |
| EosaxSFB | Salvert | MSB0496298SFB | 4.142203746 | 1.72E-05 | 0.021954498 |
| EosaxSFB | Salvert | Hamsco | 5.837333155 | 2.66E-09 | 0.017890368 |
| EosaxSFB | Salmont | AnaarSFB | 5.428600063 | 2.85E-08 | 0.018322155 |
| EosaxSFB | Saldiv198 | AnaarSFB | 5.504049635 | 1.86E-08 | 0.018765018 |
| EosaxSFB | Salmont | MSB0496298SFB | 4.563374463 | 2.52E-06 | 0.023065834 |

| | | | | | |
|---|---|---|---|---|---|
| EosaxSFB | Saldiv198 | MSB0496298SFB | 4.153172878 | 1.64E-05 | 0.021983512 |
| EosaxSFB | Salmont | Hamsco | 5.814043657 | 3.06E-09 | 0.018562158 |
| EosaxSFB | Saldiv198 | Hamsco | 5.013416035 | 2.68E-07 | 0.015374691 |
| Saldiv | Haam | Salopp | 6.87142818 | 3.20E-12 | 0.165722802 |
| Salopp | Saldiv | Salweb | 17.90478377 | 0 | 0.890045942 |
| Salopp | Saldiv | Salgeni | 17.15921203 | 0 | 0.884757119 |
| Saldiv | Salopp | Hagl | 13.93916517 | 0 | 0.326434619 |
| Salopp | Saldiv | CarveSFB | 8.267401689 | 0 | 0.96463917 |
| Salopp | Saldiv | MSB0496298SFB | 10.80766285 | 0 | 0.909840039 |
| Salopp | Saldiv | Salmont | 11.45519013 | 0 | 0.305015353 |
| Haam | Saldiv | Salweb | 13.87468221 | 0 | 0.917559524 |
| Haam | Saldiv | Salgeni | 12.43262669 | 0 | 0.918627451 |
| Saldiv | Haam | Hagl | 18.58608835 | 0 | 0.380989995 |
| Haam | Saldiv | CarveSFB | 4.608599585 | 2.03E-06 | 0.980727273 |
| Haam | Saldiv | MSB0496298SFB | 7.033665046 | 1.01E-12 | 0.943846297 |
| Saldiv | Haam | Hamsco | 10.06609728 | 0 | 0.195944353 |
| Haam | Saldiv | Salmont | 13.6510704 | 0 | 0.403737659 |
| Saldiv | Salgeni | Salweb | 9.043801223 | 0 | 0.235469449 |
| Salweb | Saldiv | AnaarSFB | 17.04918226 | 0 | 0.12733764 |
| Salweb | Saldiv | Hagl | 24.11900274 | 0 | 0.189183546 |
| Saldiv | Salweb | CarveSFB | 10.07160839 | 0 | 0.900063586 |
| Salweb | Saldiv | MSB0496298SFB | 17.40857657 | 0 | 0.327142332 |
| Salweb | Saldiv | Hamsco | 19.54570103 | 0 | 0.121212121 |
| Saldiv | Salmont | Salweb | 6.968022026 | 1.62E-12 | 0.958575522 |
| Salgeni | Saldiv | AnaarSFB | 18.6427319 | 0 | 0.148107912 |
| Salgeni | Saldiv | Hagl | 24.82946313 | 0 | 0.205622933 |
| Saldiv | Salgeni | CarveSFB | 8.041948915 | 4.44E-16 | 0.925614877 |
| Salgeni | Saldiv | MSB0496298SFB | 21.34109942 | 0 | 0.417562724 |
| Salgeni | Saldiv | Hamsco | 19.84415417 | 0 | 0.131753555 |
| Saldiv | Salmont | Salgeni | 6.089539834 | 5.68E-10 | 0.96026018 |
| Saldiv | Saldiv198 | Salvert | 6.156917569 | 3.72E-10 | 0.591973244 |
| AnaarSFB | Saldiv | Hagl | 9.249009006 | 0 | 0.464408726 |
| AnaarSFB | Saldiv | CarveSFB | 4.943877301 | 3.83E-07 | 0.974623407 |
| AnaarSFB | Saldiv | MSB0496298SFB | 10.44808687 | 0 | 0.89160221 |
| Hagl | Saldiv | CarveSFB | 5.18910726 | 1.06E-07 | 0.966176596 |
| Hagl | Saldiv | MSB0496298SFB | 11.99263694 | 0 | 0.858792185 |
| Saldiv | Hamsco | Hagl | 11.33995287 | 0 | 0.31033237 |
| Hagl | Saldiv | Salmont | 11.79341429 | 0 | 0.233402184 |
| CarveSFB | Saldiv | MSB0496298SFB | 4.824929382 | 7.01E-07 | 0.043704024 |
| CarveSFB | Saldiv | Hamsco | 6.468190388 | 4.98E-11 | 0.028866959 |
| MSB0496298SFB | Saldiv | Hamsco | 10.69814119 | 0 | 0.090042076 |
| Saldiv | Salmont | MSB0496298SFB | 4.912048899 | 4.51E-07 | 0.963288719 |
| Hamsco | Saldiv | Salmont | 12.75633826 | 0 | 0.291512915 |
| Salopp | Haam | Salweb | 4.57783631 | 2.35E-06 | 0.976030362 |
| Salopp | Haam | Salgeni | 4.935712457 | 4.00E-07 | 0.972190285 |
| Salopp | Haam | Salvert | 7.093568603 | 6.58E-13 | 0.832402235 |
| Salopp | Haam | Saldiv198 | 8.596415122 | 0 | 0.808176981 |
| Salopp | Salgeni | Salweb | 9.148263792 | 0 | 0.174479167 |
| Salopp | Salvert | Salweb | 16.8990144 | 0 | 0.895301805 |
| Salweb | Salopp | Hagl | 12.12611343 | 0 | 0.081347729 |
| Salopp | Salweb | CarveSFB | 14.95810323 | 0 | 0.839246364 |

| | | | | | |
|---|---|---|---|---|---|
| Salopp | Salmont | Salweb | 22.99957879 | 0 | 0.850891152 |
| Salopp | Saldiv198 | Salweb | 16.95817685 | 0 | 0.893084261 |
| Salopp | Salvert | Salgeni | 14.84027202 | 0 | 0.89827601 |
| Salgeni | Salopp | Hagl | 13.14549818 | 0 | 0.09345887 |
| Salopp | Salgeni | CarveSFB | 13.86336078 | 0 | 0.862497411 |
| Salgeni | Salopp | MSB0496298SFB | 6.923898096 | 2.21E-12 | 0.255864309 |
| Salopp | Salmont | Salgeni | 21.57895061 | 0 | 0.845464043 |
| Salopp | Saldiv198 | Salgeni | 16.70916934 | 0 | 0.886964682 |
| Salvert | Salopp | Hagl | 15.10103255 | 0 | 0.33963964 |
| Salopp | Salvert | CarveSFB | 7.25859626 | 1.97E-13 | 0.968875176 |
| Salopp | Salvert | MSB0496298SFB | 10.12939476 | 0 | 0.913945409 |
| Salopp | Salvert | Salmont | 10.9776734 | 0 | 0.301162381 |
| AnaarSFB | Salopp | Hagl | 11.65534172 | 0 | 0.219506173 |
| Hagl | Salopp | MSB0496298SFB | 4.55782834 | 2.59E-06 | 0.956347276 |
| Hagl | Salopp | Hamsco | 4.041105018 | 2.66E-05 | 0.20746888 |
| Hagl | Salopp | Salmont | 17.11005601 | 0 | 0.697142857 |
| Hagl | Salopp | Saldiv198 | 14.5422651 | 0 | 0.670336391 |
| Salopp | Salmont | CarveSFB | 9.055931893 | 0 | 0.959459459 |
| Salopp | Saldiv198 | CarveSFB | 7.208917866 | 2.84E-13 | 0.968727554 |
| Salopp | Salmont | MSB0496298SFB | 15.08190782 | 0 | 0.866964105 |
| Salopp | Saldiv198 | MSB0496298SFB | 10.01167433 | 0 | 0.91439493 |
| Hamsco | Salopp | Salmont | 5.276469943 | 6.60E-08 | 0.903800217 |
| Salopp | Saldiv198 | Salmont | 10.53854683 | 0 | 0.295795247 |
| Haam | Salgeni | Salweb | 10.76055792 | 0 | 0.204767986 |
| Haam | Salvert | Salweb | 13.11509442 | 0 | 0.92185032 |
| Salweb | Haam | AnaarSFB | 4.88460171 | 5.19E-07 | 0.040616874 |
| Salweb | Haam | Hagl | 15.75752156 | 0 | 0.098836184 |
| Haam | Salweb | CarveSFB | 13.67073548 | 0 | 0.856976127 |
| Salweb | Haam | MSB0496298SFB | 7.564411233 | 1.97E-14 | 0.199748428 |
| Salweb | Haam | Hamsco | 6.246749727 | 2.10E-10 | 0.031717235 |
| Haam | Salmont | Salweb | 18.83589368 | 0 | 0.879059795 |
| Haam | Saldiv198 | Salweb | 13.08966001 | 0 | 0.92114094 |
| Haam | Salvert | Salgeni | 10.68771378 | 0 | 0.929485592 |
| Salgeni | Haam | AnaarSFB | 6.292968424 | 1.56E-10 | 0.054504377 |
| Salgeni | Haam | Hagl | 16.86270022 | 0 | 0.109855791 |
| Haam | Salgeni | CarveSFB | 11.39938038 | 0 | 0.888155922 |
| Salgeni | Haam | MSB0496298SFB | 12.28764639 | 0 | 0.325006308 |
| Salgeni | Haam | Hamsco | 7.752640855 | 4.55E-15 | 0.040848185 |
| Haam | Salmont | Salgeni | 16.57717645 | 0 | 0.881824525 |
| Haam | Saldiv198 | Salgeni | 11.73542989 | 0 | 0.923685512 |
| Salvert | Haam | Hagl | 20.25290036 | 0 | 0.410068303 |
| Haam | Salvert | CarveSFB | 4.868988178 | 5.62E-07 | 0.980155039 |
| Haam | Salvert | MSB0496298SFB | 7.129717099 | 5.06E-13 | 0.943352278 |
| Salvert | Haam | Hamsco | 10.45429724 | 0 | 0.209296666 |
| Haam | Salvert | Salmont | 14.26370689 | 0 | 0.407067618 |
| AnaarSFB | Haam | Hagl | 14.56116529 | 0 | 0.245749614 |
| AnaarSFB | Haam | Salmont | 5.02049078 | 2.58E-07 | 0.246745897 |
| Hagl | Haam | MSB0496298SFB | 7.871279335 | 1.78E-15 | 0.932286446 |
| Hagl | Haam | Hamsco | 6.32683206 | 1.26E-10 | 0.264129181 |
| Hagl | Haam | Salmont | 19.81975265 | 0 | 0.679387667 |
| Hagl | Haam | Saldiv198 | 19.8955739 | 0 | 0.603487491 |

| | | | | | |
|---|---|---|---|---|---|
| Haam | Salmont | CarveSFB | 5.939742182 | 1.43E-09 | 0.973925718 |
| Haam | Saldiv198 | CarveSFB | 4.738225497 | 1.08E-06 | 0.980619548 |
| Haam | Salmont | MSB0496298SFB | 11.9448539 | 0 | 0.894879791 |
| Haam | Saldiv198 | MSB0496298SFB | 5.722259084 | 5.27E-09 | 0.953747854 |
| Hamsco | Haam | Salmont | 8.922549934 | 0 | 0.856812488 |
| Hamsco | Haam | Saldiv198 | 10.65323024 | 0 | 0.792102759 |
| Haam | Saldiv198 | Salmont | 13.51428184 | 0 | 0.427665921 |
| Salweb | Salgeni | Salvert | 8.688937522 | 0 | 0.768948655 |
| Salweb | Salgeni | AnaarSFB | 7.844011485 | 2.22E-15 | 0.852556237 |
| Salweb | Salgeni | Hagl | 6.334185378 | 1.20E-10 | 0.882250868 |
| Salweb | Salgeni | MSB0496298SFB | 4.503255109 | 3.35E-06 | 0.91593973 |
| Salweb | Salgeni | Hamsco | 8.028117854 | 4.44E-16 | 0.851266464 |
| Salweb | Salgeni | Salmont | 9.480734138 | 0 | 0.725392059 |
| Salweb | Salgeni | Saldiv198 | 8.087084788 | 3.33E-16 | 0.776500639 |
| Salweb | Salvert | AnaarSFB | 16.19182742 | 0 | 0.122084453 |
| Salweb | Salvert | Hagl | 24.55707078 | 0 | 0.193283504 |
| Salvert | Salweb | CarveSFB | 10.0547137 | 0 | 0.900191939 |
| Salweb | Salvert | MSB0496298SFB | 17.72693665 | 0 | 0.324353643 |
| Salweb | Salvert | Hamsco | 19.50437584 | 0 | 0.120850429 |
| Salweb | Salmont | Salvert | 6.954149069 | 1.78E-12 | 0.041431262 |
| Salweb | AnaarSFB | Hagl | 9.011024562 | 0 | 0.096992182 |
| AnaarSFB | Salweb | CarveSFB | 15.12018933 | 0 | 0.837812002 |
| Salweb | Salmont | AnaarSFB | 22.60949005 | 0 | 0.162300636 |
| Salweb | Saldiv198 | AnaarSFB | 17.05049031 | 0 | 0.128855678 |
| Hagl | Salweb | CarveSFB | 10.9155779 | 0 | 0.822820037 |
| Salweb | MSB0496298SFB | Hagl | 6.683779308 | 1.17E-11 | 0.221727846 |
| Salweb | Hamsco | Hagl | 10.2088261 | 0 | 0.072561497 |
| Salweb | Salmont | Hagl | 27.32319904 | 0 | 0.234395898 |
| Salweb | Saldiv198 | Hagl | 23.62575094 | 0 | 0.190157565 |
| CarveSFB | Salweb | MSB0496298SFB | 14.69213436 | 0 | 0.163386039 |
| CarveSFB | Salweb | Hamsco | 15.79984094 | 0 | 0.167875539 |
| CarveSFB | Salweb | Salmont | 8.842287694 | 0 | 0.083010006 |
| CarveSFB | Salweb | Saldiv198 | 10.37746033 | 0 | 0.102421256 |
| Salweb | Salmont | MSB0496298SFB | 22.98996008 | 0 | 0.367628993 |
| Salweb | Saldiv198 | MSB0496298SFB | 17.66957367 | 0 | 0.330800593 |
| Salweb | Salmont | Hamsco | 25.21531511 | 0 | 0.164387747 |
| Salweb | Saldiv198 | Hamsco | 18.61516912 | 0 | 0.1164807 |
| Salweb | Salmont | Saldiv198 | 6.897479439 | 2.66E-12 | 0.041764312 |
| Salgeni | Salvert | AnaarSFB | 17.21157239 | 0 | 0.138784144 |
| Salgeni | Salvert | Hagl | 24.15942526 | 0 | 0.202491862 |
| Salvert | Salgeni | CarveSFB | 8.675787262 | 0 | 0.920433098 |
| Salgeni | Salvert | MSB0496298SFB | 21.21131436 | 0 | 0.409917945 |
| Salgeni | Salvert | Hamsco | 18.82350668 | 0 | 0.12505978 |
| Salgeni | Salmont | Salvert | 6.643571298 | 1.54E-11 | 0.042868414 |
| Salgeni | AnaarSFB | Hagl | 9.553590326 | 0 | 0.108965323 |
| AnaarSFB | Salgeni | CarveSFB | 12.52105579 | 0 | 0.869833436 |
| Salgeni | AnaarSFB | MSB0496298SFB | 5.270955002 | 6.80E-08 | 0.209594096 |
| Salgeni | Salmont | AnaarSFB | 23.67848354 | 0 | 0.182057353 |
| Salgeni | Saldiv198 | AnaarSFB | 18.38922322 | 0 | 0.146182751 |
| Hagl | Salgeni | CarveSFB | 9.51836499 | 0 | 0.852696729 |
| Salgeni | Hamsco | Hagl | 10.62808723 | 0 | 0.078412009 |

| | | | | | |
|---|---|---|---|---|---|
| Salgeni | Salmont | Hagl | 26.98319836 | 0 | 0.246933962 |
| Salgeni | Saldiv198 | Hagl | 24.03033938 | 0 | 0.202331108 |
| CarveSFB | Salgeni | MSB0496298SFB | 13.55721065 | 0 | 0.145554518 |
| CarveSFB | Salgeni | Hamsco | 12.44023581 | 0 | 0.128662159 |
| CarveSFB | Salgeni | Salmont | 7.80205965 | 3.11E-15 | 0.067636986 |
| CarveSFB | Salgeni | Saldiv198 | 8.744908854 | 0 | 0.080523402 |
| Salgeni | Hamsco | MSB0496298SFB | 5.74931661 | 4.49E-09 | 0.215363031 |
| Salgeni | Salmont | MSB0496298SFB | 24.57844228 | 0 | 0.435262577 |
| Salgeni | Saldiv198 | MSB0496298SFB | 21.1395335 | 0 | 0.410562489 |
| Salgeni | Salmont | Hamsco | 24.28671578 | 0 | 0.171238502 |
| Salgeni | Saldiv198 | Hamsco | 19.29030977 | 0 | 0.126686776 |
| Salgeni | Salmont | Saldiv198 | 5.74639638 | 4.57E-09 | 0.037955144 |
| AnaarSFB | Salvert | Hagl | 10.29613445 | 0 | 0.507079182 |
| AnaarSFB | Salvert | CarveSFB | 5.361371465 | 4.14E-08 | 0.972798605 |
| AnaarSFB | Salvert | MSB0496298SFB | 10.8980864 | 0 | 0.887505558 |
| Hagl | Salvert | CarveSFB | 5.987757491 | 1.07E-09 | 0.96165838 |
| Hagl | Salvert | MSB0496298SFB | 12.13681285 | 0 | 0.856071964 |
| Salvert | Hamsco | Hagl | 12.40047822 | 0 | 0.334989354 |
| Hagl | Salvert | Salmont | 10.57669569 | 0 | 0.208603418 |
| CarveSFB | Salvert | MSB0496298SFB | 4.65601228 | 1.61E-06 | 0.04206501 |
| CarveSFB | Salvert | Hamsco | 7.074282549 | 7.56E-13 | 0.030868099 |
| MSB0496298SFB | Salvert | Hamsco | 10.61347891 | 0 | 0.090413318 |
| Salvert | Salmont | MSB0496298SFB | 4.452160674 | 4.25E-06 | 0.966593069 |
| Hamsco | Salvert | Salmont | 12.95535791 | 0 | 0.292182227 |
| AnaarSFB | Hamsco | Hagl | 12.18438878 | 0 | 0.241292277 |
| AnaarSFB | Saldiv198 | Hagl | 8.962728833 | 0 | 0.486388385 |
| AnaarSFB | Salmont | CarveSFB | 7.049044711 | 9.07E-13 | 0.96499743 |
| AnaarSFB | Saldiv198 | CarveSFB | 5.719799495 | 5.35E-09 | 0.970961674 |
| AnaarSFB | Salmont | MSB0496298SFB | 14.92649746 | 0 | 0.85303241 |
| AnaarSFB | Saldiv198 | MSB0496298SFB | 10.18203219 | 0 | 0.892743403 |
| Hagl | Salmont | CarveSFB | 5.179017695 | 1.12E-07 | 0.963833797 |
| Hagl | Saldiv198 | CarveSFB | 4.912185345 | 4.51E-07 | 0.967652415 |
| Hagl | Hamsco | MSB0496298SFB | 4.361686993 | 6.46E-06 | 0.957275244 |
| Hagl | Salmont | MSB0496298SFB | 14.74357902 | 0 | 0.807473599 |
| Hagl | Saldiv198 | MSB0496298SFB | 11.24252978 | 0 | 0.861876742 |
| Hagl | Hamsco | Salmont | 13.77288158 | 0 | 0.720525188 |
| Hagl | Hamsco | Saldiv198 | 13.19918108 | 0 | 0.664776518 |
| Hagl | Saldiv198 | Salmont | 10.10278305 | 0 | 0.212054276 |
| CarveSFB | Salmont | MSB0496298SFB | 6.145705555 | 4.00E-10 | 0.051973684 |
| CarveSFB | Saldiv198 | MSB0496298SFB | 5.093423536 | 1.76E-07 | 0.045959905 |
| CarveSFB | Salmont | Hamsco | 8.617733095 | 0 | 0.039800726 |
| CarveSFB | Saldiv198 | Hamsco | 6.384513677 | 8.64E-11 | 0.02805878 |
| MSB0496298SFB | Salmont | Hamsco | 15.47131571 | 0 | 0.138619735 |
| MSB0496298SFB | Saldiv198 | Hamsco | 9.538613321 | 0 | 0.081750699 |
| MSB0496298SFB | Salmont | Saldiv198 | 5.049172875 | 2.22E-07 | 0.038639269 |
| Hamsco | Saldiv198 | Salmont | 13.54212132 | 0 | 0.314252674 |

**Table S4**: Fifty trees of photosynthesis pathway-associated genes characteristic.

| Pathway assignment | Gene annotation | Arabidopsis Tair_id | No. sequences | min sequence length | max sequence length | average sequence length | Sd agg sister to | bootstrap support (sister) | Sd agg in clade with | bootstrap support(clade) | notes on gene trees |
|---|---|---|---|---|---|---|---|---|---|---|---|
| C4 | Adenosine monophosphate kinase (AMK2) | AT5G47840 | 20 | 456 | 903 | 819 | C3 | 56 | C3 | 10 | no support on important nodes |
| C4 | Alanine aminotransferase (Ala-AT1) | AT1G17290 | 23 | 444 | 1584 | 1347 | both | 66 | neither (its own clade) | | Salmont in clade with C4 |
| C4 | Aspartat aminotransferase (Asp-AT5) | AT4G31990 | 23 | 324 | 1380 | 1185 | both | 70 | C4 | 7 | Salmont in clade with C4, Sd agg sister to this clade; one copy of Sv in C4 clade but no support |
| C4 | Beta carbonic anhydrase (BCA3) | AT1G23730 | 45 | 255 | 966 | 739 | Sv to C3; Sd sister to C4 | 88;81 | | | two gene copies, one nested clade is missing Salmont |
| C4 | Bile acid:sodium symporter family protein (BASS2) | AT2G26900 | 25 | 369 | 1266 | 921 | both | 60s | | | Salmont in clade with C4 |
| C4 | Bile acid:sodium symporter family protein (BASS4) | AT3G56160 | 48 | 309 | 1203 | 771 | | | C4;C3 | 99;94 | several gene copies, two clades |
| C4 | Dicarboxylate transporter (Dit1) | AT5G12860 | 21 | 525 | 1698 | 1387 | C3 | 86 | | | otherwise, many nodes with no support; several copies of Salvert |

| | | | | | | | | | | | |
|---|---|---|---|---|---|---|---|---|---|---|---|
| C4 | Dicarboxylate transporter (Dit2) | AT5G64280 | 43 | 342 | 1692 | 1355 | C3 | 57 | C3 | 85 | several copies, two clades, in one clade Salmot missing and no (or low) support, in other clade support higher than sister support |
| C4 | NADP-dependent malate dehydrogenase (NADP-MDH) | AT5G58330 | 18 | 684 | 1320 | 1244 | C4 | 61 | C4 | 96 | Salmont sister to Anabasis (BS=47), Sd agg in clade with C4 including Salsod (BS=96) |
| C4 | NADP-dependent malate enzyme (NADP-ME1) | AT2G19900 | 62 | 390 | 1722 | 1335 | C3 | 85 | both;C4 | 99; 62 | several gene copies, three clades |
| C4 | NADP-dependent malic enzyme (NADP-ME2) | AT5G11670 | 62 | 390 | 1743 | 1347 | C3;C3;C4 | 98;79;46 | both;C4 | 100;65 | several gene copies, three clades, in two C4 not monophyletic; in one Sd agg not monophyletic |
| C4 | NADP-dependent malic enzyme (NADP-ME4) | AT1G79750 | 65 | 300 | 1737 | 1315 | C3 | 66,95 | C4 | 61 | three gene copies in separate clades, very short branches |
| C4 | PEP carboxylase (PEPC1) | AT1G53310 | 38 | 549 | 2901 | 2341 | C4 | 98 | C3 | 100 | two gene copies/two clades |
| C4 | PEP carboxylase (PEPC2) | AT2G42600 | 38 | 549 | 2901 | 2341 | C4; C3 | 98;100 | | | two clades |

| | | | | | | | | | | | |
|---|---|---|---|---|---|---|---|---|---|---|---|
| C4 | PEP carboxylase kinase (PEPC-K1) | AT1G08650 | 24 | 456 | 840 | 793 | | | C3 | 45 | several copies of Sd agg in one clade with Salmont |
| C4 | PEP carboxylase kinase (PEPC-K2) | AT3G04530 | 24 | 456 | 840 | 793 | | | C3 | 53 | several copies of Sd agg |
| C4 | PEP carboxylase-related protein (PEPC-RP) | AT3G42628 | 35 | 360 | 2898 | 2134 | C4 | 99 | | | several gene copies, one clade missing Salmont and Sd agg. not monophyletic |
| C4 | PEP/phosphate translocator (PPT1/CUE1) | AT5G33320 | 20 | 483 | 1290 | 1225 | C4 | 78 | neither (its own clade) | 35 | several copies of Sd agg, low support on some important nodes |
| C4 | PEP/phosphate translocator (PPT2) | AT3G01550 | 31 | 477 | 1284 | 1172 | both | 100 | both | 94 | several copies of Sd and Sv; Sd agg is outgoup in one clade and polyphyletic in the other clade |
| C4 | PPdK-related protein (PPdK-RP1) | AT4G21210 | 25 | 300 | 1152 | 1006 | C4 | 81 | C4 | 98 | two copies of Sd198, not monophyletic |
| C4 | PPdK-related protein (PPdK-RP2) | AT3G01200 | 25 | 300 | 1155 | 1007 | C3 | 45 | both | 100 | one copy of Saldiv198 sister to Kali collina, otherwise C4 clade separate of Sd agg |
| C4 | Pyrophosphatase (PPase6) | AT5G09650 | 22 | 366 | 897 | 769 | C3 | 70 | C3 | 77 | one copy per individual in Sd agg |
| C4 | Pyruvate orthophosphat dikinase (PPdK) | AT4G15530 | 22 | 477 | 2865 | 2125 | C3; C4 | 85;72 | | | several gene copies of Sd |

| | | | | | | | | | | | |
|---|---|---|---|---|---|---|---|---|---|---|---|
| | | | | | | | | | | | agg; one copy sister to C3 othe copies monophyletic and sister to C4 |
| C4 | Sodium:hydrogen antiporter (NHD2) | AT1G49810 | 18 | 867 | 1731 | 1574 | C4 | 89 | C4 | 99 | |
| C4 | Triose phosphate translocator (TPT) | AT5G46110 | 24 | 405 | 1224 | 977 | C4 | 8 | | | no support;several copies; C4 not monophyletic; Salmont in big C4 clade |
| C4TF | BEL1-like homeo domain 7 | AT2G16400 | 19 | 654 | 1794 | 1347 | C4 | 17 | C4 | 57 | in clade with C4 |
| C4TF | shortroot | AT4G37650 | 23 | 477 | 1656 | 1305 | C4 | 86 | | | Salmont as outgroup to clade with Sd agg and C4 |
| Glyoxylate cycle | Isocitrate lyase | AT3G21720 | 7 | 330 | 1731 | 1002 | no Sd agg | | | | no Sd agg, no Salmont |
| Glyoxylate cycle | Malate synthase | AT5G03860 | 18 | 462 | 1707 | 1482 | | | C4 | 58 | C4 clade not monophyletic, low support within clade including Sd agg |
| Glyoxylate cycle | PEP carboxykinase (PEP-CK1) | AT4G37870 | 23 | 417 | 2001 | 1328 | C3 | 86 | C4 | 97 | one copy of Sv sister to Salmont, other copies of Sd agg in clade with C4 |
| Glyoxylate cycle | PEP carboxykinase (PEP-CK2) | AT5G65690 | 24 | 411 | 2001 | 1282 | | | both | 98 | several copies of sd agg, separated in clades with C3 or C4 |

| | | | | | | | | | | | |
|---|---|---|---|---|---|---|---|---|---|---|---|
| Photorespiration | 2-phosphoglycolate phosphatase (PGP1/PGLP1) | AT5G36700 | 19 | 663 | 1107 | 1004 | C4 | 83 | | | several copies of Salvert; C4 clade not monophyletic |
| Photorespiration | Glutamate:glyoxylate aminotransferase (GGT1) | AT1G23310 | 18 | 315 | 1443 | 1223 | C3 | 35 | C3 | 98 | |
| Photorespiration | Glycerate 3-kinase (GLYK) | AT1G80380 | 20 | 483 | 1254 | 1001 | | | C3 | 74 | |
| Photorespiration | Glycine cleavage H-protein (GDC H-protein) | AT2G35120 | 22 | 354 | 462 | 447 | | | C4 | 71 | general low support and polytomies |
| Photorespiration | Glycine cleavage H-protein (GDC H-protein) | AT1G32470 | 18 | 300 | 492 | 459 | both | 57;24 | both | 72 | several copies of Sd and Sv |
| Photorespiration | Glycine decarboxylase L-protein (GDC L-protein) | AT1G48030 | 24 | 360 | 1512 | 1118 | C3 | 64 | C3 | 95 | Sd agg monophyletic |
| Photorespiration | Glycine decarboxylase P-protein (GDC P-protein) | AT2G26080 | 22 | 453 | 3126 | 2370 | C4 | 58 | C4 | 99 | |
| Photorespiration | Glycine decarboxylase T-protein (GDC T-protein) | AT1G11860 | 26 | 387 | 1221 | 971 | | | C4 | 22 | whole tree low bs, several copies per species |
| Photorespiration | Glycolate oxidase (GOX1) | AT3G14420 | 49 | 279 | 1110 | 757 | C4 | 100;17 | both | 73 | several copies, one clade is missing Salmont; one clade with polytomy |
| Photorespiration | Glycolate oxidase (GOX2) | AT3G14415 | 49 | 300 | 1110 | 756 | C4 | 99;100 | | | several copies in two of three clades, one of which is missing Salmont |
| Photorespiration | Glycolate oxidase (GOX3) | AT4G18360 | 48 | 300 | 1110 | 766 | C4 | 98 | both | 98 | three clades, one without Sd |

| | | | | | | | | | | | |
|---|---|---|---|---|---|---|---|---|---|---|---|
| | | | | | | | | | | | agg, one without Salmont, one with polytomy |
| Photorespiration | NADH-dependent hydroxypyruvate reductase (HPR1) | AT1G68010 | 18 | 465 | 1158 | 1033 | | | C4 | 79 | in clade with all C4 Salsoleae |
| Photorespiration | Serine hydroxymethyltransferase (SHMT1) | AT4G37930 | 47 | 414 | 1545 | 1104 | C4 | 36 | both | 51 | several gene copies, Sd agg not monophyletic, low support |
| Photorespiration | Serine:glyoxylate aminotransferase (AGT/SGT) | AT3G08860 | 27 | 426 | 1431 | 1190 | C4 | 78 | | | |
| Photorespiration | Serine:glyoxylate aminotransferase (AGT1/SGT1) | AT2G13360 | 21 | 327 | 1203 | 1031 | C3 | 71 | both | 82 | two copies of Saldiv and Salvert |
| Photorespiration | Serine:glyoxylate aminotransferase (AGT2/SGT2) | AT4G39660 | 20 | 600 | 1440 | 1304 | C3 | 57 | C3 | 72 | C4 clade not monophyletic |
| Photorespiration | Serine:glyoxylate aminotransferase (AGT3/SGT3) | AT2G38400 | 28 | 426 | 1431 | 1180 | C4 | 77 | C4 | 17 | low support on important nodes |
| Photosynthesis | RuBisCO small subunit 1A | AT1G67090 | 47 | 282 | 546 | 467 | no Sd agg | | | | no Sd agg; low support in general |
| Photosynthesis | RuBisCO small subunit 3b | AT5G38410 | 47 | 282 | 546 | 467 | no Sd agg | | | | no Sd agg |

**Material & Methods S1**

**$CO_2$ compensation point measurements**

We used the portable gas exchange measurement system GFS-3000 (Heinz Walz GmbH, Germany) with the standard measuring head 3010-S equipped with a standard leaf area cuvette and an LED-Array. $CO_2$ zero point and span calibration were made weekly using a 30 ppm calibration gas and $H_2O$ zero point and span calibration monthly using the LI-610 portable dew point generator (LI-COR Biosciences GmbH, Germany). The measuring system was controlled via an external notebook running the GFS-Win V3.51 software. For measurements, the youngest part of the stem having young and adult leaves was enclosed in the cuvette. Photosynthetic active radiation (PAR) was set to an intensity of 1400 µmol $m^{-2}$ $s^{-1}$, relative humidity (rh) was set to 45%, the temperature inside the cuvette (Tcuv) to 25°C, the flow to 750 µmol $s^{-1}$ and the impeller to maximum speed. For approximately 10 minutes the plant was adapted to an initial $CO_2$ value of 380 ppm. The $CO_2$ response curve was determined from measurements at 380 ppm and 120 ppm to 60 ppm in 20 ppm steps and 60 ppm to 0 ppm in 10 ppm steps, taking a zero point for every concentration. Dark respiration was measured accordingly.

**Carbon isotope measurements - additional notes on statistics**

We compared $\delta^{13}C$ values between control, dry and salt treatments within six populations (six individuals per treatment). The Shapiro-Wilk test was used to check for normality of distribution and the Levene's Test for homogeneity of variance. Excluding outlier 198IIr of population 198, all treatment groups of all populations were normally distributed. All populations had homogeneous variances except population 188. Since not all assumptions for an ANOVA test were met, we chose the Kruskal-Wallace rank sum test to compare treatments within and between populations.

**PEPC activity measurements - Sample preparation**

Approximately 0.2 g leaf material was frozen in liquid nitrogen and subsequently ground in a pre-chilled mortar, using 1 mL extraction buffer containing 50 mM HEPES buffer (pH 8.0), 1% (w/v) polyvinylpyrrolidone (PVP), 1mM ethylenediaminetetraacetic acid (EDTA), 10 mM dithiothreitol (DTT) and 0.1% (v/v) Triton X-100 with 20 µl protease inhibitors. After centrifugation at 10000 x *g* for 1 min, 500 µl of the supernatant was mixed with 7.5 µl 1 M magnesium chloride solution ($MgCl_2$) and 15 µl 500 mM sodium hydrogen carbonate solution ($NaHCO_3$) and stored on ice for 30 min before using it in a coupled enzyme assay at 340 nm. The assay buffer contained 100 mM EPPS buffer (pH 8.0), 20 mM $MgCl_2$, 1 mM EDTA, 0.2 mM NADH, 5 mM glucose-6-phosphate, 1 mM $NaHCO_3$ and 2.4 units malate dehydrogenase.

**Transcriptome processing and nuclear phylogenetic analyses**

To evaluate the potential hybrid origin of the *S. divaricata* agg., we generated gene trees from a transcriptome dataset comprising $C_3$, $C_4$ and $C_2$ species of Salsoleae; in total 13 transcriptomes representing 12 species. Three samples of these were newly sequenced on an Illumina HiSeq2500 platform at the University of Minnesota Genomics Center (paired-end 125 bp). For RNA extraction protocol and library preparation refer to Morales-Briones *et al.* (2021). Additionally, we sampled two species of Camphorosmeae and one of Caroxyloneae, and *Beta vulgaris* L. subf. Betoideae - Amaranthaceae as outgroups. The *Beta vulgaris* genome is the best annotated genome closest to our group of interest. **Table 1** lists all samples with their photosynthesis type, ploidy if known and SRA accession number.

We followed Morales-Briones *et al.* (2021) for raw read processing, transcriptome assembly, transcript clustering, and homolog tree inference. Sequencing errors in raw reads were corrected with Rcorrector (Song and Florea 2015) and reads flagged as uncorrectable were removed. Sequencing adapters and low-quality bases were removed with Trimmomatic v0.36 (ILLUMINACLIP: TruSeq_ADAPTER: 2:30:10 SLIDINGWINDOW: 4:15 LEADING: 5 TRAILING: 5 MINLEN: 25; Bolger et al. 2014). Additionally, chloroplast and mitochondrial reads were filtered with Bowtie2 v 2.3.2 (Langmead and Salzberg 2012) using publicly available Caryophyllales organelle genomes from the Organelle Genome Resources database (RefSeq; [Pruitt et al. 2007]; last accessed on October 17, 2018) as references. Read quality was assessed with FastQC v 0.11.7 (https://www.bioinformatics.babraham.ac.uk/projects/fastqc/). Overrepresented sequences detected with FastQC were discarded. *De novo* assembly was carried out with Trinity v 2.5.1 (Grabherr et al. 2011) with default settings, but without in silico normalization. Assembly quality was assessed with Transrate v 1.0.3 (Smith-Unna et al. 2016). Low quality and poorly supported transcripts were removed using individual cut-off values for three contig score components of Transrate: 1) proportion of nucleotides in a contig that agrees in identity with the aligned read, s(Cnuc) $\leq$ 0.25; 2) proportion of nucleotides in a contig that have one or more mapped reads, s(Ccov) $\leq$0.25; and 3) proportion of reads that map to the contig in correct orientation, s(Cord) ≤ 0.5. Furthermore, chimeric transcripts (*trans*-self and *trans*-multi-gene) were removed following the approach described in Yang and Smith (2013) using *Beta vulgaris* as the reference proteome, and percentage similarity and length cutoffs of 30 and 100, respectively. In order to remove isoforms and assembly artifacts, filtered reads were remapped to filtered transcripts with Salmon v 0.9.1 (Patro et al. 2017) and putative genes were clustered with Corset v 1.07 (Davidson and Oshlack 2014) using default settings, except that we used a minimum of five reads as threshold to remove transcripts with low coverage (-m 5). Only the longest transcript of each putative gene inferred by Corset was retained (Chen et al. 2019). Filtered transcripts were translated with TransDecoder v 5.0.2 (Haas et al. 2013) with default settings and the proteome of *Beta vulgaris* and *Arabidopsis thaliana* to identify open reading frames. Finally, coding sequences (CDS) from translated amino acids were further reduced with CD-HIT v 4.7 (-c 0.99; [Fu et al. 2012]) to remove near-identical sequences. Scripts and instructions for read processing, assembly, translation, and homology and orthology search can be found at https://bitbucket.org/yanglab/phylogenomic_dataset_construction/.

Initial homology inference was carried out following Yang and Smith (2014) with some modifications. First, an all-by-all BLASTN search was performed on CDS using an *E* value cutoff of 10 and max_target_seqs set to 100. Raw BLAST output was filtered with a hit fraction of 0.4. Then putative homologs groups were clustered using MCL v 14-137 (van Dongen 2000) with a minimum minus log-transformed *E* value cutoff of 5 and an inflation value of 1.4. Finally, only clusters with a minimum of 25 taxa were retained. Individual clusters were aligned using MAFFT v 7.307 (Katoh and Standley 2013) with settings '–genafpair –maxiterate 1000'. Aligned columns with more than 90% missing data were removed using Phyx (Brown et al. 2017). Homolog trees were built using RAxML v 8.2.11 (Stamatakis 2014) with a GTRCAT model and clade support assessed with 200 rapid bootstrap (BS) replicates. Spurious or outlier long tips were detected and removed with TreeShrink v 1.0.0. by maximally reducing the tree diameter (Mai and Mirarab 2018). Monophyletic and paraphyletic tips that belonged to the same taxon were removed, keeping the tip with the highest number of characters in the trimmed alignment. After visual inspection of ca. 50 homolog trees, we determined that internal branches longer than 0.25 were likely representing deep paralogs. These branches were cut apart, keeping resulting subclades with a minimum of 17 taxa. Homolog tree inference, tip and outlier removal, and deep paralog cutting was carried out for a second time using the same settings to obtain final homologs. Orthology inference was carried out following the 'monophyletic outgroup' approach from Yang and Smith (2014), keeping only ortholog groups with all 17 taxa. The MO approach filters for trees that have outgroup taxa being monophyletic and single-copy, and therefore filters for single- and low-copy genes. This approach roots the gene tree by the outgroups (here *Beta vulgaris*), traverses the rooted tree from root to tip, and removes the side with less taxa when gene duplication is detected. If no taxon duplication is detected in a homolog tree, the MO approach outputs a one-to-one ortholog. Scripts used can be found at https://bitbucket.org/yanglab/phylogenomic_dataset_construction/src/master/.

We used concatenation and coalescent-based methods for phylogenetic reconstruction. Individual orthologs were aligned using MAFFT v 7.307 (--genafpair –maxiterate 1000; Katoh & Standley, 2013) and trimmed by a minimal column occupancy of 0.3 using phyx (Brown *et al*., 2017). For the concatenation approach, we prepared a supermatrix by keeping only ortholog alignments with at least 300 bp and all 17 taxa. We estimated a maximum likelihood (ML) tree with RAxML v 8.2.11 (Stamatakis, 2014) using a partition by gene scheme with a GTRGAMMA model for each partition. Clade support was assessed with 100 rapid bootstrap (BS) replicates. To estimate a species tree that is statistically consistent with the multi-species coalescent (MSC), we first inferred individual ML gene trees using RAxML with a GTRGAMMA model, and 100 BS replicates to assess clade support. Individual gene trees were then used to estimate a species tree using ASTRAL-III v.5.6.3 (Zhang *et al*., 2018) using local posterior probabilities (LPP; Sayyari & Mirarab, 2016) to assess clade support. To examine nuclear gene tree discordance, we first calculated the internode certainty all (ICA) value to quantify the degree of conflict on each node of the species tree given individual gene trees (Salichos *et al*., 2014). Also, we calculated the number of conflicting and concordant bipartitions on each node of the species trees. We calculated both the ICA scores and the number of conflicting and concordant bipartitions with Phyparts (Smith *et al*., 2015) using individual gene trees with BS support cutoff 50% for each node, and both the ASTRAL and RAxML topologies as mapping trees. To visualize the proportions of conflicting and concordant bipartitions we used the script phypartspiecharts.py available at https://github.com/mossmatters/MJPythonNotebooks.

Additionally, to distinguish strong conflict from weakly supported branches, we evaluated tree conflict and branch support with Quartet Sampling (QS; Pease *et al*., 2018) using 1,000 replicates. Quartet Sampling takes a topology (here a species tree) and an alignment to subsample quartets and assesses the confidence, consistency, and informativeness of each internal branch by the relative frequency of the three possible quartet topologies at each node (Pease *et al*., 2018).

**Assessment of hybridization**

To detect possible hybridization, first we inferred species networks under a maximum pseudo-likelihood (Yu & Nakhleh, 2015) approach using PhyloNet v.3.6.9. (Than *et al*., 2008) with the command "InferNetworks_MPL" and using the individual ML gene trees as input. Network searches were performed, allowing for up to five hybridization events and optimizing the branch lengths and inheritance probabilities of the returned species networks under the full likelihood. To estimate the best number of hybridizations and test whether the species network fits our gene trees better than a strictly bifurcating tree, we computed the likelihood scores of the nuclear and plastid trees, given the individual gene trees, as implemented in Yu *et al*. (2012), using the command 'CalGTProb'. Finally, we performed model selection using the bias-corrected Akaike information criterion (Sugiura, 1978) and the Bayesian information criterion (Schwarz, 1978). The number of parameters equals the number of branch lengths, plus the number of inheritance probabilities. The number of gene trees was used to correct for finite sample size. We also tested for hybridization with HyDe (Blischak *et al*., 2018) which uses site pattern frequencies (Kubatko & Chifman, 2019) to estimate the amount of admixture (γ) between two parental lineages that form a third hybrid lineage. We tested all triples combinations in all directions using the script 'run_hyde.py, *Beta vulgaris* as outgroup, the nuclear concatenated alignment, and a mapping file to assign all individuals as distinct 'taxa' (only S. *divaricata*, had two sampled individuals). Test significance was assessed with a Bonferroni correction ($\alpha$= 0.05) for the number of hypothesis tests conducted with estimates of γ between 0 and 1 (Blischak *et al*., 2018).

**Plastome assembly and phylogenetic analysis**

To investigate phylogenetic signal from plastid sequences, *de novo* assemblies were carried out with the Fast-Plast v.1.2.6 pipeline (https://github.com/mrmckain/Fast-Plast) using the filtered organelle reads obtained from the transcriptome raw read processing. No complete plastome could be assembled, so filtered contigs produced by Spades v 3.9.0 (Bankevich *et al*., 2012) were mapped to *Haloxylon persicum* (GenBank NC_027669), with one copy of the Inverted Repeat removed, and manually edited with Geneious v.11.1.5 (Kearse *et al*., 2012) to produce the final oriented contigs. Contiguous plastome contigs, including the reference plastome of *Beta vulgaris* (GenBank KR230391), were aligned with MAFFT with the setting

‘--auto’ and ambiguous alignment sites were removed using GBLOCKS v0.9b (Castresana, 2000; -t=d -b1=9 -b2=9 -b3=10 -b4=10 -b5=a). A ML tree was inferred with IQ-TREE v.1.6.1 (Nguyen *et al.*, 2015) using the automated model selection (Kalyaanamoorthy *et al.*, 2017) and 200 standard non-parametric BS replicates for branch support. Additionally, we used QS with 1,000 replicates, to detect potential plastome conflict in the backbone as seen in other groups of Amaranthaceae s.l. (Morales-Briones *et al.*, 2021).

**Analysis of photosynthetic gene trees**

We reconstructed gene trees for 50 genes that are important in photosynthesis to determine if the genes in *Salsola divaricata* agg. show a tendency to group with either a $C_3$ or a $C_4$ species. We conducted a baited homology inference and topology analysis of such key photosynthesis genes using the pre-filtered coding and protein sequences produced during the transcriptome processing. We downloaded coding and protein sequences from *Arabidopsis thaliana* as baits from the TAIR website (https://www.arabidopsis.org/tools/bulk/sequences/index.jsp). These loci were determined based on Lauterbach *et al.* (2017a,b) and included two RuBisCO subunit genes, four genes coding for proteins of the glyoxylate cycle, 17 genes encoding photorespiratory proteins, and 25 genes coding for $C_4$-associated proteins (Lauterbach *et al.*, 2017b) as well as two $C_4$-associated transcription factors (Lauterbach *et al.*, 2017a). The scripts used can be found at https://bitbucket.org/yangya/c2_salsola and the TAIR locus IDs with gene annotation and pathway assignment can be found in **Supporting Information Table S3**. After manually checking the alignments and gene trees, we listed if the *S. divaricata* agg. was either in a sister relationship with $C_3$ species *S. montana* or the $C_4$ clade. When there were several gene copies or weak bootstrap support, we additionally noted whether our group of interest was in a clade with the $C_3$ or the $C_4$ species.

**References Material & Methods S1**


**Bankevich A, Nurk S, Antipov D, Gurevich AA, Dvorkin M, Kulikov AS, Lesin VM, Nikolenko SI, Pham S, Prjibelski AD, *et al.* 2012**. SPAdes: A New Genome Assembly Algorithm and Its Applications to Single-Cell Sequencing. *Journal of Computational Biology* **19**: 455–477. doi:10.1089/cmb.2012.0021

**Blischak PD, Chifman J, Wolfe AD, Kubatko LS**. **2018**. HyDe: A Python Package for Genome-Scale Hybridization Detection (D Posada, Ed.). *Systematic Biology* **67**: 821–829. doi:10.1093/sysbio/syy023

**Brown JW, Walker JF, Smith SA**. **2017**. Phyx: phylogenetic tools for unix. *Bioinformatics* **33**: 1886–1888. doi:10.1093/bioinformatics/btx063.

**Castresana J**. **2000**. Selection of Conserved Blocks from Multiple Alignments for Their Use in Phylogenetic Analysis. *Molecular Biology and Evolution* **17**: 540–552. doi:10.1093/oxfordjournals.molbev.a026334.

**Kalyaanamoorthy S, Minh BQ, Wong TKF, von Haeseler A, Jermiin LS**. **2017**. ModelFinder: fast model selection for accurate phylogenetic estimates. *Nature Methods* **14**: 587–589. doi:10.1038/nmeth.4285.

**Katoh K, Standley DM**. **2013**. MAFFT Multiple Sequence Alignment Software Version 7: Improvements in Performance and Usability. *Molecular Biology and Evolution* **30**: 772–780. doi:10.1093/molbev/mst010.

**Kearse M, Moir R, Wilson A, Stones-Havas S, Cheung M, Sturrock S, Buxton S, Cooper A, Markowitz S, Duran C, *et al.* 2012**. Geneious Basic: An integrated and extendable desktop software platform for the organization and analysis of sequence data. *Bioinformatics* **28**: 1647–1649. doi:10.1093/bioinformatics/bts199.

**Kubatko LS, Chifman J**. **2019**. An invariants-based method for efficient identification of hybrid species from large-scale genomic data. *BMC Evolutionary Biology* **19**: 112. doi:10.1186/s12862-019-1439-7.

**Lauterbach M, Billakurthi K, Hankeln TM, Westhoff P, Gowik U, Kadereit G. 2017a.** Organ-specific gene expression profiling of cotyledons and leaves of $C_4$ *Salsola oppositifolia* and insights into regulation of $C_2$ and $C_4$ photosynthesis in Salsoleae. In: Lauterbach M. *$C_4$ photosynthesis in Salsoleae: Ontogenetic gene expression profiling of closely related $C_3$, $C_3$-$C_4$ and $C_4$ species using RNA-Seq*. PhD thesis, Mainz, Germany.

**Lauterbach M, Billakurthi K, Kadereit G, Ludwig M, Westhoff P, Gowik U**. **2017**. $C_3$ cotyledons are followed by $C_4$ leaves: Intra-individual transcriptome analysis of *Salsola* *soda* (Chenopodiaceae). *Journal of Experimental Botany* **68**: 161–176.

**Morales-Briones DF, Kadereit G, Tefarikis DT, Moore MJ, Smith SA, Brockington SF, Timoneda A, Yim WC, Cushman JC, Yang Y**. **2021**. Disentangling Sources of Gene Tree Discordance in Phylogenomic Datasets: Testing Ancient Hybridizations in Amaranthaceae s.l. *Systematic Biology* **70**: 219–235. doi:10.1093/sysbio/syaa066.

**Nguyen L-T, Schmidt HA, von Haeseler A, Minh BQ**. **2015**. IQ-TREE: A Fast and Effective Stochastic Algorithm for Estimating Maximum-Likelihood Phylogenies. *Molecular Biology and Evolution* **32**: 268–274. doi:10.1093/molbev/msu300.

**Pease JB, Brown JW, Walker JF, Hinchliff CE, Smith SA**. **2018**. Quartet Sampling distinguishes lack of support from conflicting support in the green plant tree of life. *American Journal of Botany* **105**: 385–403. doi:10.1002/ajb2.1016.

**Salichos L, Stamatakis A, Rokas A**. **2014**. Novel Information Theory-Based Measures for Quantifying Incongruence among Phylogenetic Trees. *Molecular Biology and Evolution* **31**: 1261–1271. doi:10.1093/molbev/msu061.

**Sayyari E, Mirarab S**. **2016**. Fast Coalescent-Based Computation of Local Branch Support from Quartet Frequencies. *Molecular Biology and Evolution* **33**: 1654–1668. doi:10.1093/molbev/msw079.

**Smith SA, Moore MJ, Brown JW, Yang Y**. **2015**. Analysis of phylogenomic datasets reveals conflict, concordance, and gene duplications with examples from animals and plants. *BMC Evolutionary Biology* **15**: 150. doi:10.1186/s12862-015-0423-0.

**Stamatakis A**. **2014**. RAxML version 8: a tool for phylogenetic analysis and post-analysis of large phylogenies. *Bioinformatics* **30**: 1312–1313. doi:10.1093/bioinformatics/btu033.

**Schwarz G**. **1978**. Estimating the Dimension of a Model. *The Annals of Statistics* **6**: 461–464. doi:10.1214/aos/1176344136.

**Sugiura N**. **1978**. Further analysts of the data by akaike's information criterion and the finite corrections. *Communications in Statistics - Theory and Methods* **7**: 13–26. doi:10.1080/03610927808827599.

**Than C, Ruths D, Nakhleh L**. **2008**. PhyloNet: A software package for analyzing and reconstructing reticulate evolutionary relationships. *BMC Bioinformatics* **9**: 1–16. doi:10.1186/1471-2105-9-322.

**Yang Y, Smith SA**. **2014**. Orthology inference in nonmodel organisms using transcriptomes and low-coverage genomes: Improving accuracy and matrix occupancy for phylogenomics. *Molecular Biology and Evolution* **31**: 3081–3092.

**Yu Y, Degnan JH, Nakhleh L**. **2012**. The probability of a gene tree topology within a phylogenetic network with applications to hybridization detection. *PLoS Genetics* **8**. doi:10.1371/journal.pgen.1002660.

**Yu Y, Nakhleh L**. **2015**. A maximum pseudo-likelihood approach for phylogenetic networks. *BMC Genomics* **16**: S10. doi:10.1186/1471-2164-16-S10-S10.

**Zhang C, Rabiee M, Sayyari E, Mirarab S. 2018.** ASTRAL-III: polynomial time species tree reconstruction from partially resolved gene trees. *BMC bioinformatics* **19**: 15–30. doi:10.1186/s12859-018-2129-y
